# Designing Molecular RNA Switches with Restricted Boltzmann Machines

**DOI:** 10.1101/2023.05.10.540155

**Authors:** Jorge Fernandez-de-Cossio-Diaz, Pierre Hardouin, Francois-Xavier Lyonnet du Moutier, Andrea Di Gioacchino, Bertrand Marchand, Yann Ponty, Bruno Sargueil, Rémi Monasson, Simona Cocco

**Affiliations:** CNRS UMR 8023, Laboratory of Physics of the Ecole Normale Supérieure & PSL Research, Sorbonne Université, 24 rue Lhomond, 75005 Paris, France; Université Paris-Saclay, CNRS, CEA, Institut de Physique Théorique, 91191, Gif-sur-Yvette, France; CNRS UMR 8038, CitCoM, Université de Paris, 4 avenue de l’observatoire, 75006 Paris, France; CNRS UMR 7161, LIX, Ecole Polytechnique, Institut Polytechnique de Paris, 1 rue Estienne d’Orves, 91120 Palaiseau, France

**Author notes:** Equal contribution (Jorge Fernandez-de-Cossio-Diaz, Pierre Hardouin).

## Abstract

Riboswitches are structured allosteric RNA molecules that change conformation upon metabolite binding, triggering a regulatory response. Here we focus on the *de novo* design of riboswitch-like aptamers, the core part of the riboswitch undergoing structural changes. We use Restricted Boltzmann machines (RBM) to learn generative models from homologous sequence data. We first verify, on four different riboswitch families, that RBM-generated sequences correctly capture the conservation, covariation and diversity of natural aptamers. The RBM model is then used to design new SAM-I riboswitch aptamers. To experimentally validate the properties of the structural switch in designed molecules, we resort to chemical probing (SHAPE and DMS), and develop a tailored analysis pipeline adequate for high-throughput tests of diverse sequences. We probe a total of 476 RBM-designed and 201 natural sequences. Designed molecules with high RBM scores, with 20% to 40% divergence from any natural sequence, display *≈* 30% success rate of responding to SAM with a structural switch similar to their natural counterparts. We show how the capability of the designed molecules to switch conformation is connected to fine energetic features of their structural components.

## INTRODUCTION

Riboswitches are regulatory RNA elements found mostly in bacterial and in some eukaryotic messenger RNAs. Usually located upstream of coding sequences, they modulate the expression of the downstream gene at the transcriptional or translation level in the presence of a specific metabolite [48, 66, 75, 77]; some riboswitches placed within genes even regulate alternative splicing [42]. In order to perform their function, these RNA motifs switch between two stable conformations in response to binding of their cognate metabolite to the aptamer domain of the riboswitch (Figure 1). This change of conformation, in turn, affects the expression platform, where the regulation signals are located. Understanding how the aptamer domain by itself is able to implement a structural switch in response to the ligand, and how this is encoded in the sequence, is an important step towards the characterization of the full riboswitch regulation.

**FIG. 1.**
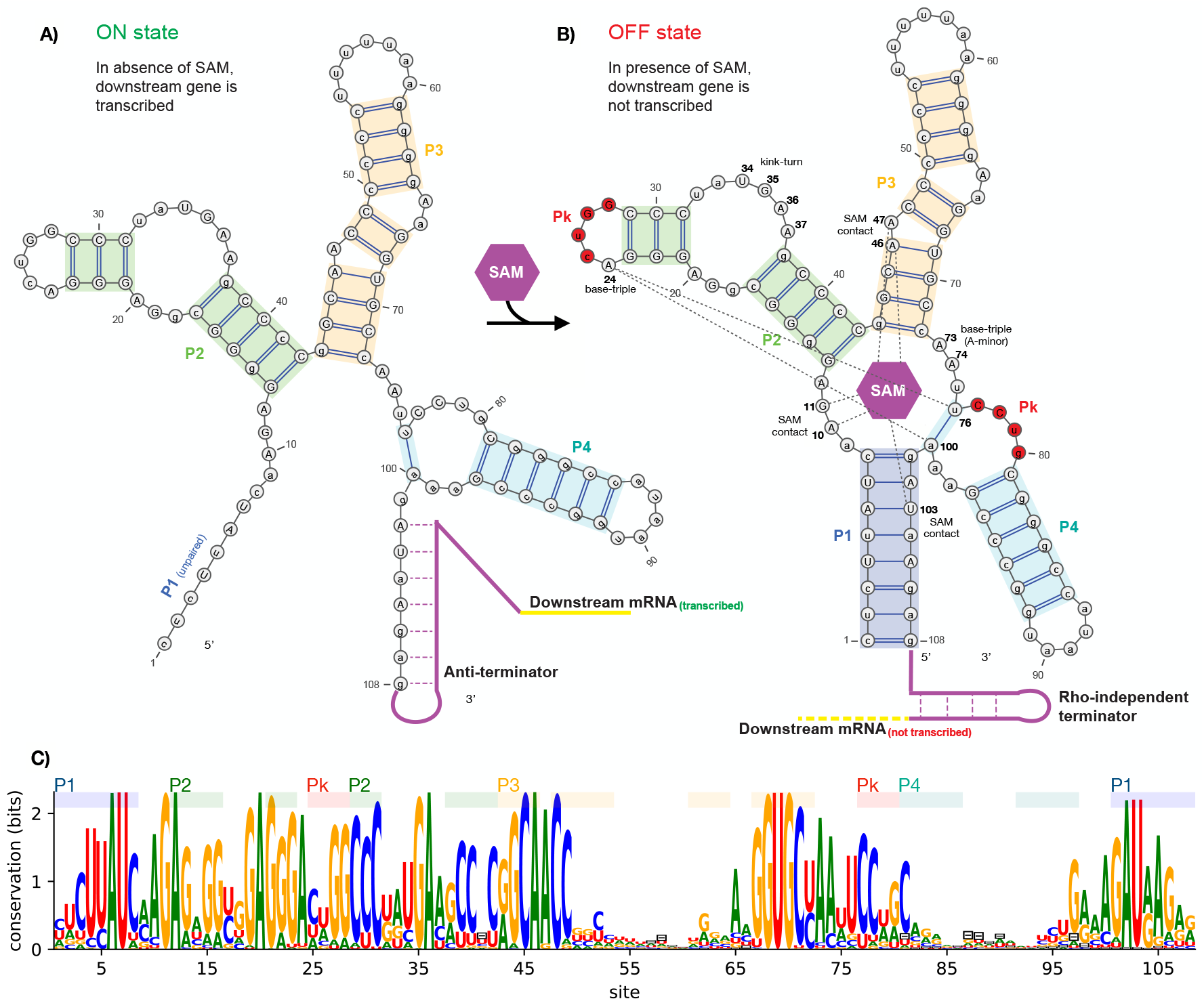
Structure, regulatory function, and sequence conservation of the aptamer domain of the SAM-I riboswitch, acting at a transcriptional level. **A)** In absence of SAM, the P1 helix of the aptamer domain is unpaired, leaving the 3’-end free to pair with the anti-terminator segment of the expression platform. This conformation is incompatible with the terminator motif, resulting in transcription of the downstream gene (ON state). **B)** SAM (represented by the purple hexagon) is captured in a groove contacting several sites around the central four-way junction. In the bound-state conformation, the P1 helix is fully base-paired. The expression platform is then free to form a Rho-independent terminator hairpin, which stops transcription of the nascent RNA, thus blocking the expression of a downstream gene (OFF state). The figure also shows several structural elements of the consensus secondary structure of the aptamer domain, including helices P1, P2, P3, P4, and a pseudoknot (Pk) in red. Other sites of interest participating in tertiary contacts (dashed lines) in response to SAM are highlighted in bold, including SAM contacts and base-triples. Secondary structure plots are obtained with VARNA [17]. **C)** Sequence conservation logo of aligned homologs of the SAM-I riboswitch aptamer domain family (RF00162 on Rfam). Gaps are indicated by the character ‘⊟’.

The sequence-to-function mapping of structured RNAs is a complex problem. In the course of evolution, sequence patterns necessary for function are conserved, suggesting that large sequence datasets can shed light on this mapping. Comparative analysis of homologous RNA sequences collected in Multiple Sequence Alignments (MSA) [56] have been successful to predict secondary RNA structures, tertiary structural motifs, and even the entire three dimensional architecture of complex RNA [11, 23, 28, 51, 64, 79]. Covariation analysis has also been used to predict pseudoknots and other tertiary contacts from statistical couplings inferred from conservation and covariation across the MSA columns [19, 92], or by including positive and negative evolutionary information such as in the Cascade covariation Folding Algorithm (CacoFold) [63]. Machine learning approaches have recently shown promising results in RNA structure prediction. Among them Rosetta FARFAR2 [89] uses Monte-Carlo-based fragment assembly methods and can be aided by geometric deep learning approaches such as ARES [84] to score putative structures. DeepFoldRNA [59] significantly outperformed the state-of-the-art tertiary structure prediction from sequence only. Although these approaches look promising, AlphaFold-level accuracies [39] (for proteins) are not yet reached in RNA structure prediction [61, 81].

The mirroring problem of designing RNA sequences capable of folding in a particular target structure or of performing a desired function has also long been investigated. One successful approach is based on directed evolution (SELEX). RNA sequences are selected from an initial random library to optimize a target function, such as the switching dynamics for bistable aptamers [49]. Models trained on such data are capable of classifying sequences according to their functionality and of extracting key sequence-features for the desired function [1, 21, 24, 26, 37]. Classifiers have been used downstream of random mutagenesis to filter out good sequences, but this approach only works if the libraries already contain good candidates. In parallel, much effort has been devoted to the rational design of secondary structures, in particular with minimum free energy approaches [25, 93]. However, due to algorithmic complexity [64, 88], those approaches often ignore pseudoknots and other tertiary contacts known to be essential for the function of some RNAs, such as riboswitches or ribozymes.

To date, building generative models effective in designing RNA sequences with tertiary structural targets remains a challenging problem. From this point of view, riboswitches, in addition to their fundamental interest in biology and relevance for the RNA world hypothesis [42], offer a difficult design problem, as their sequences encode not only two conformational structures but also a metabolite-mediated switching mechanism between them. In the present work, we address this challenging issue and show how to design functional RNA switches (albeit devoid of expression platform) from natural sequence data.

One of the largest identified groups of riboswitches recognize S-adenosyl-methionine (SAM) as their effector metabolite [27, 60]. While six different SAM binding structural motifs have been identified, this study focuses on those harbouring type I SAM aptamers (SAM-I) [3]. Figure 1A shows the secondary structure of the aptamer domain in absence of SAM, where transcription is allowed (ON state), while panel B depicts the structure when SAM is bound and transcription continuation is prevented (OFF state). Upon SAM binding, the aptamer cooperatively folds into the closed structure characterised by the stabilisation of P1, three triple base pairs and a pseudoknot (red in the figure) [67]. The closed state of the aptamer is stabilized by direct tertiary contacts between SAM and specific nucleotides forming the SAM binding pocket [53, 60].

Hereafter we employ Restricted Boltzmann machines (RBM), a two-layer generative neural network to design new SAM aptamers (Fig. 2A). RBMs have recently been shown to provide interpretable models of proteins in various contexts [7, 8, 50, 86], with application to design [21, 47]. By learning the sequence statistics of the SAM-I riboswitch family, the RBM models the constraints that enable aptamers to adopt the correct secondary structure, form tertiary contacts and effect a conformational switch in response to SAM presence.

**FIG. 2.**
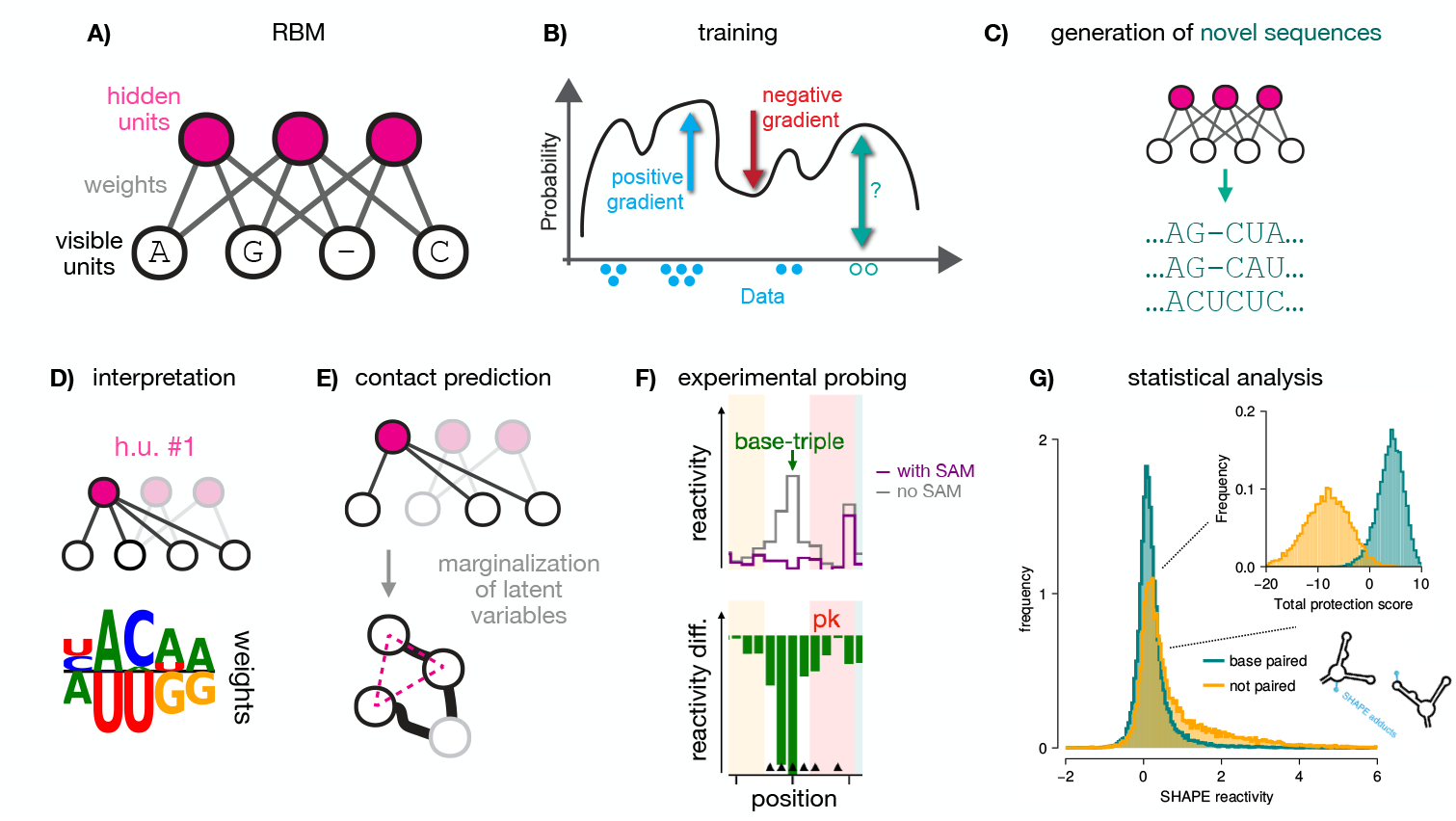
RNA generative modeling with RBM and experimental validation. **A)** A Restricted Boltzmann machines (RBM), with the visible layer carrying nucleotides A, C, G, U, or – (alignment gap symbol), and a hidden layer extracting features. The two layers are connected by *weights*. **B)** The RBM is trained by maximization of a regularized likelihood, see Eq. (S4). A gradient term increases the probability of regions in sequence space populated by data, automatically discovering features desirable for functional sequences (blue), while an opposite gradient term lowers the probability of regions void of data (red). The RBM may also assign large probability to potentially interesting sequences not covered by data (teal). **C)** The model can be sampled to generate novel sequences that may significantly differ from the natural ones (teal). **D)** Hidden units extract latent features (nucleic-acid motifs) through the weights. Weight values, either positive or negative, are shown above or below the zero-weight horizontal bar in the logo plots, see Methods. Combining these motifs together allows RBM to design functional RNA sequences. **E)** The RBM is able to model complex interactions along the RNA sequence. Here, a hidden unit interacting with three visible units is highlighted. After marginalizing over hidden-unit configurations, effective interactions arise between the visible sites, see Eq. (5). Here we represent schematically a three-body interaction, arising from the three weights onto the marginalized hidden unit. **F)** Designed sequences are tested experimentally with chemical probing approaches. Reactivities of sites to the probes may differ when SAM is absent or present (top); difference in reactivities between the two conditions is informative about structural changes (bottom). **G)** Distributions of reactivities obtained with SHAPE-MaP slightly differ for paired and unpaired nucleotides. Statistical resolution of global structural changes triggered by SAM can then be enhanced by aggregating multiple sites. Inset: distributions over 24 sites, see Methods, Section I and Supplementary Figures S25, S39.

The RBM model was used to design 476 sequences, which we experimentally tested with SHAPE-MaP and DMS, two chemical probing methods giving information about paired and unpaired residues in the structures. Comparison of the reactivity profiles in the presence or absence of SAM allows us to assess the effectiveness of the structural switch for each tested molecule. This highthroughput analysis is made possible by the introduction of an automated Bayesian analysis of the SHAPE and DMS reactivity profiles. Our results for RBM-generated aptamers are compared to experiments on 201 natural sequences, and 58 sequences designed by RFAM Covariance Models, another generative model capturing local conservation and secondary-structure covariation only.

## RESULTS

Our pipeline is described in Fig. 2 and includes: sequence data acquisition from Rfam [41], training and sampling the RBM to design artificial SAM-I aptamers, experimental characterization of SAM-induced conformational switch in natural and designed sequences by chemical probing (SHAPE [18, 73] and DMS [52]), and statistical analysis of the measured reactivities.

### A. Generative models of SAM riboswitch aptamers

We train an RBM (Figs. 2A,B) on a multiple sequence alignment (MSA) of natural homologues of the aptamer domain of SAM-I riboswitches, gathered from the Rfam [41] database (Rfam ID: RF00162). RBM are energybased generative models, that once trained, define a score, −*E*_eff_(**v**), over all possible sequences **v**. Sequences with high scores (equivalently, low energies) are then “good” fits to the family, according to the model. Artificial sequences of high score can be generated by sampling the resulting Boltzmann measure, 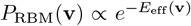, see Fig. 2C and Methods for details.

The weights between visible units, carrying the RNA sequence, and hidden units, extract latent factors of variation in the data, Fig. 2D). After marginalization over those latent variables, effective interactions between pairs of residues can be computed [86], defining epistatic scores between sites (Fig. 2E and Supplementary Eq. (S12) for precise definition). Pairs of sites with large epistatic scores correspond to major secondary and tertiary contacts in folded aptamers, see heatmap in Fig. 3A. Interestingly, epistatic scores at P1 are weaker than in other helices, reflecting the flexibility of P1, which is able to open or close in concert with SAM binding (Fig. 1). The pseudoknot is also correctly identified (red in Fig. 3A), proving the capability of RBM to identify tertiary motifs. Besides structural contacts, the RBM hidden units capture extended motifs, most likely relevant for tertiary structure formation and SAM binding, see weights in Figs. 3B,C.

**FIG. 3.**
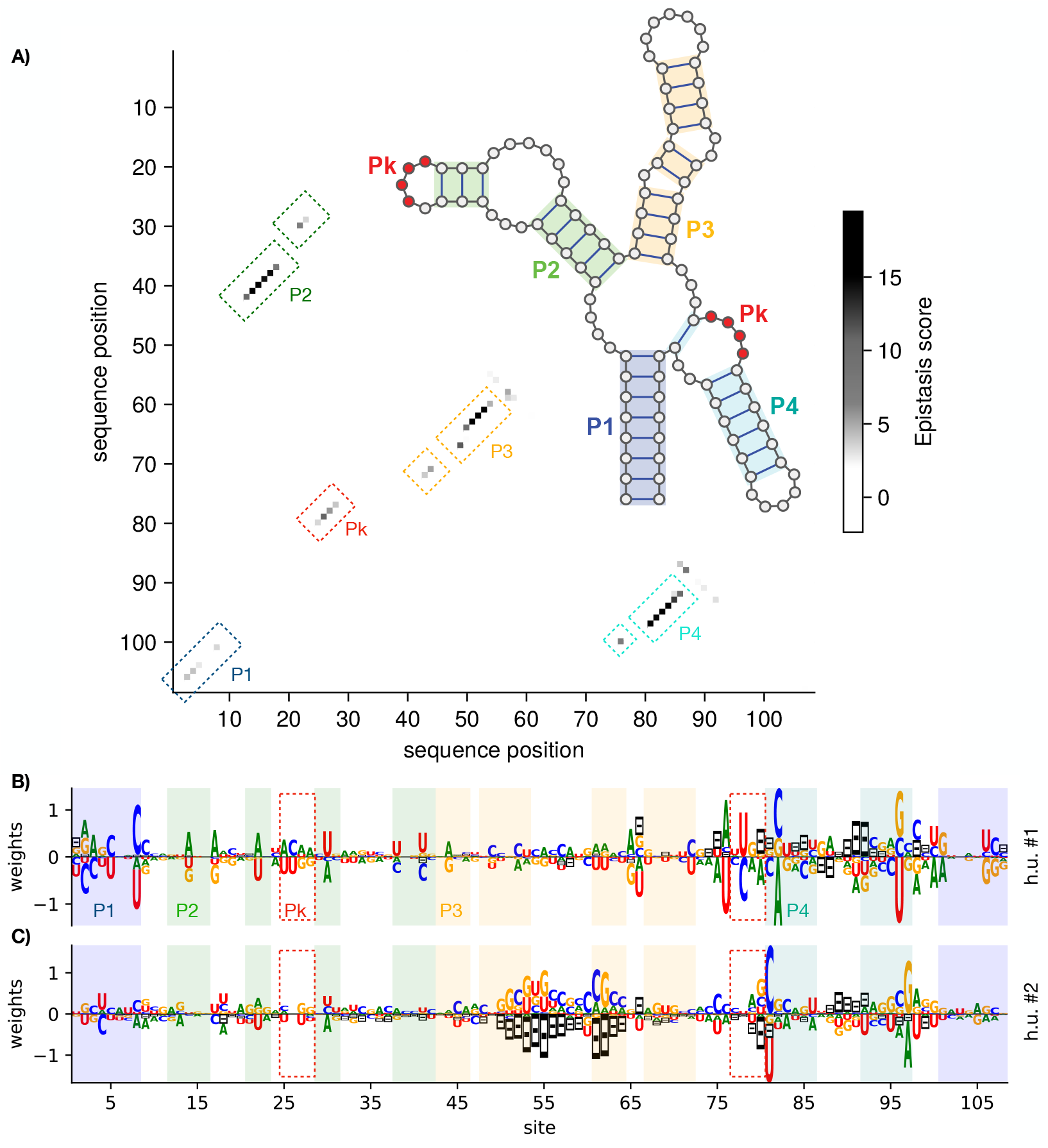
Interpretation of RBM extracted features. **A)** Contact map based on the epistatic scores for nucleotide pairs computed with the RBM [86]. The highest epistatic scores correspond to major secondary and tertiary contacts of the SAM-bound aptamer structure, shown in the inset. **B**,**C)** Sequence logos of the weights *w*_*iµ*_(*v*_*i*_) attached to exemplary hidden units (#1 and #2) of the RBM, selected by having the highest weight norms. Each letter size in the logo is proportional to the corresponding weight, see Figure 2D and [82, 86]). Sites are colored according to the secondary structure element they belong to, including the paired (P) helices P1 (light purple), P2 (green), P3 (yellow), and P4 (teal). Sites participating in the pseudoknot (Pk) are also highlighted (red dashed box). In hidden unit #1, Watson-Crick complementarity along P1 (*e*.*g*., site 8 with 101) is favored, in agreement with base pairing of these positions at the 5’ and 3’ ends of the P1 helix. The same unit also puts weights on complementarity along the pseudoknot (*e*.*g*. sites 25-28 with 77-80), helping stabilize this tertiary contact. The fact that these complementarity constraints, belonging to different structural motifs, are enforced by the same unit, suggests that P1 and the pseudoknot stabilize in a concerted manner (*c*.*f*. Fig. 1) in response to SAM. Hidden unit #2, on the other hand, places significant weight in the complementarity between sites 81 and 97, stabilizing P4 and along various P3 sites, favouring a dichotomy between stabilizing complementarity or deletions in this segment. Indeed, some natural sequences lack a hairpin loop at P3 (sites 50–64), consistently with a negative activation of h.u. #2.

We then evaluate the sequences designed by the RBM by comparing their scores to the ones of natural sequences and sequences designed by Covariance Models (CM). CM capture the conservation of residues along the sequence, as well as correlations due to the complementarity of base pairs in the secondary structure [22], but are unable to model tertiary motifs (such as pseudoknots). As Rfam sequence alignments [40] are based on CM [56], our first baseline model for RF00162 was directly downloaded from Rfam (Methods), and will be referred to as Rfam CM (rCM) in the following.

In Fig. 4A, we show a scatter plot of rCM vs RBM scores for natural, RBM- and CM-generated sequences. RBM-generated sequences have rCM scores comparable to the natural ones, indicating that RBM samples satisfy the constraints imposed by the rCM model to the same extent as natural sequences. Moreover, RBM samples have RBM scores comparable to natural sequences, while rCM samples have significantly smaller scores, suggesting that the RBM impose further constraints beyond those captured by rCM, such as tertiary contacts (*e*.*g*. pseudo-knot), which could be important for the aptamer function. We also check that R-scape [64] supports significant covariation across pseudoknot sites for RBM samples, contrary to rCM samples as expected (see Supplementary Section E for details). In addition, RBM recapitulates several statistical properties of natural sequences in the MSA, including conservation, covariation, distribution of lengths, and distributions of Hamming distances between sequences (see Supplementary Section B and Supplementary Figs. S2 and S3).

**FIG. 4.**
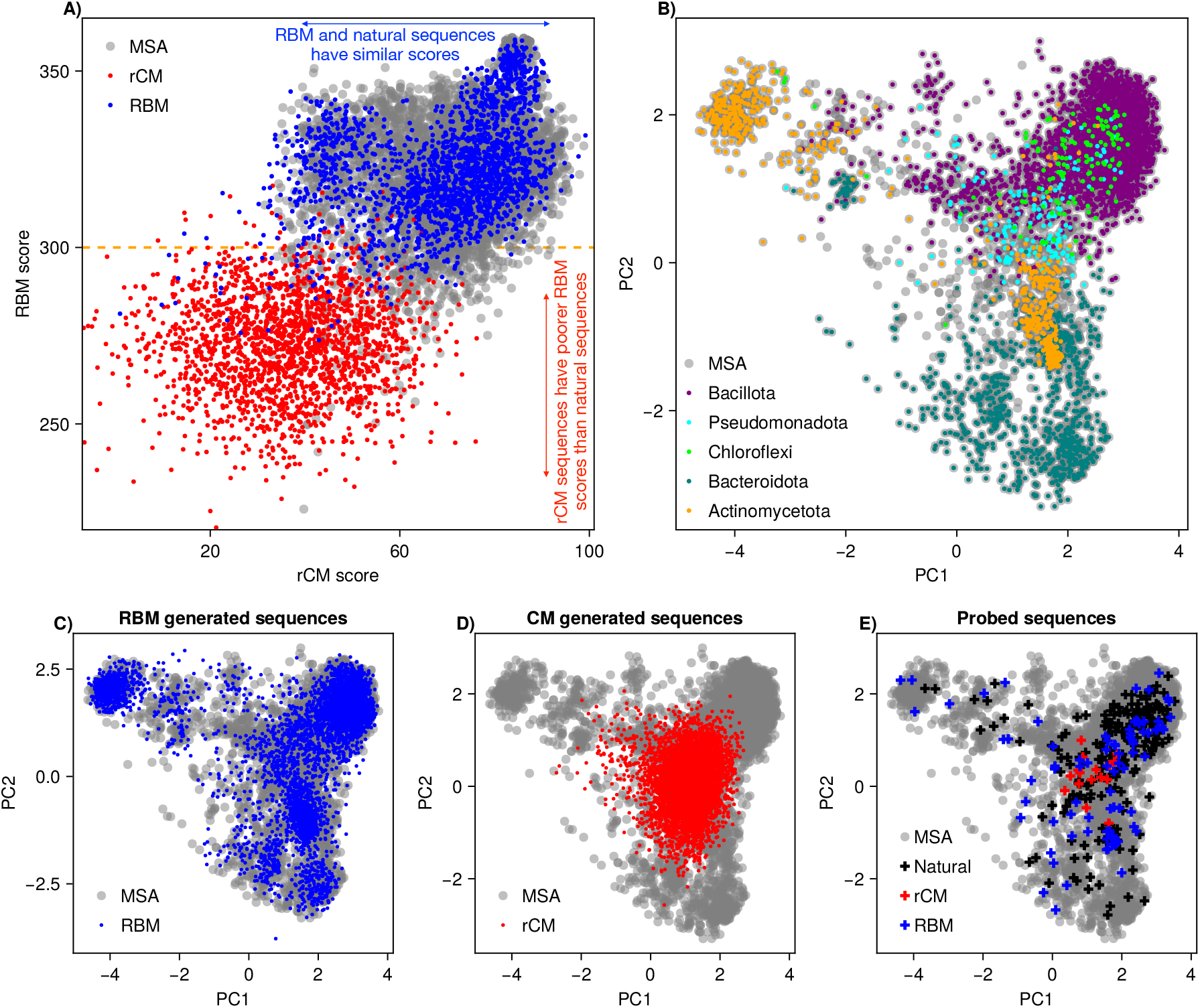
Sequence generative models. **A)** Scatter plot of rCM scores (*x*-axis) vs. RBM scores (*y*-axis), for natural sequences (gray), rCM sampled sequences (red), and RBM sampled sequences (blue). A threshold at RBM score = 300 (orange dashed line) separates the majority of rCM generated sequences from the majority of RBM and natural sequences. **B)** Projection of natural MSA sequences (seed + hits) onto the top two principal components of the MSA correlation matrix (gray). The largest taxonomic groups (with *>* 100 member sequences) are highlighted in colors. Taxonomic annotations were obtained from NCBI. **C)** Projection of RBM generated sequences (in blue) on the top two principal components of the MSA, with the natural sequences in the background (gray). **D)** Projection of rCM generated sequences (in red) on the top two principal components of the MSA, with the natural sequences in the background (gray). **E)** Projection of all probed sequences on the top two principal components of the MSA, with the natural sequences shown in background (gray). The 301 probed sequences in the first experimental batch are colored by their origin: Natural (black), rCM (red), and RBM (blue).

Next, we carry out principal component analysis (PCA) of the natural MSA. The top principal component (PC) captures a mode of variation associated to deletion of the P4 helix, as can be seen from the large number of gaps in this region (Supplementary Fig. S5). Figure 4B shows the projections of the natural sequences, annotated by their taxonomic class, onto the top two PCs. The PCs appreciably separate taxonomic clusters of natural sequences. In particular, a group of Actinomycetota, in the top left corner, have very short or no P4 helix segments. SAM aptamers can function in the absence of P4 [85], although the affinity for SAM decays with decreasing length of P4 [34].

RBM-generated sequences also span the PC space, covering all the taxonomic clusters (Fig. 4C and Supplementary Fig. S5). In contrast, rCM-generated sequences, shown in Fig. 4D remain confined to a central region. The capability of RBM to capture complex constraints in the sequence distribution allows them to model the full variability present in homologues.

We then select a fraction of the generated sequences for experimental validation, see Methods for details about the selection criteria. Their PCA projections are shown in Fig. 4E, colored by their origin (Natural, rCM, RBM), and span a wide range of the natural variability.

### B. Reactivity profiles of natural and generated aptamers with SHAPE and DMS

We resort to high-throughput chemical probing to characterize the structure of generated aptamers and their possible changes upon SAM addition. DMS mainly focuses on single-stranded A and C nucleotides, while SHAPE is sensitive to the conformational flexibility of individual nucleotides [73]. Generally speaking, paired nucleotides tend to show lower reactivities than residues left single stranded. Similarly, aptamer nucleotides bound to SAM are expected to be less reactive. SHAPE and DMS probing are routinely used to monitor aptamer structure, complexion with their ligand and structural rearrangment [4, 29–31, 33, 45, 58, 62, 70, 83].

The general result of an experiment for an aptamer is two profiles of site-dependent reactivities, one in the absence and the other in the presence of SAM (Fig. 2F). Changes in reactivities between the two conditions are expected to be informative about sites involved in interactions with SAM and in the structural switch, see Fig. 2G. However, because of the delicate nature of reactivity measurements, it is useful to benchmark the approach with natural aptamers, before turning to the analysis of the generated aptamers.

We probe a set of 208 natural sequences with SHAPE and a subset of 152 sequences with DMS in the presence or absence of SAM. These sequences are representative of Rfam ID RF00162 (Methods) and are shown by black crosses in Fig. 4E. We first present our approach and results for SHAPE-MaP. After standard processing [73], we obtain the reactivity values *r*_*i,n,c*_ assigned to each site *i*, for each aptamer *n*, and in each condition tested *c* (with or without SAM). We can then compute the difference in reactivities with and without SAM, Δ*r*_*i,n*_ = *r*_*i,n*,SAM_ − *r*_*i,n*,no SAM_. Figure 5 shows reactivity profiles from our experiments for two selected aptamers. Panel A displays the profiles obtained for *yitJ* aptamer from B. subtilis, for which a ligand-bound crystal structure was reported in [45] (PDB id: 4KQY). Interaction with SAM is confirmed by strong reactivity changes (Fig. 5B) due to the ligand at various key sites, such as SAM contacts, and sites involved in a base-triple (Fig. 1B). The *T. tengcongensis* aptamer [53] (PDB id: 2GIS) shows a similar behavior (Supplementary Fig. S26). In both cases, reactivity is low along the pseudoknot in absence of SAM, consistent with previous studies [76] that report this element is already stable in the apo form (requiring only Mg^+^ for its formation). Figure 5C, D show another aptamer (from Deltaproteobacteria), where SAM response is evidenced by reactivity drops at SAM contacts, the base-triple and also the kink-turn and the pseudoknot. Our data may thus reveal the existence of variable responses to SAM across aptamers, in terms of which sites (*e*.*g*., the pseudoknot) become more protected when SAM is present or not.

**FIG. 5.**
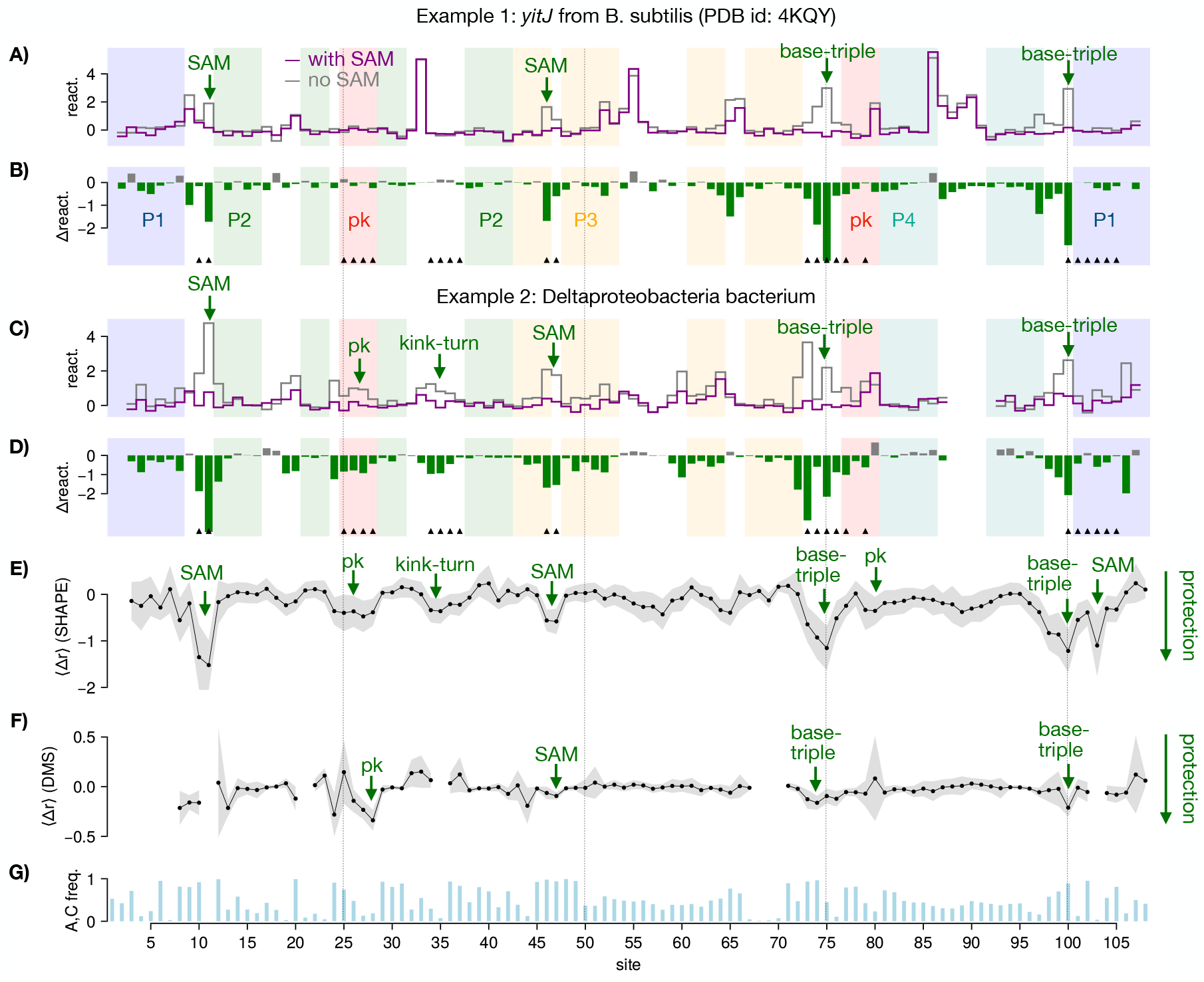
Reactivity profiles of natural aptamers with chemical probing. Key sites, involved in direct or indirect SAM interactions according to the consensus secondary structure (RF00162), are shown with black triangles. Sites 10, 11, 46, 47, 103 are in direct contact with SAM, while the remaining highlighted sites are involved in tertiary motifs that stabilize in presence of SAM: a pseudoknot (pk), kink-turn (kt), and base-triples. **A, B)** yitJ B subtilis aptamer. A. SHAPE reactivities *r*_*i*_ with and without SAM. B. SHAPE differential reactivities ⟨Δ*r*_*i*_⟩. **C**,**D)** Same as A,B for the Deltaproteobacteria bacterium aptamer. **E)** Average SHAPE differential reactivity profile Δ*r*_*i*_ over all tested natural aptamers. The thickness of the bands indicates the standard deviations. **F)** Same as E for DMS differential reactivities. **G)** Sum of A and C site-frequencies computed over natural aptamers along the sequence.

The difference in reactivities with and without SAM, Δ*r*_*i,n*_, once averaged over all probed natural sequences *n*, to better extract functional sites at the level of the family [73], defines a site-dependent Δ-reactivity template, ⟨Δ*r*_*i*_⟩_nat._, shown in Fig. 5E. We observe reactivity decreases (also called protection) for the pseudoknot (sites 25-28, 77, 79), sites involved in base triples (24, 76, 100, 73, 74) or flanking them (75), and for some of the sites directly in contact with SAM (10, 11, 46, 103). These hallmark sites, listed in Supplementary Table S2, were previously recognized for their relevance to the structural switch by previous studies using crystal structures, chemical probing, and mutagenesis experiments [33, 45, 53], see Fig. 1B. Supplementary Section Q summarizes the literature supporting these choices.

Results for DMS probing are compatible with the above findings. We report in Supplementary Fig. S34 the reactivity profiles *r*_*i,n,c*_ of the same natural sequences as in Fig. 5A-D obtained with DMS. The profiles are sparser due to the generally low reactivities of sites carrying G or U nucleotides.

Figure 5F shows the site-dependent differential reactivity profile, ⟨Δ*r*_*i*_⟩_nat._, averaged over all 152 probed natural sequences. Contrary to its SHAPE counterpart (Fig. 5E), this differential profile vanishes on most sites along the sequence. This is expected from the fact that sites may often be occupied by G or U nucleotides (Fig. 5G) and therefore weakly sensitive to DMS probing. As a result, DMS data are often less informative about SAM-induced changes than their SHAPE counterparts. However, we also observe that the few sites on which DMS differential reactivities are non zero show finer spatial resolution, e.g. on site *i* = 100, and lower sequence-to-sequence variability around the average profile (gray band around the average DMS signal), see for instance site *i* = 28 and its neighborhood. Interestingly, this latter site, which carries mostly G’s and U’s, is sensitive to DMS probing, as it is located at the junction of a stem and a loop [70].

In summary, both SHAPE and DMS average differential profiles confirm that the natural sequences probed in our experiments are mostly SAM binders and, moreover, recapitulate expected structural changes upon binding. Sequences in the seed alignment (a manually curated subset [41]) show the same average reactivity responses (Supplementary Fig. S16).

The reactivity profiles of two representative RBM generated sequences are reported in Figure 6A-D. Panels A, B show an example of a RBM-generated sequence for which the differential reactivity profiles are compatible with a global structural switch, as evidenced by reactivity changes (highlighted by arrows) in most of the hallmark sites (Supplementary Table S2), including sites in direct contact with SAM, but also the pseudoknot, the kink-turn and a base-triple motif that are known to be stabilized by the presence of SAM.

**FIG. 6.**
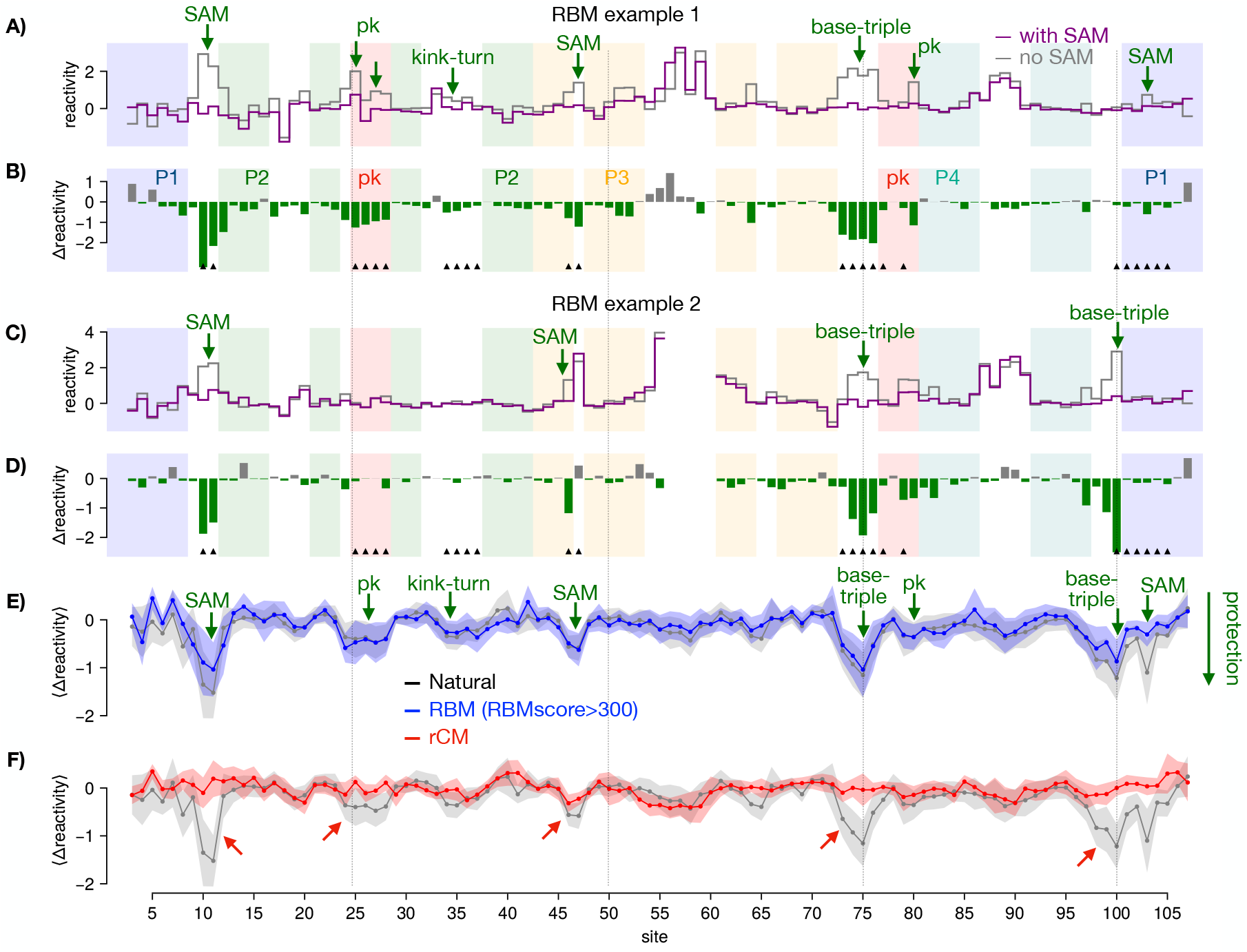
Reactivity profiles of generated aptamers with chemical probing. Black triangles refer to highlighted key sites, see Fig. 5. **A**,**B)** SHAPE reactivity and differential reactivity profiles for one RBM-generated aptamer with RBM score 321.41. **C**,**D)** Same as A,B for one RBM-generated aptamer with RBM score 357.79. **E)** Average differential reactivities in response to SAM of 54 RBM generated sequences with high RBM scores (*>* 300) (blue), across the 108 sites of the alignment. For comparison, the average differential reactivities for 204 natural sequences are shown in the background (gray). High-RBM score sequences recapitulate protection of sites involved in the structural switch in response to SAM binding (highlighted in green). **F)** Average differential reactivities in response to SAM of rCM generated sequences (red). Natural sequences are shown in background for comparison. rCM sequences fail to recapitulate the expected protections associated to the structural switch (red arrows). In both panels (E,F), the thickness of the bands indicates the standard deviation. The correlations between the site-dependent differential reactivities are 0.84 between Natural and RBM (score*>*300) (E) and 0.18 between Natural and rCM (F) with an empirical bootstrap p-value *<* 10^−6^, see Supplementary Fig. S24.

Figure 6C, D shows another RBM generated aptamer for which the differential reactivity is localized to fewer hallmark sites. In contrast to the previous example, sites at the kink-turn and pseudoknot do not exhibit significant reactivity changes in response to SAM. Reactivity changes in the base-triple and SAM contact sites strongly suggest a ligand-binding event, and are compatible with a global structural switch from an open to a closed conformation.

We emphasize that the variety in the patterns of response to SAM seen across generated aptamers is reminiscent of what is observed in natural ones. Manual inspection of all experimentally tested 201 natural aptamers, reveals that some molecules rearrange structurally upon binding SAM, others bind without significant conformational shift, and some showing no evidence of binding (no reactivity change). Examples are shown in Supplementary Figures S11 and S12. Global results of this manual inspection are summarized below.

We report in Fig. 6E the average differential reactivity profile of RBM-generated sequences having high scores (*>* 300). An excellent match with the differential reactivity profile of natural sequences is observed. In particular, protections compatible with SAM binding and the expected structural switch are found at hallmark sites. We also check that these RBM-generated sequences reproduce the reactivity response to magnesium of natural sequences (Supplementary Fig. S17). In contrast, RBM sequences with lower scores (*<* 300) show clear discrepancies (Supplementary Fig. S18) with the average profile of natural sequences.

For the sake of comparison, we show in Fig. 6F the average differential reactivities of sequences sampled from rCM (in red). Contrary to high-score RBM-generated sequences, this group of sequences shows an appreciable lack of protection at key sites, such as 10-11 (SAM contact), 25-28 (pseudoknot), 73-76 (base triples), and 103 (SAM contact in P1). Differential reactivity profiles for DMS are shown in Supplementary Fig. S35.

In summary, RBM-generated sequences with high scores exhibit, on average, the same structural response to SAM as natural aptamers. In contrast, aptamers generated by the rCM and RBM sequences with lower scores do not reproduce the characteristic features associated with structural switch (Supplementary Fig. S18).

### C. Statistical evaluation and properties of generated aptamers

Reactivity profiles are notoriously variable at the single-site level, with small differences between the distributions of reactivities expected for paired and unpaired sites. This variability can be ignored when looking at average effects over a large class of many molecules, *e*.*g*. natural or generated sequences, as done above. However, predictions for single sequences require the introduction of a proper statistical framework that integrates reactivities over a set of multiple hallmark sites and enhances the statistical signal.

SHAPE and DMS reactivities are intrinsically stochastic, and the distinction between closed and open bases should be understood in probabilistic terms. We show in Figure 7A the histogram of SHAPE reactivities of sites expected to be base-paired (teal) or unpaired (gold) in presence of SAM according to the consensus secondary structure. Unpaired sites are characterized by a different distribution of reactivities with a longer tail on high values than base-paired sites; further validating the consensus secondary structure [3] obtained by the covariation in the alignment and the large epistatic scores in Fig. 3 for secondary contacts. This picture also holds for DMS reactivity distributions, see histograms for base-paired and unpaired nucleotides in Fig. 7B.

**FIG. 7.**
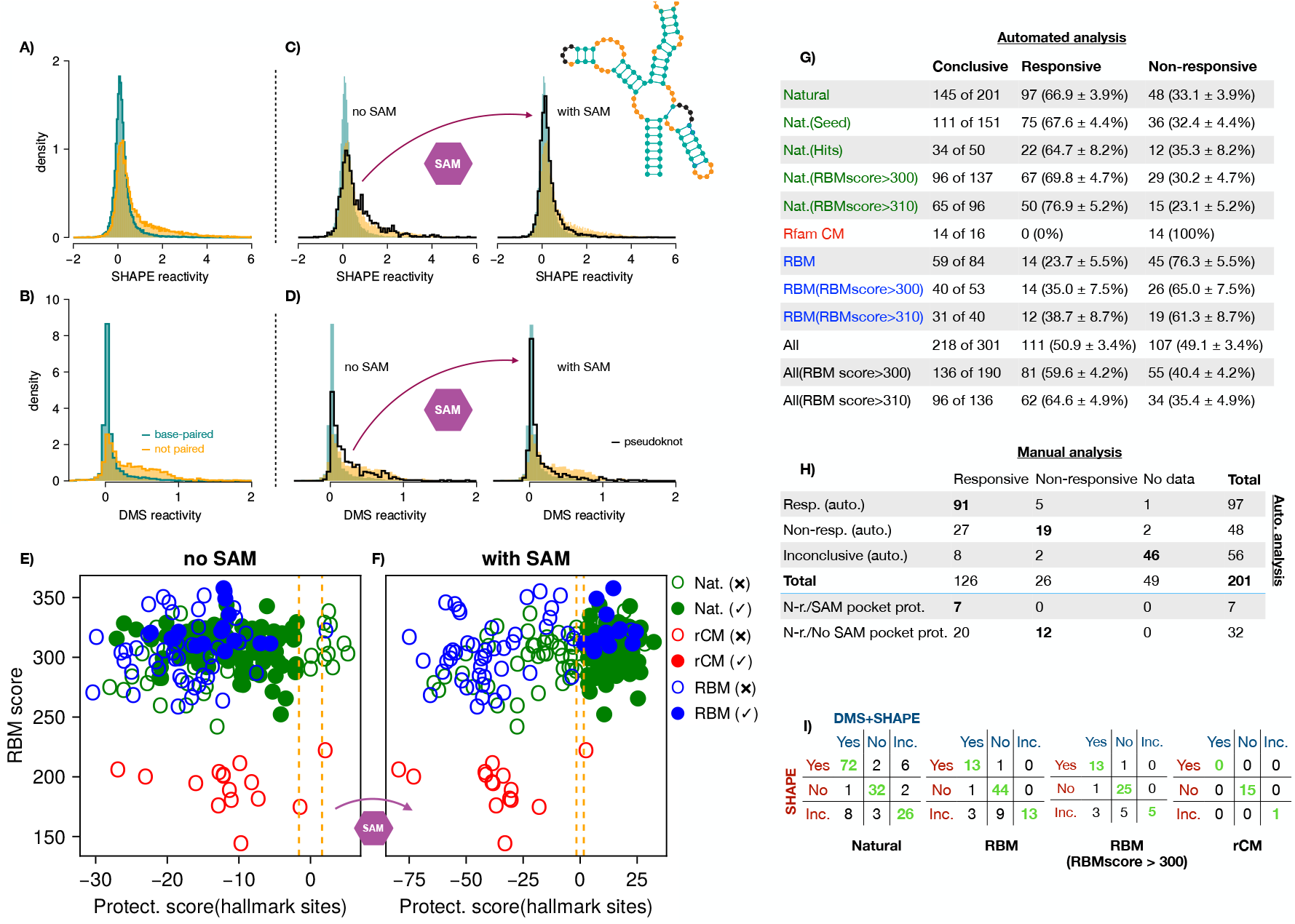
Statistical analysis of SHAPE and DMS reactivities for natural and generated aptamers. **A**,**B)** Empirical density histograms of SHAPE (A) and DMS (B) reactivities of base-paired (teal) and unpaired sites (gold) for the probed natural sequences in presence of SAM. **C**,**D)** Empirical density histogram of SHAPE (B) and DMS (D) reactivities for the pseudoknot sites (black) in the absence of SAM (left), and in the presence of SAM (right). Inset: consensus secondary structure of the SAM-I riboswitch aptamer domain, highlighting base-paired (teal) and unpaired (gold) sites. The sites forming the pseudoknot in presence of SAM (black in the inset) are not included in these histograms. **E**,**F)** SHAPE protection scores 𝒮 vs. RBM scores for all probed sequences. Panels: E) without SAM, F) with SAM. Responsive aptamers are shown with filled circles. Colors refer to the sequence origin: Natural, rCM, or RBM. Dashed orange vertical lines locate thresholds ±𝒮_0_. See Supplementary Fig. S38 for the protection scores computed from DMS data. **G)** Numbers of responsive and non-responsive aptamers in each class based on SHAPE protection scores. Error bars reflect the uncertainty in the estimated fractions based on the limited numbers of conclusive aptamers in each case (Methods). **H)** Comparison of manual (columns) and automatic (rows) classification of natural aptamers with SHAPE protection scores. The bottom two rows show how globally non-responsive (N-r.) aptamers are classified according to the protection scores of the SAM binding pocket sites only. **I)** Classification of natural, RBM-generated (all and high scores only), rCM-generated aptamers according to protection scores computed from SHAPE alone and SHAPE+DMS combined data. Yes: responsive, No: non-responsive, Inc.: inconclusive.

A clear confirmation that structural information can be extracted at the distribution level is presented in Figs. 7C and D corresponding to, respectively SHAPE and DMS data. The histogram of the reactivities of the sites associated with the pseudoknot (black) in the absence of SAM is compatible with the histogram of unpaired sites, consistently with the expected conformation of most aptamers in this condition (Fig. 1). In the presence of SAM, the histogram of pseudoknot reactivities shifts towards the distribution of paired sites. This is consistent with the occurrence of a conformational switch in most aptamers, leading to formation of the pseudoknot upon SAM addition. Similar observations can be made for the P1 helix (Supplementary Fig. S15).

Based on the findings above, we introduce a statistical approach to capture the information about structural changes present at the distribution-level in reactivity data. Let ℳ be the set of hallmark sites showing significant reactivity changes in natural aptamers in response to SAM (Fig. 5E). This set includes the pseudoknot, SAM contacts, a kink turn and sites involved in base triples (see Supplementary Table S2).

We then define, for each aptamer and each condition (with or without SAM), a *Protection Score* 𝒮 for the propensity that sites in ℳ are paired. Formally, 𝒮 is a log-likelihood ratio between these sites being all paired and all unpaired [23, 79] computed from the histograms of paired and unpaired sites in Figs. 7A (SHAPE) & B (DMS). The score also accounts for sampling noise arising from limitations on the sequencing depth [73], which may strongly impact some experiments, see Methods. We emphasize that aggregating multiple sites in the score is crucial to reduce the statistical noise intrinsic to chemical probing measurements (see Fig. 2G and Supplementary Figures S25, S39). Furthermore, when SHAPE and DMS data are available for the same aptamer, the two protection scores can be summed up to obtain a more robust predictor, which we refer to as DMS+SHAPE below.

Figure 7E reports the SHAPE protection scores without (left) and with (right) SAM for natural aptamers. For aptamers switching in response to SAM, we observe that shifts 𝒮 from negative values in the absence of SAM (indicating the hallmark sites are likely to be unpaired) to positive values in the presence of SAM (indicating that these sites are involved in an interaction). Hereafter, we will call

- *responsive* every aptamer, whose protection score 𝒮 is lower than −𝒮_0_ in the absence of SAM and larger than +𝒮_0_ in the presence of SAM;
- *non-responsive* every aptamer, whose protection score 𝒮 is larger than −𝒮_0_ in the absence of SAM or lower than +𝒮_0_ in the presence of SAM;
- *inconclusive* if either score (with or without SAM) is smaller than 𝒮_0_ in absolute value.

We adopt a 5-fold significance threshold 𝒮_0_ = ln(5), see Methods.

As shown in Fig. 7G, aptamers responsive according to SHAPE protection scores (both natural and generated) tend to have high RBM scores. In particular, 35% of RBM-designed aptamers with RBM score *>* 300 structurally switch in response to SAM, exhibiting significant responses in the hallmark sites. These sequences differ by 10 to 30 residues from the closest natural sequences (Supplementary Fig. S4). In the case of failing RBM-generated sequences, the structural motifs (pseudoknot, P1, etc.) remain either protected even in the absence of SAM, or reactive in the presence of SAM. We find that most of the 45 RBM non-responsive sequences fail in the second manner: they do not have the necessary contacts even in presence of SAM. Non-responsive natural sequences can fail in both ways. None of the sequences generated with rCM is functional, possibly due to the inability of rCM to model tertiary motifs [22, 56]. Let us stress that the number of inconclusive sequences is deeply affected by the read depth of the experiment, with lower depth leading to more inconclusive sequences, see Methods Section I for a detailed analysis of this effect.

The outcomes of the manual and automated analysis based on protection scores are compared in Fig. 7H. The two analyses are in agreement for 110 out of the 142 (77.5%) aptamers where they are both conclusive. Out of the 32 disagreements, 27 (19% of conclusives for both) are responsive in the manual analysis but not in the automated one. Manual inspection focuses on localized responses that are evidence of SAM binding. The protection-score-based analysis is more stringent, requiring a global response compatible with a structural switch across most hallmark sites. The automated analysis can also detect local responses, by focusing on smaller subsets of the hallmark sites (see last two rows of Fig. 7H, and Supplementary Section N).

To provide evidence for the reproducibility of our results, we perform two replicates of the experiment, the first one on the total set of 301 natural and artificial sequences and the second one on the 201 natural sequences only, see Supplementary Section K for a detailed description. Although some aptamers in the first replicate exhibit an overall lower response to SAM (natural and artificial), the fractions of responsive sequences in each group are consistent with the results reported in Fig. 7. Moreover, 80% of identified responders in the replicates were also responsive in the first experiment, confirming the robustness of the automated analysis (Supplementary Fig. S19).

The results above, obtained from SHAPE data, are corroborated by chemical probing with DMS. Using Eq. (11), we compute protection scores combining SHAPE and DMS reactivity data for enhanced discrimination. Fig. 7I compares the results from SHAPE alone and combined DMS+SHAPE. Let us focus on natural sequences first. SHAPE and DMS+SHAPE provide the same classification (responsive, non-responsive, or inconclusive) for about 86% of the aptamers. Among the remaining 14%, more than 12% are inconclusive for one of the two approaches, and SHAPE and DMS+SHAPE disagree on less than 2% of the aptamers only.

Similar patterns are observed for RBM-generated aptamers. For RBMscore *>* 300, we obtain consistent responsive rates (ratio of the numbers of responsive and conclusive sequences) of 35%, whether estimated from SHAPE or DMS+SHAPE data. Interestingly, 48% of RBM sequences that were inconclusive with SHAPE alone can be classified with DMS+SHAPE, with one quarter responding and three quarters not responding. No rCM-generated aptamer is considered as responsive by either SHAPE nor DMS+SHAPE. A complete comparison of the analysis of the SHAPE and DMS data is reported in Supplementary Fig. S37.

Inspired by previous experimental observations for other riboswitches [36, 90] and Sabatier’s principle for enzymes, which require intermediate substrate binding energies for proper function [65], we compute the thermodynamic energies brought by P1 helix formation using the Turner energy model as implemented in the ViennaRNA package [44] (Methods). Figure 8A shows that the sequences that respond to SAM through P1 helix stabilisation are confined to a thermodynamic energy window ranging from −10 to 0 kcal/mol. Similarly, pseudoknot (Pk) formation in response to SAM tends to occur for aptamers having a Pk pairing energy comprised between −8 and −3 kcal/mol (Fig. 8B). As P1 and Pk consists of, respectively, 8 and 4 base pairs, the flexible energetic window spans a range of 1.25 kcal/mol per base pair in both cases, close to a weak base-pairing energy [13, 44]. The leftmost panels in Fig. 8 show that RBM samples preferentially have pairing energies in this intermediate band for both P1 and Pk, and are thus compatible with the structural switch required for riboswitch function.

**FIG. 8.**
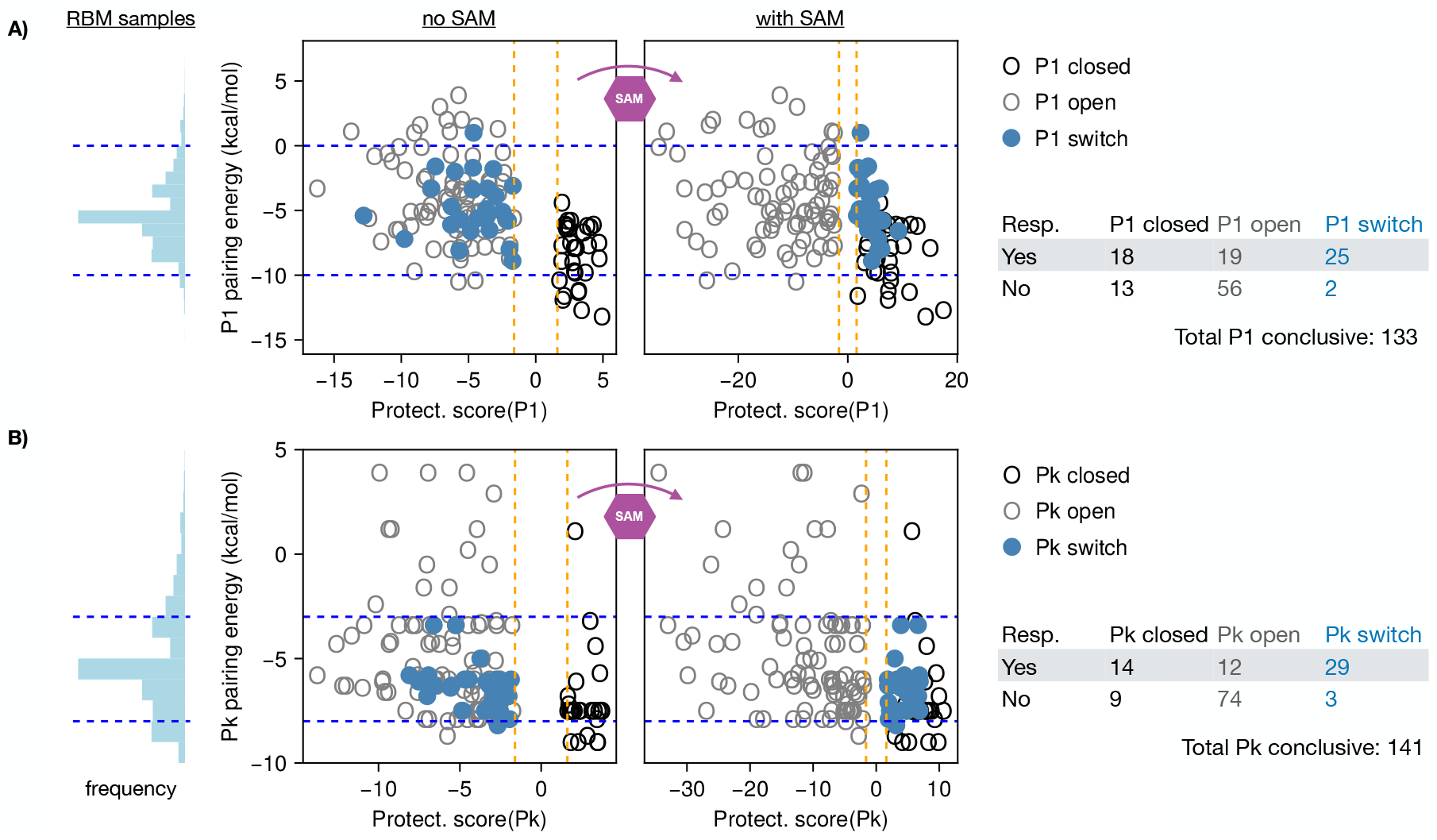
Local responses along P1 and the pseudoknot (Pk) require intermediate pairing energies. **A)** Left panel shows the histogram of Turner pairing energies for P1 (computed with the ViennaRNA package [44]) of a random sample of RBM-designed sequences. The following panels show the pairing energies without (middle) and with (right) SAM for the aptamers probed in the first batch vs. the protection scores 𝒮 (*P* 1) obtained by choosing for the hallmark set ℳ the sites in P1 only. Aptamers are colored according to their response: if 𝒮(P1)*>* 𝒮_0_ in both conditions, P1 is always closed (open black circle); if 𝒮(P1)*<* −𝒮_0_ in both conditions, P1 is always open (open gray circle); if 𝒮(P1) crosses from one side to the other, the motif switches in response to SAM (filled light blue disks). Note that only aptamers for which the P1 response is conclusive are shown (133 aptamers). The table then lists the numbers of aptamers that are responsive to SAM, compared to a local response in P1 only. **B)** Same as A), but for pseudoknot (Pk) sites.

The tables in Fig. 8 give a summary of these results. Interestingly, 25 out of the 27 aptamers that stabilize P1 in response to SAM are also responsive, in the sense of Fig. 7E,F, and show broad structural responses in other Hallmark sites (Supplementary Table S2). Similarly, 29 out of 32 aptamers that stabilize Pk are also responsive. On the other hand, out of 112 identified responsive aptamers, in natural and artificial sequences, only 19 do not stabilize P1 significantly after binding SAM. These aptamers must exhibit significant compensatory stabilization of other structural Hallmark motifs from Supplementary Table S2. It is important to note that P1 can have a more flexible behavior in the full riboswitch due to competitive interaction with the expression platform, compared with the aptamer only. As shown in Supplementary Fig. S41 the P1 helix can be destabilized in the full riboswitch context, whereas other helices like P2 or P4 are not affected, see Supplementary Figs. S42 and S43. Taken together, these results are consistent with the known importance of the pseudoknot and P1 in the response of the aptamer.

Notice that, in the central panels of Fig. 8, we show only aptamers for which the the statistical analysis yields a conclusive response for P1 or Pk. Inconclusive aptamers also tend to have intermediate pairing energies for P1 and Pk, consistent with structural flexibility (*e*.*g*. breathing).

### D. Further explorations of RNA switch diversity through design

We then perform a second batch of design and experimental validation to further assess the limits of our generative models. We probe a total of 450 generated aptamers, whose sequences are projected onto the MSA PCs in Fig. 9A.

**FIG. 9.**
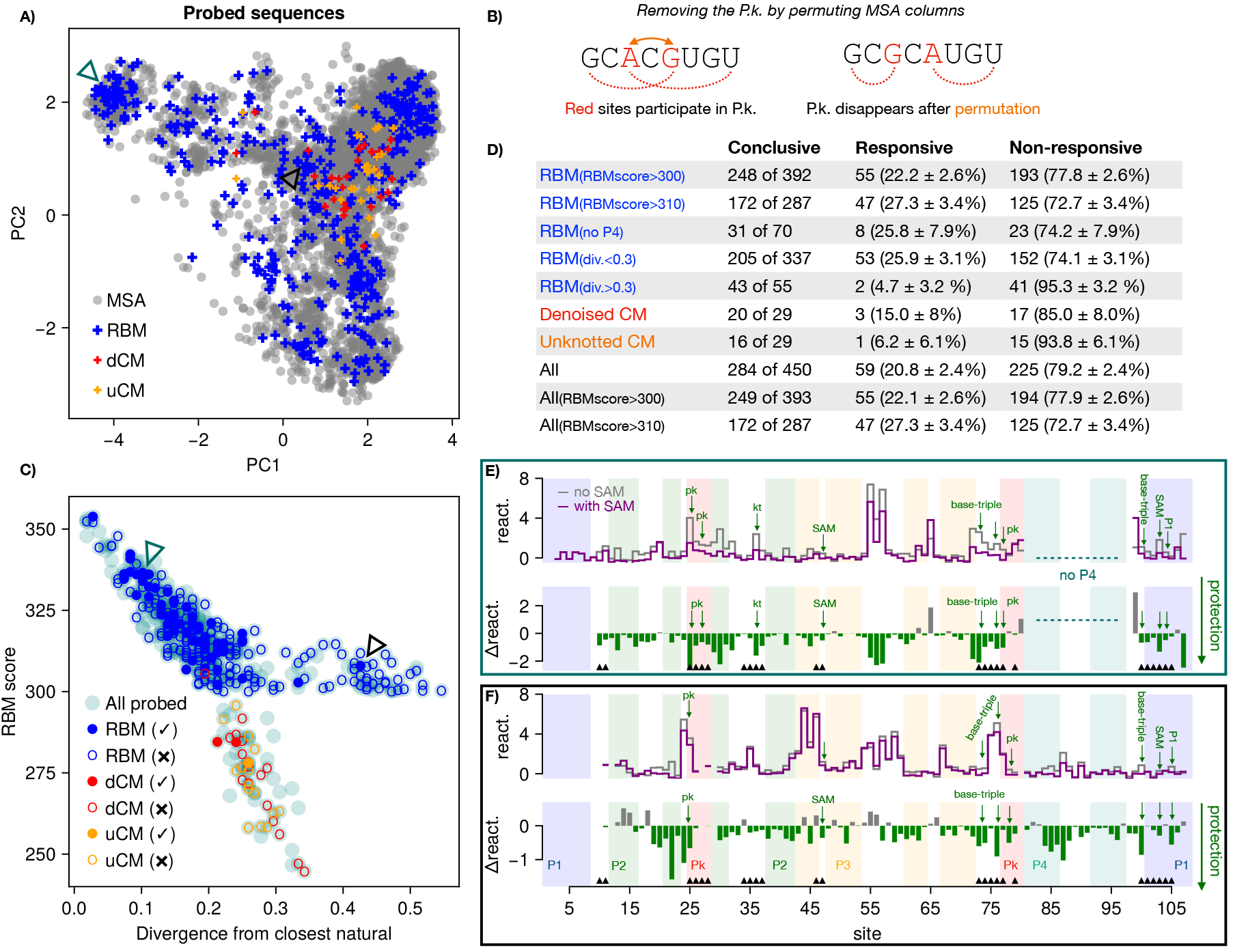
Additional generation of sequences. **A)** Projection of sequences probed in second set of designed sequences along the PCs of the natural MSA, colored by origin: RBM, Denoised and Unknotted CM (dCM, uCM). The full natural MSA is shown in background (gray) for comparison. **B)** Diagram explaining the uCM, where the pseudoknot is undone by permuting a specific set of columns in the MSA. In this manner, a CM can model covariation along a pseudoknot. See Methods for details. **C)** Divergence from closest natural sequence in the MSA (fraction of sites that differ) vs. the RBM score, for all sequences probed in the second experiment. Colored circles correspond to aptamers that switch in response to SAM (fill color) or not (empty), with the color indicating the sequence origin: RBM (blue), dCM (red), and uCM (orange). Sequences for which our analysis is inconclusive are shown in light cyan. **D)** The table summarizes the numbers of switching sequences in each group. **E)** Reactivity profile of example responsive RBM generated sequence with no P4 (indicated by teal triangle in A,C). **F)** Reactivity profile of example responsive RBM generated sequence at large distance from natural sequences (indicated by black triangle in A,C).

First, we sample sequences with the RBM model exhibiting higher distances from their closest natural counterpart, focusing on RBM scores *>* 300. In addition, as some natural sequences lack P4, we retain a subset of RBM generated sequences having severely diminished P4 lengths. These are clearly seen in Fig. 9A, clustered at the top-left corner of the plot (recall the top PC1 represents P4 deletion). We also sample more RBM sequences of high scores (*>* 300 and *>* 310) to obtain better statistics on the fractions of working aptamers.

Second, we consider two variations of rCM, which is over-regularized to capture distant sequences in Rfam alignments [40]. We rebuild a non-regularized CM trained on the same MSA, which we call Denoised CM, or dCM for short (Supplementary Fig. S8 and Methods). Furthermore, as CM are unable to model pseudoknots, we devise a permutation of the MSA columns that undoes the pseudoknot, see Fig. 9B. We trained a new CM variant on the permuted MSA, that we call Unknotted CM (uCM), properly taking into account covariations in the pseudoknot. We generate sequences with such model and permute back the pseudoknot columns (Methods).

Interestingly, both dCM and uCM share some of the properties of rCM noted previously. First, CM-generated sequences from all variants have predominantly low RBM scores *<* 300, see Supplementary Fig. S9. Second, CM generated sequences exhibit restricted diversity, concentrating in a central region of the PCA plot, as in Fig. 4D. In particular, all CM are unable to generate sequences without the P4 helix. Sequences sampled from uCM have better complementarity and Turner energies favorable for base-pairing along the pseudoknot.

We then perform SHAPE-MaP experiments and analysis. Results are summarized in Fig. 9C, and show the RBM scores of the probed aptamers against the Hamming distances to the closest natural sequence.

Out of the 248 conclusive RBM sequences in the second batch, 22% switch in response to SAM (Table in Fig. 9D). The percentage of responsive among the sequences closer to the natural ones is higher and compatible to what obtained in replicate 1 considering the error bars, see Table 7G.

Moreover, 25% of the RBM aptamers having P4 length ≤ 1, respond to SAM; an example reactivity profile is shown in Fig. 9E. We also find a few switching aptamers differing by 30 to 50 sites from any natural sequence. An example reactivity profile for such sequence is shown in Fig. 9F. The reactivity profile is compatible with the consensus secondary structure, with most reactivity peaks tending to occur in unpaired loops (except a portion of P3 that remains reactive), and an overall protection in response to SAM compatible with binding and stabilization of the aptamer. Notice that RBM generate diversity not only in highly variable parts of the sequence, but also in more conserved sites (Supplementary Fig. S4).

These results support the generalization ability of the RBM. In contrast, only 3 out of 20 conclusive dCM samples switch in response to SAM (15%), and only 1 out of 16 from uCM (≈ 6%). Thus the dCM and uCM perform better than rCM, but not as good as RBM.

## DISCUSSION

In this work, we focused on the design of small molecular RNA switches, capable of changing conformation upon binding to a metabolite. Building such aptamers is a first step in the design of functional switching RNA, with many potential applications in developing laboratory tools for gene function studies, metabolic engineering or drug design, as they can be used to regulate gene expression [1, 26, 37]. The design of allosteric and regulatory RNA is also key to DNA-RNA computing, and to the investigation of possible scenarios for the origin of life [12, 38, 71].

State-of-the-art design methods for RNA are based on computational frameworks to fold sequences in a given secondary structure from the knowledge of thermodynamic parameters for the pairing energies [87], possibly including tertiary elements such as pseudoknots [93]. Such methods have been used to obtain sequences with bistable secondary structures [25] and extended to take into account both positive and negative design elements [63, 93], as well as to community-based rational design [43]. Our design method, based on the unsupervised generative architecture of Restricted Boltzmann Machines, differs in two key ingredients: i) it exploits the sequences (of SAM-I riboswitch aptamers) sampled through evolution and collected in databases, building upon the frameworks introduced in homology and covariation detection [19, 51, 57, 63, 92]; ii) it encompasses, through learning of a unique parametric model, the arrangements of nucleotide motifs allowing natural sequences to acquire adequate secondary and tertiary structures and to undergo an allosteric response to metabolite binding.

We have verified that the RBM model learned from sequence data encode nucleotide-nucleotide contacts in the secondary structure and in the pseudoknot, performing at the same level as pairwise Potts/DCA models previously introduced to this aim [19, 92]. In contradistinction with those pairwise interaction-based models, RBM are capable of extracting extended nucleotide motifs, *e*.*g*. overlapping one or more structural elements. A major advantage of the shallowness of the RBM architecture is that these motifs can be readily accessed and interpreted through inspection of the weights (Fig. 2D and Fig. 3B&C).

To assess the sequences designed by our computational models, as well as the natural sequences belonging to the SAM-I riboswitch aptamer family, we have carried out high-throughput SHAPE and DMS screening. We have introduced and implemented a statistical pipeline to analyze the measured reactivities, based on a likelihood ratio between reactivity distributions of paired/unpaired nucleotides, called protection score [23, 79]. Our analysis takes advantage of the closely related statistics of the ensemble of tested sequences and their shared consensus secondary structure. As it does not rely on a biophysical implementation of the Turner model [44], tertiary contacts such as pseudoknots, which are essential to model complex conformational changes such as those occurring in riboswitches, are fully accounted for. Last of all, our pipeline is fully automatic and does not require manual annotation, which is time consuming for high-throughput screening.

Our analysis of SHAPE and DMS data shows that RBM are able to successfully design artificial SAM-Iriboswitch-like aptamers. Of the sequences generated with high RBM scores for which our conservative statistical analysis could reach a clear conclusion, 35% could be classified as responding to SAM in the first replicate. This fraction is significant, and shows that RBM are effective as generative models of complex RNAs. It is, however, lower than the one (70%) of natural sequences deemed as responsive according to the same criterion. We emphasize that the fraction quoted above varies with the constraints considered during the generation process. For instance, up to 50% of RBM-generated sequences were recognized as responsive when the fraction of mutated residues with respect to the closest natural sequences is of 20% (over 108 nucleotides). Pushing generation to the limits as in the second experiment made the global fraction drop down to 22%, but allowed us to generate functional aptamers with as many as 46% of mutations with respect to the closest known natural aptamers. Moreover, RBM can design responsive aptamers lacking the P4 helix (as in some natural variants), whereas CM are unable to generate such sequences.

The success of our design approach crucially relies on the capability of RBM to capture nucleotide motifs responsible for tertiary structural elements. This statement is supported by the fact that CM, while capturing the local conservation and secondary structure of the Riboswitch family, has significantly lower generative performance (≃11%, Denoised & Unknotted). In addition, RBM generate flexible structural elements, with intermediate pairing energy values, permitting them to open and close depending on the metabolite presence. From this point of view, while RBM have already been used to generate functional proteins [47] or DNA aptamers [21], this is the first time they are shown to be able to design allosteric biomolecules.

Besides the responsive/non-responsive classification based on protection scores, a pattern of phenotypes is observed in the generated sequences through manual inspection of the reactivity profiles and of their changes with SAM presence. Among the natural sequences that fail to qualify as fully responsive with our automatic statistical pipeline, many are manually seen to exhibit local reactivity responses to SAM indicative of binding (Fig. 6C,D).This response can manifest itself as a change in the reactivities of the sites related to the SAM binding pocket, or involved in P1, in the pseudoknot, or in any of the three base triples. Similar patterns are encountered in RBM-generated sequences, see Supplementary Section N. The distinction between binding to SAM and being able of undergoing conformational change we observe here agrees with recent directed evolution experiments. It was reported that evolving RNA for ligand binding alone often failed to produce functional regulatory RNAs [36, 90], highlighting the importance of the structural switch. More recently, Capture-SELEX, in which conformational change triggered by the ligand and optimal switching time are selected for was proposed for this purpose [1, 6, 26]. Supervised classifiers, learned from the experimental sequences were shown to be able to predict the functionality of the molecules [1, 26, 37].

Since this paper was posted on the archive, two works have developed generative models of structured RNA: [10] proposes a parsimonious DCA-like model, which promotes sparsity of model weights and validated experimentally generation of a tRNA family; [80] introduced a combination of Variational AutoEncoders with CM and showed that their model was generative over various ribozyme families. Our work differs in that it presents the first example of design of RNA molecules exhibiting structural switching upon metabolite binding. We have further performed a comparative analysis of the two-layer RBM-based generative model to the deep variational autoencoder (VAE) models of [80] on our data. RBM seems to detect key features in natural sequence data not extracted by VAE: VAE give similar scores to RBM-generated and natural sequences, while RBM scores are higher for natural than for VAE-generated sequences (Supplementary Fig. S7). Further investigations, in particular experimental tests, would be necessary to better understand these preliminary results.

We plan to investigate more deeply the mechanisms for conformational switching in different subfamilies of the SAM-riboswitches family. We emphasize that the RBM-based design of artificial RNA sequences can be carried out for any RNA family for which homologous sequences are available. As shown in SI, Section L, we have also learned RBM models on the aptamer domains of three other riboswitch families: cyclic di-AMP[2], Cyclic di-GMP-I[78], and Glycine riboswitches [48]. The designed sequences are of high computational quality, as proven by the similarity of the scores assigned by the RBM and the CM models and of their statistics with respect to natural sequences, see Supplementary Fig. S10.

In addition, our approach could be extended to the modeling of complete SAM riboswitches by including the expression platform. In this context, it would be interesting to perform functional tests of the designed aptamer, *e*.*g*. in yeast constructs with a GFP reporter protein [26]. It would be in particular interesting to check if the increased flexibility of P1 helix in presence of the expression platform increases the percentage of molecules responding to SAM among the tested ones. Due to the strong interactions between the latter and P1 (Supplementary Fig. S41), the RBM should be trained on full riboswitch sequences, including both the aptamer and the expression platform. However, full riboswitch sequences exhibit significant length variability, with hard-to-align regions, which would require some modifications in our model such as introduction of a convolutional layer.

Lastly, RBM could also be used to design other RNAs, including longer and more complex ribosomal RNA.

## METHODS

### A. Multiple sequence alignment of SAM-I riboswitches

The RF00162 family from the Rfam database [41] groups sequence homologs of the aptamer domain of the SAM-I riboswitch. We downloaded a manually curated seed alignment from Rfam (version 14.7), containing 457 aptamer sequences supported by literature evidence. These seed sequences are aligned to a consensus secondary structure (shown in Fig. 1B) that has been informed by the holo-form of SAM-I riboswitch crystal structures [45, 53]. After removing extended stems and variable loops, labeled as insertions in the alignment, we obtain 108 matched positions (including gaps that mark deletions) spanning four helices that interleave around a central four-way junction. We trained a covariance model (CM) [22] on this seed alignment using Infernal [56] with default settings. Following standard protocols [40], we acquired 6161 additional sequences from Rfam, collected from genome databases and filtered for significant matches to the CM. We constructed a multiple sequence alignment (MSA) with these sequences, that we refer to as the full MSA, to distinguish it from the seed MSA consisting only of the 457 manually curated seed sequences. The sequence conservation logo of the full MSA is shown in Fig. 1C.

### B. Infernal pipeline

Infernal [56] is a set of computational tools to facilitate modelling RNA sequence families under a profile stochastic context-free grammar (pSCFG) formalism, also known as covariance models (CM) [22]. A CM is capable of modelling the conservation profile of important sites along the sequence, as well as correlations between distant sites required by the complementarity of base-pairs in a given secondary structure. Infernal is routinely used in the maintenance of alignments in the Rfam database [40, 41]. We employed Infernal to construct the RF00162 full MSA, that we use to train the RBM.

By restricting to covariations in the secondarystructure, CM can be efficiently implemented with dynamical programming algorithms [22]. However, these assumptions also imply that CM is unable to include additional constraints in the probabilistic sequence model, such as pseudoknots and other tertiary contacts in the 3-dimensional fold of the RNA molecule.

#### 1. Rfam CM

The Rfam database associates a CM model to each family, trained on the seed alignment, that is used to scan large genomes for significant sequence matches to the family (hits). The raw CM model downloaded from Rfam is significantly regularized so that it is more effective in fetching far homologs of a family in deep genome searches [55]. We will refer to this CM model as Rfam CM, or rCM for short.

#### 2. Denoised CM

Since rCM is strongly regularized, in this work, we also trained a CM model variant on the full MSA, with no regularization, which we call Denoised CM, or dCM for short. This model reproduces more closely some statistics of the full MSA (conservation and covariances associated with the secondary structure).

#### 3. Unknotted CM

A CM model cannot model pseudoknots and other tertiary contacts. Based on our knowledge of the consensus secondary structure of the SAM-I riboswitch aptamer (Fig. 1B), we devised a third CM model able to account for sequence covariation in pseudoknot sites constructed as follows. Columns 77–80 of the MSA, corresponding to the sites on the 3’-end part of the pseudoknot, were moved and inserted after site 28, right next to the the sites at the 5’-end of the pseudoknot. In this way, the pseudoknot is “unknotted”, and is now representable in the CM model as part of a pseudo-secondary structure corresponding to the permuted MSA. Accordingly, we proceeded to train a CM model on the rearranged full MSA. We call the resulting model Unknotted CM, or uCM for short.

#### 4. Sampling the CM

To better understand the limitations of CM models and the advantages of RBM, we sampled 10000 sequences from each of the three CM described above. For the uCM, the rearranged columns are permuted back to their original positions after sampling. We used Infernal’s <monospace>cmemit</monospace> program with default parameters, and without insertions. Infernal computes a score of sequences aligned to the CM, related to the likelihood of the CM to emit a given sequence (also called *bit-scores*). We computed this score using <monospace>cmalign</monospace>, with -g (global) option to avoid local approximations [55].

### C. Restricted Boltzmann machines

Restricted Boltzmann Machines (RBM) [35] are bipartite graphical models over *N* visible variables **v** = {*v*_1_, *v*_2_, …, *v*_*N*_} and *M* hidden (or latent) variables {**h** = *h*_1_, *h*_2_, …, *h*_*M*_}, see Fig. 2A. Here *N* = 108 corresponds to the sequence length of the RF00162 alignment, and *v*_*i*_ encodes the nucleotide present at position *i* of a sequence. For RNA, *v*_*i*_ can take one of *q* = 5 possible values, corresponding to the nucleotides A, C, G, U, and the alignment gap symbol (⊟). The hidden variables *h*_*µ*_ are here real-valued. The two layers are connected through the interaction weights *w*_*iµ*_. An RBM defines a joint probability distribution over **v** and **h** through

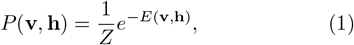

where *Z* is a normalization factor, known as the partition function, and the energy *E*(**v, h**) is given by

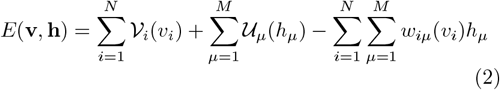

The functions 𝒱_*i*_(*v*_*i*_), 𝒰_*µ*_(*h*_*µ*_) are potentials biasing the distributions of single units. The visible units *v*_*i*_ can take a finite number of possible values, and therefore the quantities 𝒱_*i*_(*v*_*i*_), also called ‘fields’, can be stored as a *q × N* matrix. Similarly, the weights *w*_*iµ*_(*v*_*i*_) can be stored as a *q × N × M* three-dimensional tensor. The hidden variables, on the other hand, are continuous, and we chose to parameterize their potentials with the double Rectified Linear Units (dReLU) form proposed in [86],

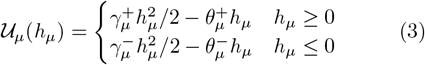

with real parameters 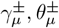, satisfying 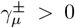. The dReLU is an attractive choice because it is expressive enough to cover several interesting settings. When 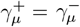 and 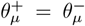, Eq. (3) becomes a quadratic (*i*.*e*., Gaussian) potential, and is closely related to Direct-Coupling Analysis models popular in protein sequence modelling [14, 19, 54, 68, 72, 91]. However, the Gaussian choice is unable to parameterize more than two-body interactions, which can be a limitation in RNA structure where some interactions are known to involve more than two sites (*e*.*g*. stacking interactions [13, 94]), as well as functional interactions that can span complex, extended structural and sequence motifs. dReLU can also adopt a bimodal form when 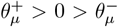, which is helpful for clustering.

The likelihood of visible configurations under the RBM can be obtained by marginalizing over the states of the hidden units:

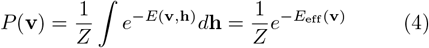

where −*E*_eff_(**v**) is the resulting RBM score that incorporates effective interactions arising from the marginalized latent variables (see Fig. 2C):

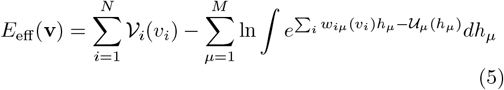

Although evaluating *P* (**v**) is computationally difficult (because the partition function *Z* is intractable), Eq. (5) shows that the score −*E*_eff_(**v**) can be computed efficiently.

The computation of epistatic scores follows [86]. Further details about our RBM implementation for training and sampling are given in Supplementary Section A.

### D. Biophysical energy calculations

We computed biophysical pairing energy predictions for the formation of P1 and the pseudoknot of various sequences using the Turner energy model, as implemented in the ViennaRNA package [44], with the RNAeval program.

- For the P1 helix, we computed the energy difference of each sequence in the consensus secondary structure of the aptamer domain, where P1 is paired (Fig. 1B), and in a conformation where P1 is unpaired (Fig. 1A).
- To estimate the pairing energy associated to pseudoknot formation, we used RNAeval on a virtual secondary structure where only the pseudoknot sites are base-paired, and all other sites are unpaired. We then considered only interior loop contributions to the resulting folding energy.

Note that, in both cases, intrinsic limitations of the ViennaRNA algorithmic implementation imply that we cannot model the pseudoknot together with other structural elements (and other tertiary contacts).

### E. Selection of sequences for first batch

We probed a total of 306 sequences, breaking down as follows.

#### RBM sequences

We generated sequences from the RBM by Gibbs sampling. Equilibration was assessed by monitoring the average score of the sample. We found that 5000 steps were more than sufficient. We then sorted these sequences by their RBM score (−*E*_eff_), and selected 70 sequences at random, uniformly spanning the range of scores observed in the sample. The table of sequences and their associated RBM scores is reported in the Supplementary Code listing [15], see Section N.

#### Infernal sequences

We then sampled sequences from the rCM of the RF00162 family, downloaded from Rfam. We used the Infernal cmemit program (see Methods) to sample a large batch of sequences. We selected 30 sequences uniformly spanning the range of bit-scores of the samples.

#### Natural sequences

We selected 151 sequences members of the seed MSA and 55 sequences members of the full MSA, as described in Section A. The selected natural sequences are diverse, spanning various taxonomic classes (see Fig. 4B). A listing of probed sequences can be found in Supplementary Data 2.

### F. Selection of sequences for second batch

In the second experiment, we generated a total of 450 sequences to be probed, of different origins. We considered:

- 58 CM sequences, with 29 from uCM and 29 from dCM (see Section B for definitions of these CM variants).
- 392 sequences sampled from the RBM, filtered to have RBM scores *>* 300. In particular, 49 of them were selected because they had no P4 helix, while 100 of them were selected because they had larger Hamming distances from any natural sequences.

The full list of designed sequences is provided as part of the Supplementary Code listing, see Section N.

### G. Selection of sequences for DMS probing

We selected a subset of aptamers from batches 1 and 2 for DMS probing. From batch 1,

- 84 sequences generated by RBM;
- 16 sequences generated by rCM;
- 152 natural sequences.

From batch 2,

- 102 sequences generated by RBM;
- 10 sequences generated by uCM or dCM.

The full list of sequences probed by DMS is provided as part of the Supplementary Code listing, see Section N.

### H. Chemical probing experiments

#### 1. RNA preparation

DNA oligonucleotides representing the 206 SAM-I natural sequences, and the two batches (100 and 450) of artificial sequences, preceded by the T7 promoter (5’CGGCGAATCTAATACGACTCACTATAGG3’) and followed by a tag sequence representing a 10 nucleotide barcode unique for each aptamer and a primer binding site, were purchased as an oligonucleotide pool (Twist bioscience). The Tag sequence was designed to avoid interference with the aptamer secondary structure using RNAFold [44] (see [32] for the tag design method). The oligo pool was PCR amplified using the T7 promoter as forward primer and five different reverse primers (5’GGAAGGAGGCGGGCAGACG3’, 5’CGTATTACCGCGGCTGCTGG3’, 5’CGACGAGATAGGCGGACACTGG3’, 5’CGACGAGATAGGCGGACACTGG3’, 5’GAAGTCGTAACAAGGTAGCCGAT3’), provided in Supplementary Data 1. RNA was transcribed, prepared, and checked for the absence of aberrant products on a 1% agarose gel [20]. See Supplementary Sec. S for details.

Read depths vary with the choice of the primer, see Supplementary Figs. S27, S28, S29. As explained in Methods Section I we have verified that our statistical analysis give consistent rates of responsive aptamers, even for primers with lower coverage.

#### 2. SHAPE and DMS probing

SHAPE chemical probing was performed as described previously [73]. Briefly, 10 pmol of RNA were diluted in 12 µL of water and denatured for 3 min at 85°C. Then, 6µL of 3X pre-warmed folding buffer with or without magnesium (0.3M HEPES pH 7.5, 0.3M KCl, 15mM MgCl2) were added and the solution was allowed to cool down to room temperature. Samples were then incubated at 30°C for 5 min. S-adenosyl-methionine (SAM) was added at final concentrations of 0, 0.1 or 1mM and samples were incubated 15 min at 30°C. 9 µL (corresponding to 5 pmoles) were aliquoted and 2 µL of 50 mM 1M7 (1-Methyl-7-nitroisatoic anhydride) or DMSO (Mock reaction) was added and allowed to react for 6 min at 30°C. For dimethyl-sulfate (DMS) probing, 0.9µL of 600mM DMS stock solution (or 0.9µL of ethanol for mock reactions) was added and allowed to react for 10 min at 30°C. DMS probing reaction was then quenched by adding Tris pH8.0 at 400mM final.

RNAs were then reverse transcribed with the Superscript III reverse transcriptase (Invitrogen®) and NGS libraries were prepared using NEBNext Ultra II DNA Library Prep Kit (New England Biolabs®). Final products were sequenced by using the Illumina technology (NextSeq 500/500 Mid 2×150 flow cell). Sequencing data were analyzed and reactivity maps were derived using ShapeMapper2 [9]. In the end, the 306 selected sequences were probed in the following conditions:

- 30°C, without Mg^2+^ and without SAM.
- 30°C, with magnesium (Mg^2+^).
- 30°C, with magnesium and two concentrations (0.1 and 1mM) of SAM.

Each probing reaction was repeated in triplicate. The two SAM concentrations were analyzed together to improve statistics, since we found no significant effect of varying the SAM concentration in the reactivity responses of the aptamers (see Supplementary Fig. S21). The reading efficiency per site (read depths reported by Shapemapper) is plotted for the tested aptamers as grouped by primers in Supplementary Figs. S27, S28 and S29.

#### 3. Manual inspection of reactivity profiles

IPANEMAP [69] was used to generate RNA secondary structure models for each sequences. For manual inspection, we considered the reactivity of the nucleotides known to be directly involved in SAM binding (U7, G11, A46, U69, G70, U103) and of those known to be protected from shape reactivity in the closed stated, *i*.*e*., nucleotides in P1 (1-8; 101-108), in the pseudoknot (25-28; 77-80), those involved in the three base triple interactions (24, 73, 74, 76, 100). Nucleotide numbering follows the profile shown in Fig. 1C. An aptamer was considered to bind SAM if at least three of these elements are noticeably less reactive upon SAM addition, and if none of the binding determinant remain highly reactive. Note that P1 and the Pk are each considered as one element, and that some of the elements may be unreactive even in absence of SAM.

### I. Statistical analysis of reactivities

#### 1. Reactivity definition

SHAPE-MaP experiments result in measurements of sequencing error rates at each site of the RNA sequence, that correlate to the locations where the SHAPE probe has reacted with the RNA. For each site *i* = 1, …, *N* of a sequence *n*, the reactivity is defined by [73]:

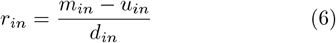

where *m*_*in*_ is the mutation rate in presence of the reagent, *u*_*in*_ is the mutation rate in its absence accounting for mutational background of the experiment, and *d*_*in*_ is the mutation rate in a denaturating condition where the RNA is expected to be unfolded, intended to cancel sequence-dependent biases. Working with *r*_*in*_ is usually better since this form should cancel site-dependent biases in the raw SHAPE mutation rates, *m*_*in*_. The basis of the SHAPE-MaP procedure relies on differences in the distribution of reactivities in base-paired and unpaired sites [73]. We have confirmed such differences are observed in our data in Fig. 7 (and also Supplementary Fig. S13).

#### 2. Statistical analysis

The finite number of sequencing reads collected at a site implies a statistical error in the reactivity computed by Eq. (6). Therefore, we cannot directly access the true reactivity *r*_*in*_ at a site, but rather an experimental measurement 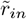 that fluctuates according to the number of reads taken at the site. To model this uncertainty, we make the simplifying assumption that the ideal reactivity of a site, *r*_*in*_, depends only on whether the site is basepaired (bp) or not (np). Under this assumption, we can write:

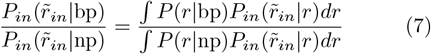

where:

- 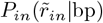 is the probability of measuring reactivity 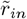at site *i* of sequence *n*, given that the site is base-paired and conditioned on the finite number of reads taken at this position.
- 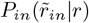 is the probability of measuring reactivity 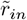 at site *i* of sequence *n*, on account of fluctuations due to a finite number of reads, conditioned on this site having a real reactivity of *r*.
- *P* (*r* | bp) is the probability distribution of reactivities of base-paired sites, at infinite read-depth, assumed to be homogeneous across sites.
- 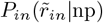 and *P* (*r* | np) are defined in a similar manner for non-paired sites.

We approximate the distributions *P* (*r* | bp) and *P* (*r* |np) by kernel density estimators fit on the corresponding empirical histograms (shown in Fig. 7A for the first experiment). The kernel function used corresponds to a standard normal, with a bandwidth set according to the Silverman rule [74]. To better estimate the histograms, we use the experimental conditions with SAM, where the secondary structure of the aptamer is expected to be more stable. We also find that these histograms can depend on the particular experiment, and therefore we fitted *P* (*r*| bp), *P* (*r*| np) for each replicate.

Applying Bayes theorem [46] in Eq. (7), we can write:

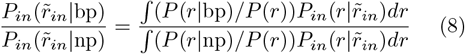

where *P* (*r*) is the histogram of real reactivities, regardless of whether a site is paired or not. The posterior 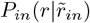 quantifies the uncertainty of the real reactivity *r* at site *i* of sequence *n*, conditioned on our information of the measurement taken at this site. This uncertainty arises from the finite sequencing reads available, which induce an experimental error in our estimate of the quantities *m, u, d* appearing in Eq. (6). Since the mutation count at a site can be modeled by a Poisson distribution [73], the posteriors of the mutation rates *m, u, d* are Gamma distributions, with a convenient choice of conjugate prior [46]. Then, we can produce a MonteCarlo estimate of 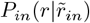 by sampling the posterior Gamma distributions of *m, u, d*, and computing the reactivity through Eq. (6). If the sampled reactivities fall predominantly far in the tails of the histograms *P* (*r* | bp) or *P* (*r* | np), respectively, the reactivity measurement is discarded as an outlier. In practice, we find that 1000 samples for each site are sufficient. These samples can then be used to approximate the numerator and denominator of the right-hand side of Eq. (8). In this way, we produce estimates of the ratios 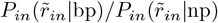, quantifying the odds that a site is paired. Supplementary Fig. S23B shows a scatter plot of reactivities in our dataset, with the standard-error estimated by the standard SHAPE-Mapper pipeline [73] (which does a firstorder error propagation through the Poisson count statistics), with each point colored according to the value of the log-odds-ratio Eq. (8). Dashed lines are approximate contours separating points that are over twice more likely to be paired (blue) or unpaired (red). The fact that these contours are not straight vertical lines indicates that, using Eq. (8), we are considering both the reactivity value and its uncertainty in assessing the plausibility that a site is paired or not. A similar approach has been proposed by [23, 79]. See also Supplementary Section I for further discussion and tests.

#### 3. Protection scores

We can exploit the likelihood ratios 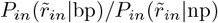 computed above to estimate the probability of the presence of a structural motif in a sequence. We define a motif of length 2*L* as a set of basepaired sites, ℳ= {*i*_1_, *j*_1_, …, *i*_*L*_, *j*_*L*_}. For example, the P1 helix motif corresponds to {1, 108, 2, 107, …, 8, 101}. We then probabilistically assess the presence or absence of the motif ℳ in molecule *n* by comparing the value of the protection score

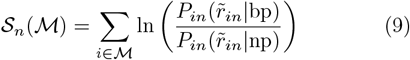

to some thresholds ±𝒮_*′*_, see Section C. This approach allows us to combine multiple reactivity measurements into a robust probabilistic measure, achieving more statistical power than when site reactivities are analyzed one by one.

This approach can be applied to SHAPE or DMS reactivity data. As DMS probing is efficient in detecting interactions involving nucleotides A or C predominantly, we only consider DMS reactivities obtained at sites where the aptamer sequence has an A or C. The base-pairing histograms *P* (*r* | bp) and *P* (*r* | np) for DMS, shown Fig. 7B, are estimated using only reactivities measured at sites with A or C nucleotides.

#### 4. Combining SHAPE and DMS data

When both SHAPE and DMS data are available for the same aptamer, we can combine them to obtain better predictions about the base-pairing status of a site. Since the SHAPE and DMS reactivities are obtained in independent experiments,

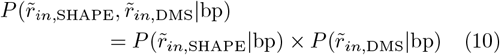

where 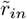 and 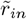 denote SHAPE and DMS reactivity data at the same site *i* of aptamer *n*. This independence implies that the log-odds ratio of the pairing status of a site or a structural motif (as in Eq. (9)), in presence of both kinds of data, can be computed by simply adding the protection scores obtained from each kind of probing alone:

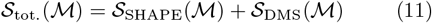

where 𝒮_SHAPE_ is the protection score obtained from SHAPE data, and 𝒮_DMS_ the protection score obtained from DMS data.

#### 5. Error bars on the rates of responsive aptamers

Given *N*_*conc*._ conclusive probed sequences, *N*_*resp*._ of which are found to be globally responsive, we estimate the response rate by 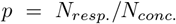. The uncertainly over *p* is, according to the binomial law, 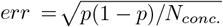. The response rates in Figs. 7, 9 are reported as (*p ± err*) *×* 100%.

We have investigated the dependence of these uncertainties on the SHAPE-Mapper read depths, which varies with the primers. Supplementary Fig. S32 shows that the inconclusive rate is strongly anti-correlated with the read depth. Both rates of responsive and non-responsive sequences increase with the read depth, a consequence of the decrease of the statistical noise. To mitigate this statistical effect, throughout this work, the response rate is computed as the ratio of responsive molecules over the number of conclusive ones, compare top and bottom panels in Supplementary Fig. S32. The dispersion due to this statistical noise are accounted for by the error bars in the results shown in Fig. 7G as explained above. We have also investigated the dependence of DMS results on the read depth (Supplementary Fig. S33). As with SHAPE, the inconclusive rate increases with the read depth.

### J. SHAPE protection scores are in agreement with consensus secondary structure

Sequence homologs in the RF00162 family are collected based on similarity to a group of manually curated sequences in the seed. Overall, for many of these sequences (both in the seed and in the full alignment), direct experimental evidence of their actual behavior and structure is limited, except for specific cases, such as the *Thermoanaerobacter tengcongensis* and the *Bacillus subtilis yitJ* SAM riboswitches, which have been extensively studied in the literature fueled by detailed knowledge of their published crystalized structures [45, 53]. For many other sequences in the MSA, their actual behavior is at most hypothesized based on indirect evidence.

We have here obtained detailed SHAPE data of *B*_seed_ = 151 sequences of the seed alignment. Our data shows that, in average, these sequences are compatible with the consensus secondary structure of the RF00162 family, shown in Fig. 1B. Indeed, we have computed the average protection scores ⟨𝒮 (*i*) ⟩ for each site *i*, over the sequences in the seed alignment probed in our experiments,

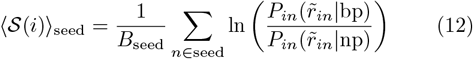

Figure 10B plots ⟨𝒮 (*i*) ⟩in the conditions with SAM and without SAM. Overall, the averaged protection scores are in detailed agreement with the consensus secondary structure of the aptamer, depicted in Fig. 10A. Helices P2, P3, P4 are seen to be base-paired in average in all conditions, with a mild overall increase in the values of 𝒮 with the addition of magnesium and then SAM, indicating overal structural stabilization. The central junction loop (CL), and the loops on the second helix L2, the third helix L3, and the fourth helix L4, are consistently measured as reactive when SAM is not present, indicating that these sites are unpaired, as expected. Besides these major structural motifs, we also appreciate finer details such as the reactivity of single isolated bulge sites in positions 46 and 65 in absence of SAM. Next, comparing the behavior across different conditions, we appreciate the effect of magnesium and SAM on the structure. We highlight (in green) sites that change significantly in response to SAM. These include sites in direct contact with SAM (as known from the crystal structure [53]), and other tertiary motifs known to form in response to SAM. We discuss these next.

**FIG. 10.**
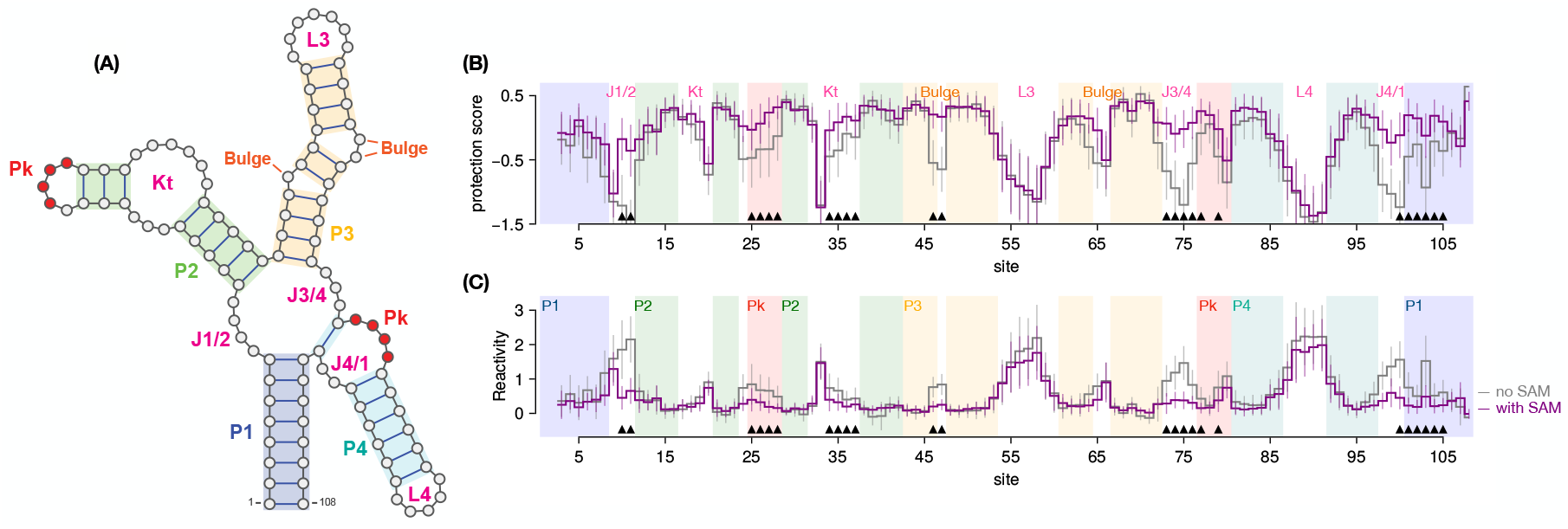
**A)** Annotated consensus secondary structure of the aptamer domain of the SAM-I riboswitch family (Rfam ID RF00162). **B)** Average protection scores, ⟨𝒮(*i*)⟩ (see Eq. (12)) per site, of the natural probed sequences, for the two conditions: with SAM and no SAM. Error bars (standard deviation) are also shown. Both statistics are computed over the *B*_seed_ = 151 probed sequences in the seed alignment. Hallmark sites (Supplementary Table S2) are indicated with black triangles. **C)** Average site reactivities with error bars (standard deviation).

### K. Selection of Hallmark sites

We selected 24 hallmark sites across the aptamer sequence, for which we could rationalize observed reactivity changes in response to SAM binding, and which are consistent with expectations from previous chemical probing studies on SAM-I riboswitches and previous structural data. These sites also exhibit significant reactivity responses across natural sequences in our data, see Fig. 10. They are listed in Supplementary Table S2. In Supplementary Section Q we include further discussion and references to several previous literature reports justifying the choices of each of these sites.

Our results are robust to minor variations in the selection of Hallmark sites used to evaluate the response of aptamers to SAM. For example, although we could not find previous reports of reactivity responses in J4/1, we find in some cases that sites 98 and 99 exhibit protection upon SAM binding (see Fig. 10). We tried adding few selected sites (such as 98, 99), or excluding some, and confirmed that our main results (such as numbers of responsive sequences) remain unchanged. Additional results are reported in Supplementary Section Q.

### L. Principal component analysis

We carried out a principal component analysis (PCA) of the natural MSA. First, we one-hot encode the natural sequences in a *q × N × B* binary tensor 𝒟, where *B* = 6161 is the number of sequences in the full MSA collected above. The tensor has 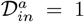 if sequence *n* of the alignment has symbol *a* ∈ {1, …, 5} at position *i*, and otherwise 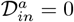. We then compute a covariance tensor, defined as follows

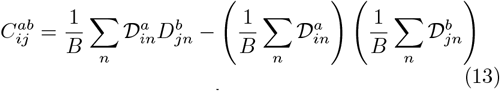

We flatten the tensor 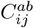 into a *qN × qN* matrix, and then perform a standard eigenvalue decomposition on it. Individual sequences are then projected along the two top components (with largest eigenvalue) of the decomposition.

## M. Data availability

Sequencing data and processed reactivity has been deposited to the Gene Expression Omnibus (GEO) database, under the accession GSE266263 [https://www.ncbi.nlm.nih.gov/geo/query/acc.cgi?acc=GSE266263]. All processed data and processing code is available on the accompanying Github repository [15] (see Code Availability).

## N. Code availability

The code used to develop the model, perform the analyses and generate results in this study is publicly available and has been deposited in Github at https://github.com/cossio/SamApp2025.jl, under MIT license. The specific version of the code associated with this publication is archived in Zenodo and is accessible via https://doi.org/10.5281/zenodo.17232573 [15]. The main repository (https://github.com/cossio/SamApp2025.jl) is provided as an opensource Julia [5, 16] package. We also provide an implementation of RBM in Python at https://github.com/cossio/SamApp2024Py and an example Google Colab notebook at https://colab.research.google.com/drive/1nOfFLWCwLy7a0aZ52cFHKUfF7erAMp5f?usp=sharing.

## Acknowledgements

We are grateful to Sean R. Eddy and Eric P. Nawrocki, for helpful discussions about Infernal. This work is principally supported by ANR Decrypted 19-CE30-0021-01. Additional fundings: PSL AI Junior Fellow program (JFdCD); ANR 19-CE30-0021-03, ANR 20-CE12-0026-02, ANR 19-CE45-0023-02,ANR 21-CE45-0034-03 (PH and FXLdM).

## Author contributions

S.C., R.M., J.F.d.C, B.S. designed the work, interpreted the data and wrote the paper. S.C., R.M., J.F.d.C designed new methods to analyse the data and revised the work. P.H., F.X.L.d.M., B.S. performed the experiments and analysed the data. Y.P., B.M., A.d.G. contributed to the data analysis.

