## Supplementary Materials for "Designing Molecular RNA Switches with Restricted Boltzmann Machines"

### SUPPLEMENTARY INFORMATION

Jorge Fernandez-de-Cossio-Diaz, Pierre Hardouin, Francois-Xavier Lyonnet du Moutier, Andrea Di Gioacchino, Bertrand Marchand, Yann Ponty, Bruno Sargueil, Rémi Monasson, Simona Cocco

### Appendix A: Further details on RBM implementation

#### 1. Training the RBM

The likelihood assigned by the RBM to a sequence depends on all the parameters of the model: the $q \times N \times M$ weights tensor $w_{i\mu}(v_i)$, the $4M$ hidden unit dReLU parameters $\gamma_\mu^\pm, \theta_\mu^\pm$, and the $q \times N$ visible unit fields $\mathcal{V}_i(v_i)$. See Eq. 2 in the main text. Given a set of aligned data sequences (multiple-sequence alignment, or MSA), these parameters are learned, in an unsupervised way, by maximizing the average log-likelihood of the data

$$\mathcal{L} = \frac{1}{B_{\text{MSA}}} \sum_{\mathbf{v} \in \text{MSA}} \ln P(\mathbf{v}) \,, \tag{S1}$$

(where the sum is taken over the $B_{\text{MSA}}$ sequences in the MSA) plus a regularization term,

$$\mathcal{R} = -\frac{\lambda_{\text{reg}}}{2} \sum_{\mu=1}^{M} \left( \sum_{i=1}^{N} \sum_{v=1}^{q} |w_{i\mu}(v)| \right)^2 \,, \tag{S2}$$

(where $\lambda_{\text{reg}}$ is a non-negative parameter). This form of the regularization, combining $L_2$ and $L_1$ norms, has been proposed by [1] to favor sparse weights with balanced norms across all hidden units. Regularization helps avoid over-fitting and promotes a smoother training and sampling of the model [2].

To train the model, we perform a variation of gradient ascent over $\mathcal{L} + \mathcal{R}$. Schematically, if $\omega_t$ denotes a parameter of the RBM (weights $w_{i\mu}$ or a parameter of the potentials $\mathcal{V}_i$, $\mathcal{U}_\mu$), at time $t$ of the training, then for the next training iteration the parameters are updated as follows:

$$\omega_{t+1} = \omega_t + \eta \frac{\partial}{\partial \omega} (\mathcal{L} + \mathcal{R}) \,, \tag{S3}$$

where $\eta$ is a small positive learning rate.

In practice, the optimization is accelerated by an adaptive momentum term [3] and a centering trick [4]. More specifically, we use the ADAM [3] optimization algorithm, with minibatches containing 128 sequences and for a total of 10000 gradient update steps.

Computing the gradient of $\mathcal{L}$ requires to estimate the moments of visible and/or hidden variables with respect to the model distribution [2]. We employ the Persistent Contrastive Divergence (PCD) algorithm [5], where a number of Markov chains sampled from the model are updated in each parameter update. We have that:

$$\frac{\partial \mathcal{L}}{\partial \omega} = \underbrace{\left\langle \frac{\partial(-E_{\text{eff}}(\mathbf{v}))}{\partial \omega} \right\rangle_{\text{MSA}}}_{\text{positive gradient}} - \underbrace{\left\langle \frac{\partial(-E_{\text{eff}}(\mathbf{v}))}{\partial \omega} \right\rangle_{\text{RBM}}}_{\text{negative gradient}} . \tag{S4}$$

The first term is an empirical average performed over the data, as in Eq. (S1), while the second term is averaged over sequences sampled from the RBM. We represent these terms as arrows in Fig. 2B. The first term (blue), tends to drive the parameters $\omega$ of the RBM such that the score $-E_{\text{eff}}(\mathbf{v})$ of sequences $\mathbf{v}$ in the data is increased. This results in the model assigning higher probabilities to regions of sequence space densely populated by data. To do this, the hidden units of the RBM must extract features shared by the data sequences and thus likely to be important for their biological function. Conservation of probability implies that regions of sequence space not populated by data sequences must be penalized. This is taken care of by the second term in Eq. (S4) (in red), which tends to increase the average energy of sequences uniformly sampled from the RBM. The net effect of the two terms (also called positive and negative phase terms in earlier papers [2, 6]), is that the RBM places most probability mass in regions densely populated by data and low probability elsewhere. However, the finite parameterization and discovered features usually extrapolate also to novel regions in sequence space, not covered by the data, where the model assigns high probability, as illustrated in green in Fig. 2B. The trained RBM model automatically extracts features and constraints from the data, which are then imposed in the generated sequences, in a manner akin to the features used for *positive* and *negative design*, in rational design approaches.

In our implementation of Persistent Contrastive Divergence (PCD) [5], we take 100 Monte Carlo steps to update the Markov chains in each iteration. The number of Monte Carlo chains equals 128. During training, the pseudolikelihood of the data is monitored to assess convergence. In addition, hidden units are re-scaled to a variance of 1 at each iteration by a simple scaling transform. The implementation is identical to the one used in [7], except for the implementation of a *standardization* trick (generalizing the centering trick of [4]), that we describe in the next section.

### 2. Standardized RBMs

We generalize the centering trick of [4] to include a normalization of the hidden unit variances. As has been shown previously [4], such centering and standardization of unit activities leads to improved training convergence rates and stability.

More precisely, we can define the standardized Restricted Boltzmann machine (stdRBM) energy function as follows:

$$E(\mathbf{v},\mathbf{h}) = \cdots - \sum_i \theta_i v_i - \sum_\mu \theta_\mu h_\mu - \sum_{i\mu} w_{i\mu} \frac{v_i - \lambda_i}{\sigma_i} \frac{h_\mu - \lambda_\mu}{\sigma_\mu} + \sum_{i\mu} \frac{w_{i\mu}}{\sigma_i \sigma_\mu} \lambda_i \lambda_\mu \tag{S5}$$

where we only show the fields for the unit potentials for simplicity. The distribution over configurations of visible and hidden units by the usual relations:

$$P(\mathbf{v},\mathbf{h}) = \frac{1}{Z} e^{-E(\mathbf{v},\mathbf{h})}, \qquad Z = \operatorname*{tr}_{\mathbf{v},\mathbf{h}} e^{-E(\mathbf{v},\mathbf{h})} \tag{S6}$$

The parameters $\lambda_i, \lambda_\mu$ and $\sigma_i, \sigma_\mu$ are meant to track the mean and standard deviations of the units, respectively.

Note that:

$$E(\mathbf{v},\mathbf{h}) = -\sum_i \left\{ \theta_i - \sum_\mu \frac{w_{i\mu}}{\sigma_i \sigma_\mu} \lambda_\mu \right\} v_i - \sum_\mu \left\{ \theta_\mu - \sum_i \frac{w_{i\mu}}{\sigma_i \sigma_\mu} \lambda_i \right\} h_\mu - \sum_{i\mu} \frac{w_{i\mu}}{\sigma_i \sigma_\mu} v_i h_\mu \tag{S7}$$

If we have a new set of parameters $\theta', w', \lambda', \sigma'$, satisfying,

$$\begin{aligned} \theta_i - \sum_\mu \frac{w_{i\mu}}{\sigma_i \sigma_\mu} \lambda_\mu &= \theta'_i - \sum_\mu \frac{w'_{i\mu} \lambda'_\mu}{\sigma'_i \sigma'_\mu}, \\ \theta_\mu - \sum_i \frac{w_{i\mu}}{\sigma_i \sigma_\mu} \lambda_i &= \theta'_\mu - \sum_i \frac{w'_{i\mu}}{\sigma'_i \sigma'_\mu} \lambda'_i \\ \frac{w_{i\mu}}{\sigma_i \sigma_\mu} &= \frac{w'_{i\mu}}{\sigma'_i \sigma'_\mu} \end{aligned} \tag{S8}$$

then, we have, $E'(\mathbf{v},\mathbf{h}) = E(\mathbf{v},\mathbf{h})$. As a consequence the distributions over unit activities remain invariant, $P'(\mathbf{v},\mathbf{h}) = P(\mathbf{v},\mathbf{h})$. Therefore, the two models are equivalent.

In particular, a standardized RBM with parameters $\theta, w, \lambda, \sigma$ is equivalent to an ordinary RBM with parameters $\theta', w'$, $\lambda' = 0$, $\sigma' = 1$ provided that:

$$\theta'_i = \theta_i - \sum_\mu \frac{w_{i\mu}}{\sigma_i \sigma_\mu} \lambda_\mu, \quad \theta'_\mu = \theta_\mu - \sum_i \frac{w_{i\mu}}{\sigma_i \sigma_\mu} \lambda_i, \quad w'_{i\mu} = \frac{w_{i\mu}}{\sigma_i \sigma_\mu} \tag{S9}$$

During training, we dynamically update the parameters $\lambda_i, \lambda_\mu$ to follow the mean activities of the visible and hidden units, while $\sigma_i, \sigma_\mu$ are dynamically tracking the standard deviations of the unit activities. If the hidden units are continuous and have location and scale parameters, we can rescale their activities after each training step so that their estimated means and variances are set to zero and one, respectively. This is compensated by a rescaling of the weights $w_{i\mu}$ attached to the hidden unit, so that the resulting RBM model is equivalent after this transformation.

In agreement with the intuition set forth in [4], we find empirically that training our RBM as an equivalent stdRBM, leads to faster and more stable training.

#### 3. Sampling the RBM

Having trained the model, sampling can be performed through a Monte Carlo procedure known as Gibbs sampling [2]. It exploits the two-layer RBM architecture, by noting that the conditional distributions of one layer given the configuration of the other layer, factorize:

$$\begin{aligned} P(\mathbf{v}|\mathbf{h}) &\propto \prod_{i=1}^{N} \exp\left(-\mathcal{V}_i(v_i) + \sum_{\mu=1}^{M} w_{i\mu}(v_i) h_\mu\right) \\ P(\mathbf{h}|\mathbf{v}) &\propto \prod_{\mu=1}^{M} \exp\left(-\mathcal{U}_\mu(h_\mu) + \sum_{i=1}^{N} w_{i\mu}(v_i) h_\mu\right) \end{aligned} \tag{S10}$$

These conditional distributions are therefore easy to sample. The Gibbs sampling algorithm consists of iteratively sampling one layer conditioned on the other layer, and vice-versa, for a number of steps, and collecting the configuration at the final iteration. If a large enough number of steps are taken, the resulting sample is guaranteed to be a good approximation of an equilibrium sample of the RBM. Equilibration can be assessed by inspecting convergence of quantities such as the average energies of the samples. For the RBM we trained in this work, we found that $\sim 10000$ Gibbs sampling steps were more than sufficient to reach a stable plateau. For more implementation details we refer to [7].

#### 4. RBM trained on the SAM-I riboswitch aptamer domain sequence family

For the RF00162 sequence family of the SAM-I riboswitch aptamer domain (downloaded from Rfam [8]), we trained an RBM with $N = 108$ Potts visible units corresponding to the aligned sequence sites, together with $M = 100$ hidden dReLU units, with a regularization weight of $\lambda_{\text{reg}} = 0.01$.

#### 5. Cross-validation analysis

The RBM employed in the main-text has $M = 100$ dReLU hidden units and was trained with a regularization penalty of $\lambda_{\text{reg}} = 0.01$. After extensive exploration of other numbers of hidden units and regularization penalties, we chose these settings as a good balance between model quality (as measured by the log-likelihood of withheld data) and simplicity. Figure S1 shows the results of the experiments performed at different regularization penalties (panel A) and numbers of hidden units (panel B). Beyond 100 hidden units there are diminishing returns in the validation log-likelihood.

We have conducted cross-validation tests of our model to control for overfitting. We trained new RBMs on reduced datasets, consisting of 80% randomly selected sequences of the MSA for training, and 20% withheld for validation. We then computed the RBM scores ($-E_{\text{eff}}$) of sequences used in training and validation, and compared their histograms in S1C. Both histograms are in close agreement, indicating that our hyper-parametric choices place the model far from an overfitted regime. To further justify our architectural choices in the RBM, we experimented various choices for the number of hidden units of the RBM and the regularization strength $\lambda_{\text{reg}}$. Regarding the regularization strength, we find that for $\lambda_{\text{reg}}$ larger than 0.05, we suffer a loss of log-likelihood of the validation data. On the other hand, for $\lambda_{\text{reg}}$ smaller than 0.01 there is no significant gain in likelihood. The results of these experiments are shown in Figure S1A. We have then set $\lambda_{\text{reg}} = 0.01$, since a non-zero regularization makes the model more robust in case of very conserved sites, and makes the model easier to sample and train [1, 2]. The hyper-parameter scan also reveals diminishing returns for more than 100 hidden units in terms of the log-likelihood of validation data, as shown in Figure S1B. Therefore, we have chosen to use 100 hidden units in this work.

#### 6. Computation of epistatic scores

The riboswitch aptamer structural fold imposes contacts between distant sites along the RNA sequence, which are reflected in covariations between nucleotides in the corresponding columns of the MSA. To assess how well the RBM

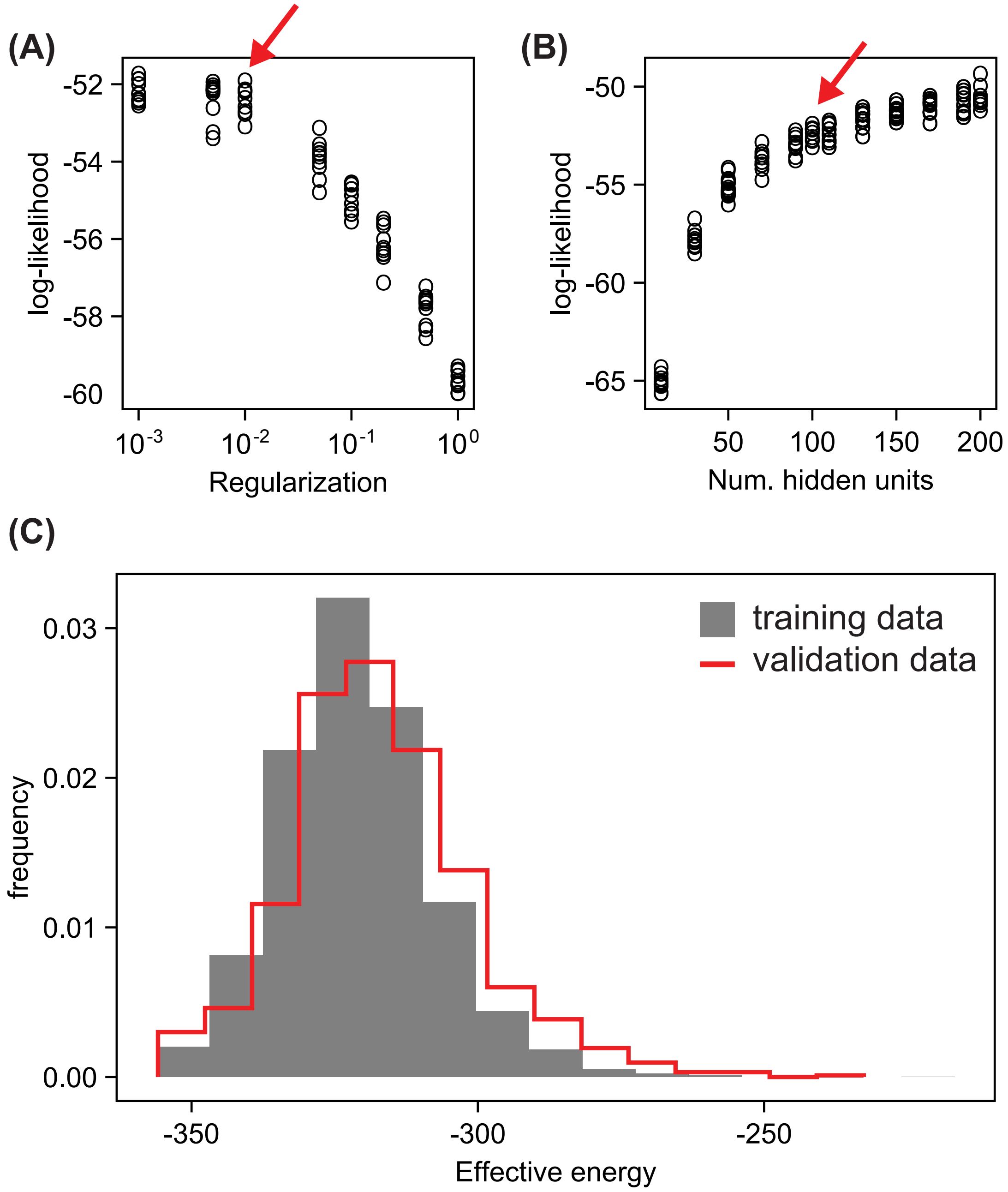


FIG. S1. Validation of RBM architectural choices. **(A)** RBMs were trained with different regularization weights, and **(B)** different numbers of hidden units (from 20 to 200), on different random subsamples containing 80% of the training data. The plots show the log-likelihood of the remaining 20% of validation data. Red arrows in D,E mark our final choices for the regularization ($\lambda_{\text{reg}} = 0.01$) and number of hidden units ($M = 100$). **(C)** An RBM with 100 hidden units and $\lambda_{\text{reg}} = 0.01$, trained on a randomly selected subset containing 80% of training sequences was used to compute the effective energies of sequences in its training set and the remaining 20% of sequences in the validation set. Histograms of the effective energies ($E_{\text{eff}}$ from (5) in the main text) of the two groups (training data in gray, validation data in red) of sequences shows excellent agreement, suggesting no overfitting has occurred.

102 captures sequence features connected to these structural constraints, we compute the following epistatic score [1]:

$$\mathcal{J}_{ij} = \sum_{a,b} \left\langle \frac{1}{25} \sum_{a',b'} \ln \left[ \frac{P(\mathbf{v}_{ij}^{a,b}) P(\mathbf{v}_{ij}^{a',b'})}{P(\mathbf{v}_{ij}^{a',b}) P(\mathbf{v}_{ij}^{a,b'})} \right] \right\rangle_{\text{MSA}}^{2} \tag{S11}$$

for pairs of sites $i, j$ along the sequence. Here, $\mathbf{v}_{ij}^{a,b}$ denotes a sequence $\mathbf{v}$ from the MSA, which has suffered a double mutation: site $i$ was modified to symbol $a$, and site $j$ was modified to symbol $b$. $P(\mathbf{v}_{ij}^{a,b})$ denotes the likelihood, Eq. (4), of this modified sequence, and the average $\langle \dots \rangle_{\text{MSA}}$ is taken over all sequences of the MSA. Note that we average over all possible pair of mutations $a', b'$ at sites $i, j$, dividing by $25 = 5^2$, the number of possible letters (4 nucleotides and a gap symbol) at these two positions. This score, introduced by [1], is closely related to the Frobenius norm of interactions used in Direct-Coupling analysis for contact prediction in proteins [9], and measures how the epistatic effect of a pair of mutations is enhanced in comparison to the effects of the single mutations by themselves. In addition, we apply the average-product correction (APC) to the matrix $\mathcal{J}_{ij}$, which has been argued to decrease the impact of phylogenetic biases in contact prediction [9, 10]. The APC corrected contact matrix, $\tilde{\mathcal{J}}_{ij}$, is defined by:

$$\tilde{\mathcal{J}}_{ij} = \mathcal{J}_{ij} - \frac{\sum_{kl} \mathcal{J}_{kj} \mathcal{J}_{il}}{\sum_{kl} \mathcal{J}_{kl}} \tag{S12}$$

### Appendix B: RBM reproduces statistics of natural sequences

The quality of the model fit after training, and the quality of convergence, can be assessed by comparing statistics of sampled sequences against the empirical statistics of the MSA. In Figure S2A, we compare the single-site statistics, computed as the frequency of occurrences of each nucleotide (or gap symbol) at each position of the alignment. The agreement is excellent (Pearson correlation = 0.98), indicating that the RBM reproduces the conservation of important sites (cf. Figure 1C). Furthermore, RBM sampled sequences reproduce the covariance of pairs of sites of the natural sequences, as shown in Figure S2D. In this case, we compute the deviation of the frequencies of co-occurring nucleotides at pairs of sites from the expectation arising from their independent conservations. Such joint covariations arise from interactions across the sequence, related for example to secondary or tertiary contacts, or other functional constraints. The agreement is also excellent (Pearson correlation = 0.97), indicating that the RBM is able to reproduce the covariation of the natural MSA. We also evaluated the RBM scores of sampled sequences compared to the energies assigned by the RBM to the natural sequences. As we show in Figure S2C, the two histograms are in close agreement to each other.

To evaluate the diversity of a set of sequences, natural or generated, we compute the matrix of all possible pairwise Hamming distances between pairs of distinct sequences, where the Hamming distance is defined as the number of positions where the two sequences differ. Figure S2E shows the histogram of these pairwise distances for the natural sequences in gray. Typically, two randomly selected natural sequences differ in about 40% of sites, or 43 out of the 108 aligned sites. We then sampled 5000 sequences from the RBM, and computed the histogram of their pairwise distances (between themselves). We plot the result in Figure S2E in red. We see that the histogram closely resembles the histogram of the natural sequences. We conclude that the RBM generated sequences recapitulate the natural diversity of the sequence homologs family. Furthermore, the RBM generates novel sequences, not seen in the data. Indeed, Figure S2F shows the histogram of distances between each sampled sequence, and the closest natural sequence to it. Typical RBM samples differ in 20 sites from the closest sequence in the MSA, and therefore constitutes a truly novel sequence.

Finally, we observed a strong variation in sequence lengths in natural sequences. In particular, dramatic variations of the P4 helix have been reported in the literature [11, 12], where riboswitches without P4 have been shown to be functional although with lower affinities to SAM. Although our RBM is not able to model insertions, it is still able to emit sequences of varying lengths by having more or less gaps in the sequence. We therefore compared the distribution of sequence lengths generated by the RBM, with the histogram of natural sequence lengths (not considering inserts) from the MSA. The plot in Figure S2B confirms that the RBM reproduces the correct length statistics.

Overall, these results suggest that the RBM is able to reproduce accurately several statistical features of the natural sequences.

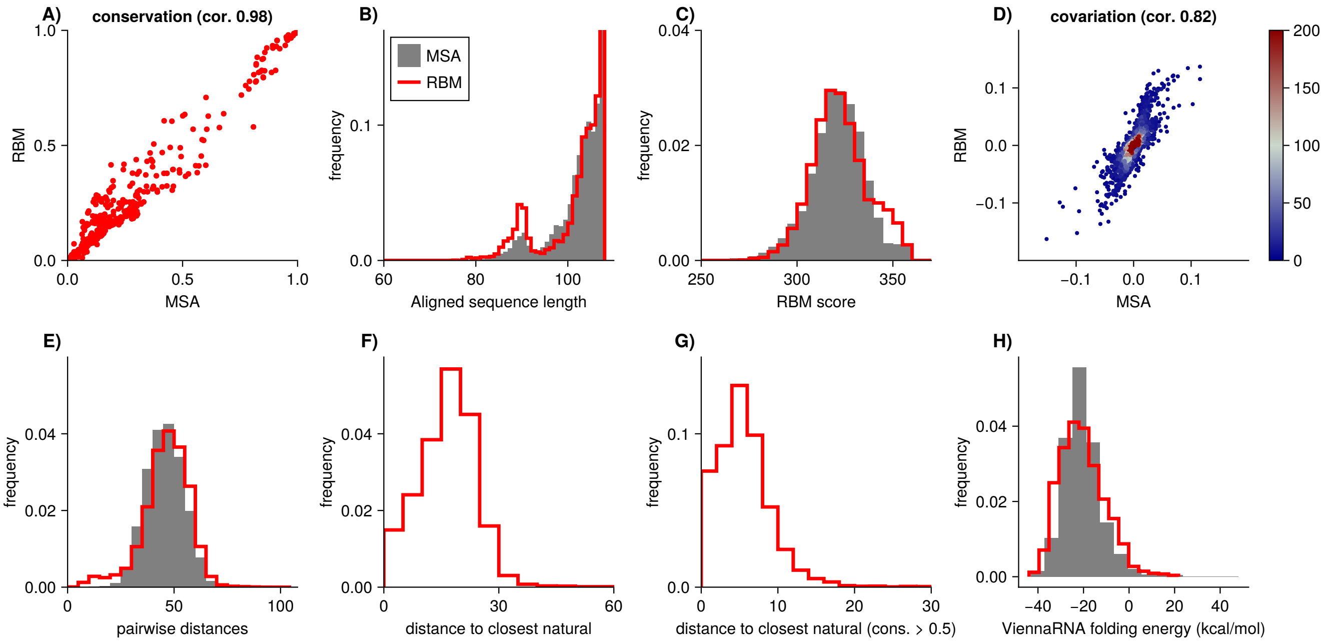


FIG. S2. RBM generates novel and diverse sequences that recapitulate statistics of natural homologues. **(A)** RBM samples reproduce single-site nucleotidic conservation of the natural sequences (Pearson correlation = 0.98). **(B)** Histogram of natural sequence lengths (gray) and of RBM generated sequences (red). Note that insertions are discarded. Sequence length is defined as the number of aligned sites that are not gaps (deletions). **(C)** Histograms of RBM scores (-$E_{\text{eff}}$ in Eq. (5) of the main text) of natural sequences (gray) and of RBM samples (red). **(D)** RBM samples reproduce the statistics of nucleotidic covariation of natural sequences (Pearson correlation = 0.82). Since the number of paired sites is very large, the points are colored by their density in the plot, according to the color bar legend. **(E)** Histograms of pairwise Hamming distances, among natural sequences (gray) and among RBM samples (red). **(F)** Histogram of Hamming distances, from each RBM sampled sequence, to its closest natural sequence. **(G)** Like (F), but considering only sites with a conservation > 0.5. **(H)** Histogram of folding energies (into the consensus secondary structure of the RF00162 Rfam family) of natural sequences (gray) and RBM samples (red), estimated with the ViennaRNA package [13].

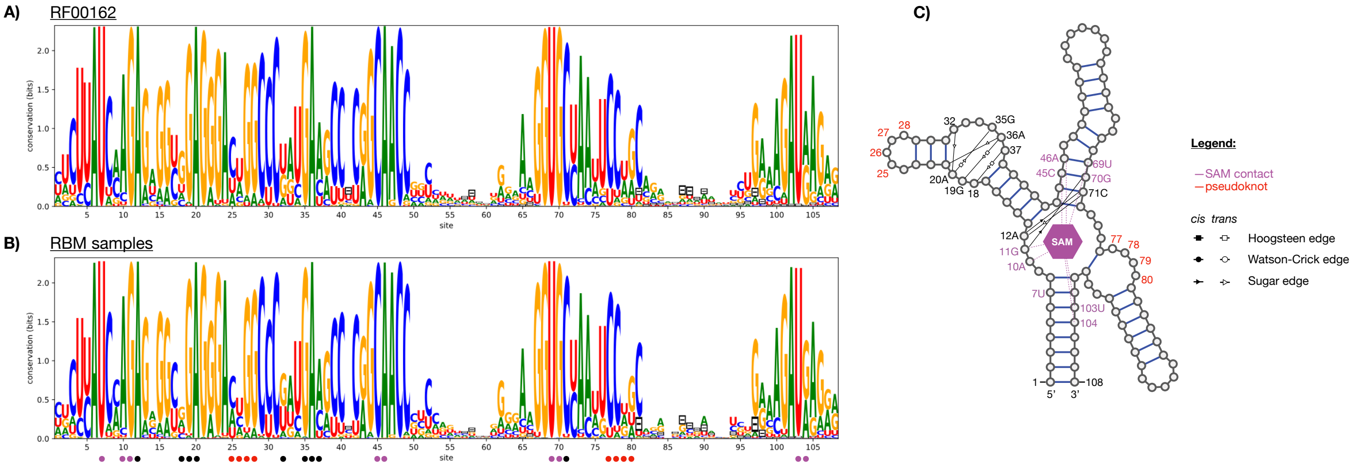


FIG. S3. Conservation and coevolution in the RF00162 SAM-riboswitch family can be explained by structural alignment. **A)** Sequence logo of the natural sequences in the RF00162 family. The strong conservation and covariation of several sites or pairs in contact (indicated below by dots of different colors) is related to structural and functional constraints in the family. **B)** Sequence logo of sequences sampled from the RBM. The RBM reproduces closely the conservation profile of the natural sequences. **C)** Interaction network diagram of the SAM riboswitch natural aptamer, based on the 2GIS PDB crystal from T. tengcongensis. The sequence numbering is based on the alignment to the RF00162 family consensus. The annotations of secondary structure and tertiary contacts is based on [14] (see in particular Fig. 6B of the cited work). We highlight sites involved in direct contact with SAM (in purple), pseudoknot (red), or other tertiary contacts (black), which in turn exhibit strong conservation and coevolution signals in A,B. Tertiary contacts are classified as in [14]. For example, sites 18-20 and 35-37 interact through *trans* Hoogsteen/Sugar-edges, which are associated to A/G base-pairs, explaining the conservation of these nucleotides at these positions.

#### 1. Distance of RBM generated aptamers to closest natural sequences

The RBM is capable of sampling novel aptamer sequences, distinct from any natural sequence. An example of this is shown below, in Fig. S2E, which shows that typical RBM samples are between 20 and 40 mutations away from the closest natural sequence (out of 108 sites in total). Figure S4 plots the Hamming distance to the closest natural sequence ($x$-axis) vs. the RBM score of the probed RBM generated sequences in the first experiment. We find functional sequences up to 30 mutations away from the closest natural sequence.

In the right panel in Figure S4, the Hamming distance to natural sequences is computed only considering conserved sites. To define these sites, we have computed the conservation score:

$$S_{\text{cons}}(i) = \log_2(5) + \sum_a P_{\text{MSA}}(a,i) \log_2 P_{\text{MSA}}(a,i) \tag{S1}$$

for each site, where $P_{\text{MSA}}(a,i)$ is the probability of observing letter $a \in \{A,C,G,U,-\}$ at position $i \in \{1,2,\ldots,108\}$ in the sequences of the Rfam MSA. Then the 76 conserved sites are those for which $S_{\text{cons}}(i) > 0.5$.

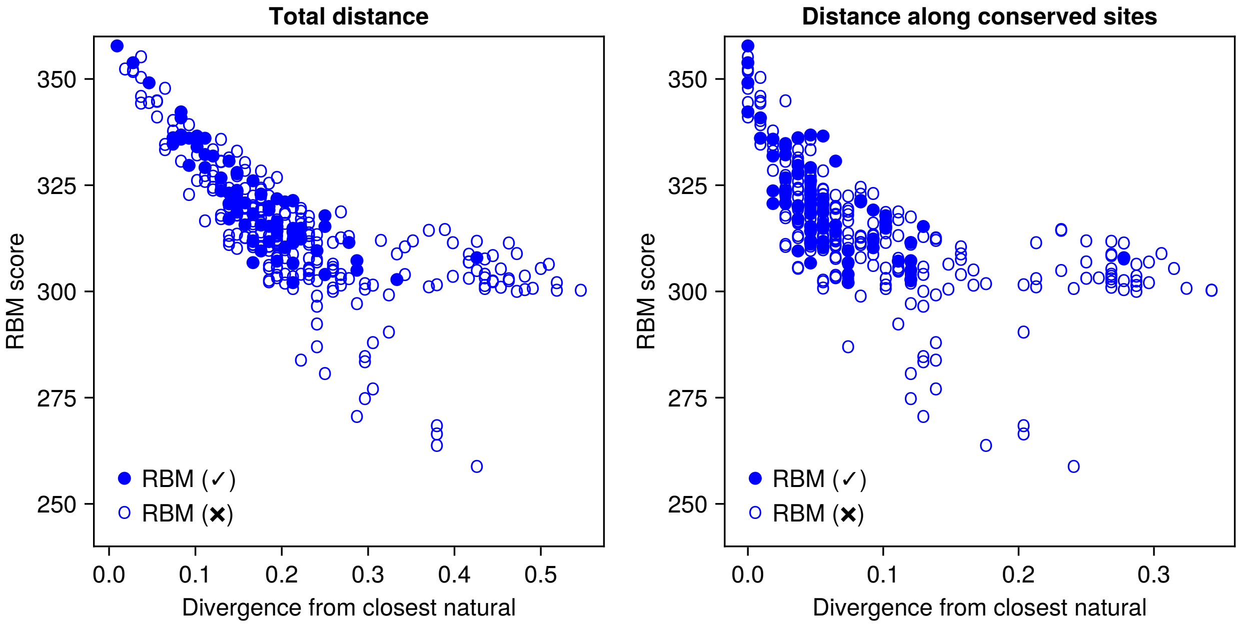


FIG. S4. Distance to closest natural sequence of RBM generated probed sequences (x-axis) vs. RBM score (y-axis). Filled blue points correspond to sequences that respond to SAM, according to the definition in the main text (see Methods). Empty blue points denote sequences that do not respond to SAM. In the left panel a standard Hamming distance across all 108 sites is computed. In the right panel, the Hamming distance is computed considering only the 76 sites that have a conservation score of $> 0.5$ in the RF00162 Rfam family.

### Appendix C: Principal components analysis of the natural RF00162 MSA

We performed a principal component analysis (PCA) on the sequences of the full RF00162 multiple-sequence alignment (MSA), as explained in the main text (Methods). Figure S5 shows the top two components, in sequence logo representation. The first component notably reflects the deletion of the P4 helix in a cluster of natural sequences, mostly Actinomycetota. This is appreciated from the prominence of gaps in sites from 81 to 99. See also Fig. 4B in the main text. Thus the top component separates sequences in which P4 is present (and thus have a positive projection onto this component) from sequences in which P4 is deleted (having a negative projection onto this component). Various reports in the literature have discussed the role of P4 in the function of the riboswitch. In particular, [12] found sequences without P4 able to bind SAM, and were able to correlate experimentally the length of P4 with SAM affinity.

Figure S6 shows the projection of the sequences probed in the first experiment along the two top principal components (PC), annotated by our protection score as switcher (filled circles) or non-switcher (empty circles). As can be seen in the figure, probed sequences span the space defined by these two PCs, suggesting adequate coverage of the diversity represented in the MSA.

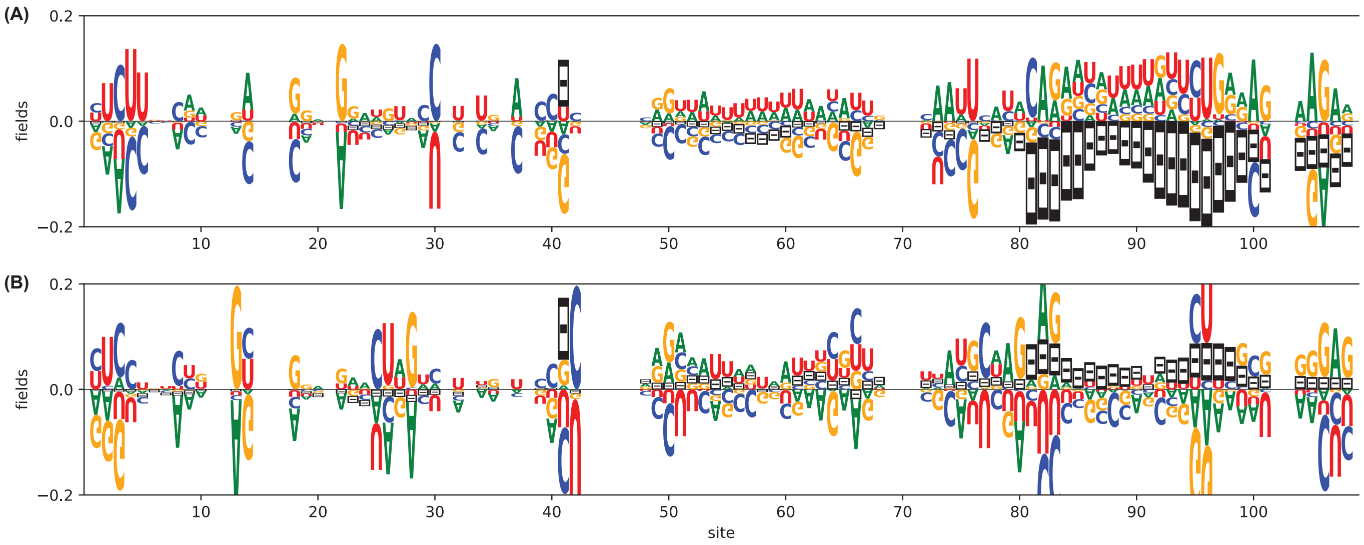


FIG. S5. **(A)** Sequence logo of the eigenvector corresponding to the largest eigenvalue of the correlation matrix of the RF00162 MSA. **(B)** Sequence logo of the eigenvector corresponding to the second largest eigenvalue of the correlation matrix of the RF00162 MSA.

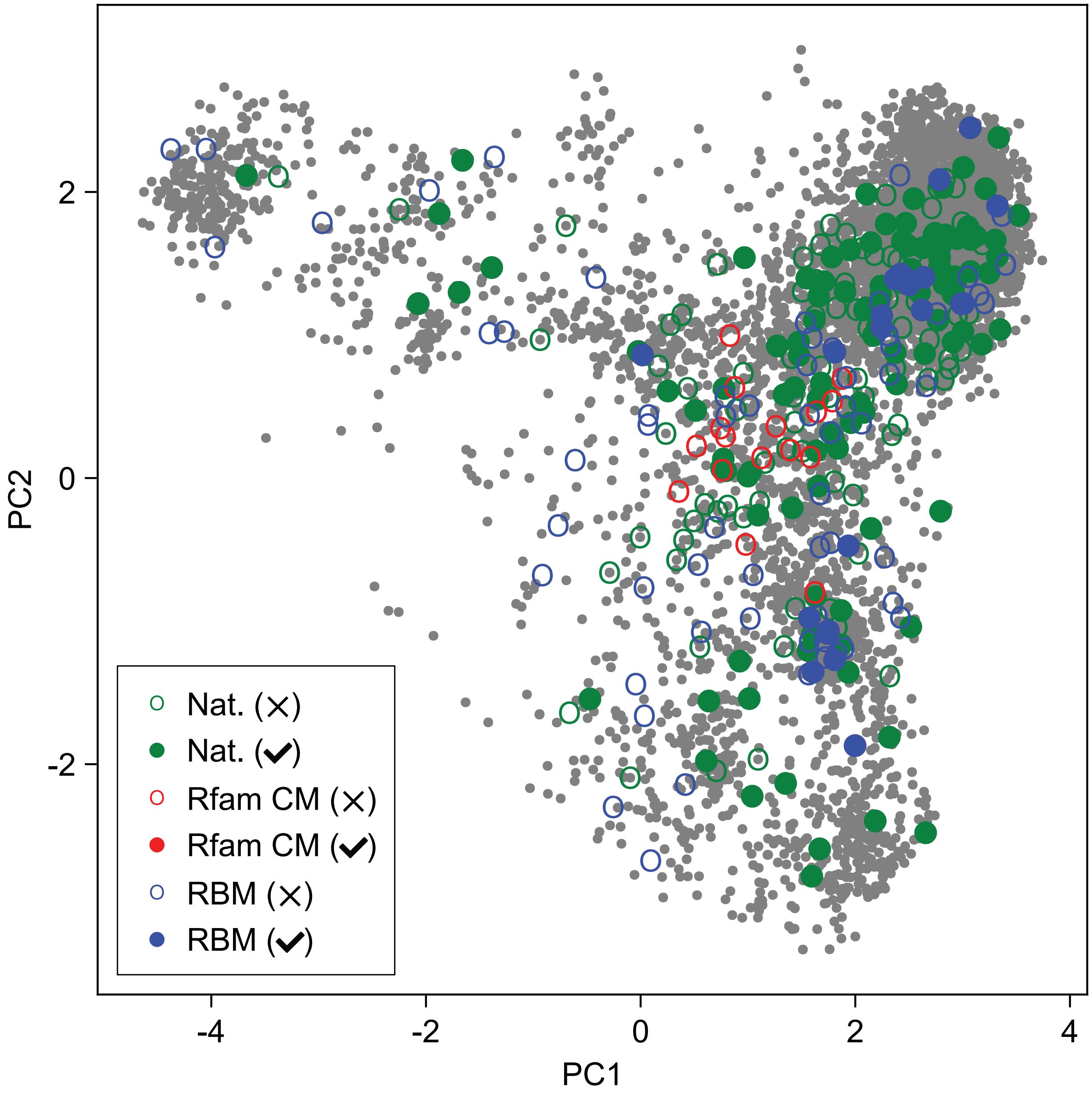


FIG. S6. Projection of probed sequences in the first batch onto principal components of the RF00162 MSA. Sequences are colored by their origin: Natural (green), Rfam CM (red), and RBM (blue). Sequences that respond to SAM are shown as filled disks, whereas unresponsive sequences are shown as empty circles.

### Appendix D: Comparison to RfamGen

While this work was a preprint on the Biorxiv, a recent paper [15] proposed a variational autoencoder (VAE) architecture, that samples transitions in a covariance model grammar to design novel RNA sequences. In comparison, we believe that RBM are a much simpler approach and yields similar results.

In SI Figure S7, we have computed the RBM score of RfamGen sampled sequences for the SAM-I riboswitch family RF00162.

Note that RfamGen offers two modes for generating novel sequences:

- The traversal across the CM grammar is stochastic, which yields more diverse (but possibly noisy) sequences. In this mode, the CM-VAE should approximate statistics of natural sequences it was trained on. This is similar to how RBM sequences are sampled.

- The traversal of the CM grammar is deterministic (after conditioning on the VAE output), by selecting for each step the most likely transition up to that point. This limits the diversity of sampled sequences but increases the chances of generating better sequences. However, since this differs from how the CM-VAE was trained, in this mode the of statistics sampled sequences are likely heavily distorted in comparison to their natural counterparts. This is analogous (though not precisely) to sampling from the RBM but at a lower temperature.

Since in this work we have only sampled RBM sequences at temperature one, and we have tried to match the statistics of natural sequences, we focus on the first generation mode of RfamGen.

In SI Figure S7A we compute the RBM score of RfamGen samples, in comparison to the natural sequences. We find that RfamGen sequences have lower RBM scores than natural sequences. In contrast, the Infernal score (computed with the Rfam CM) of RfamGen sequences closely matches the statistics of the natural sequences (SI Figure S7C).

Lastly, we would like to evaluate the likelihood of RBM designed sequences in the RfamGen model. Since evaluating the marginal likelihood of VAEs is intractable, we decided to map all samples to the VAE latent space, where the probability is simply Gaussian. We see in Fig. S7B that MSA and RBM sequences have similar latent scores under RfamGen, while rCM sequences (used here as a control), have lower scores. This suggests that RfamGen does not distinguish between RBM generated samples and the natural sequences.

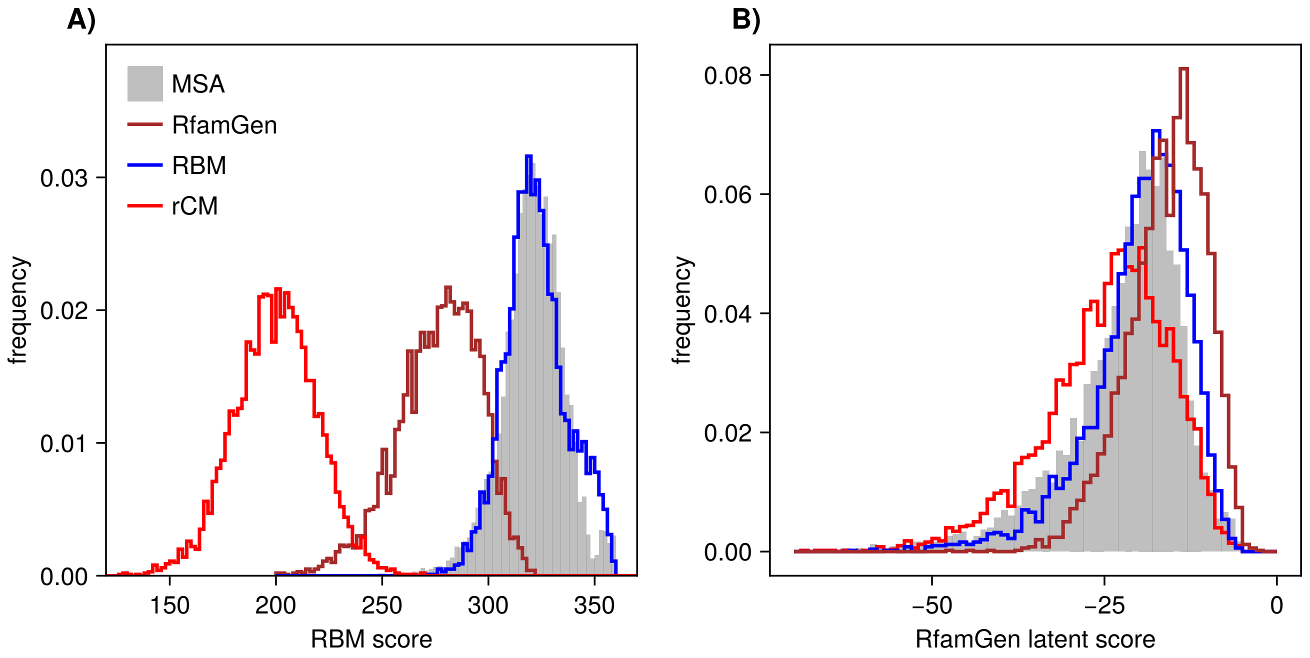


FIG. S7. Comparison to RfamGen [15]. **A)** RBM scores of RfamGen sampled sequences, compared to natural sequences. **B)** RfamGen latent scores of natural, RBM, rCM, and RfamGen sampled sequences, under the RfamGen model.

### Appendix E: R-scape analysis of covariation in sampled sequences

R-scape [16] is a statistical tool to assess how covariations observed in an input multiple sequence alignment support a conserved secondary structure. While correlations between a pair of sites can be due to structural contacts, they can also be explained by phylogenetic coincidences, *i.e.*, groups of sequences that share a common ancestor. To disentangle this effect, for each possible base-pair, R-scape compares the observed covariation with that expected under the null hypothesis that there is no contact but the sequences share a common ancestor. If the covariation cannot be explained by phylogenetics alone, then R-scape considers it to be significant.

We sampled 5000 RBM sequences, 5000 sequences from the Rfam CM, and 5000 sequences from the Denoised CM. For each of these samples, we used R-scape to detect significantly covarying base-pairs [16]. Besides base-pairs in the consensus secondary structure, which are respected by the CMs and the RBMs alike, we focused on the pseudoknot sites that the CMs are unable to model. Confirming our expectations, we find that RBM generated samples exhibit significant covariation across all 8 sites involved in the 4 pseudoknot base-pairs (all sites with E-values $< 10^{-6}$). However, samples from both CM models have no statistically significant covariation across the pseudoknot.

Finally as a consistency check, we also sampled 5000 sequences from the Untangled CM, designed to reproduce the covariation of natural sequences at the pseudoknot. As expected, R-scape finds significant covariation at the pseudoknot sites in these samples (all sites with E-values $< 10^{-6}$).

The covariation scores and E-values for all base-pairs in each sample computed by R-scape are reported in supplementary tables. R-scape was run with default parameters from the web server `http://eddylab.org/R-scape/`.

### Appendix F: Further analysis of the Rfam, Denoised and Unknotted CM variants

In order to be able to capture more distant sequences in the alignments, the Rfam CM models are regularized, resulting in a less constrained model. We therefore considered two CM variants: Denoised and Unknotted CMs (see Methods in the main text for precise definitions). The regularization of Rfam CM manifests in the fact that it does not fit precisely the conservation profile of natural sequences, as shown in Fig. S8 (second row). On the other hand, we have checked that Denoised and Unknotted CMs fit accurately the MSA conservation profiles, Fig. S8 (third and fourth rows).

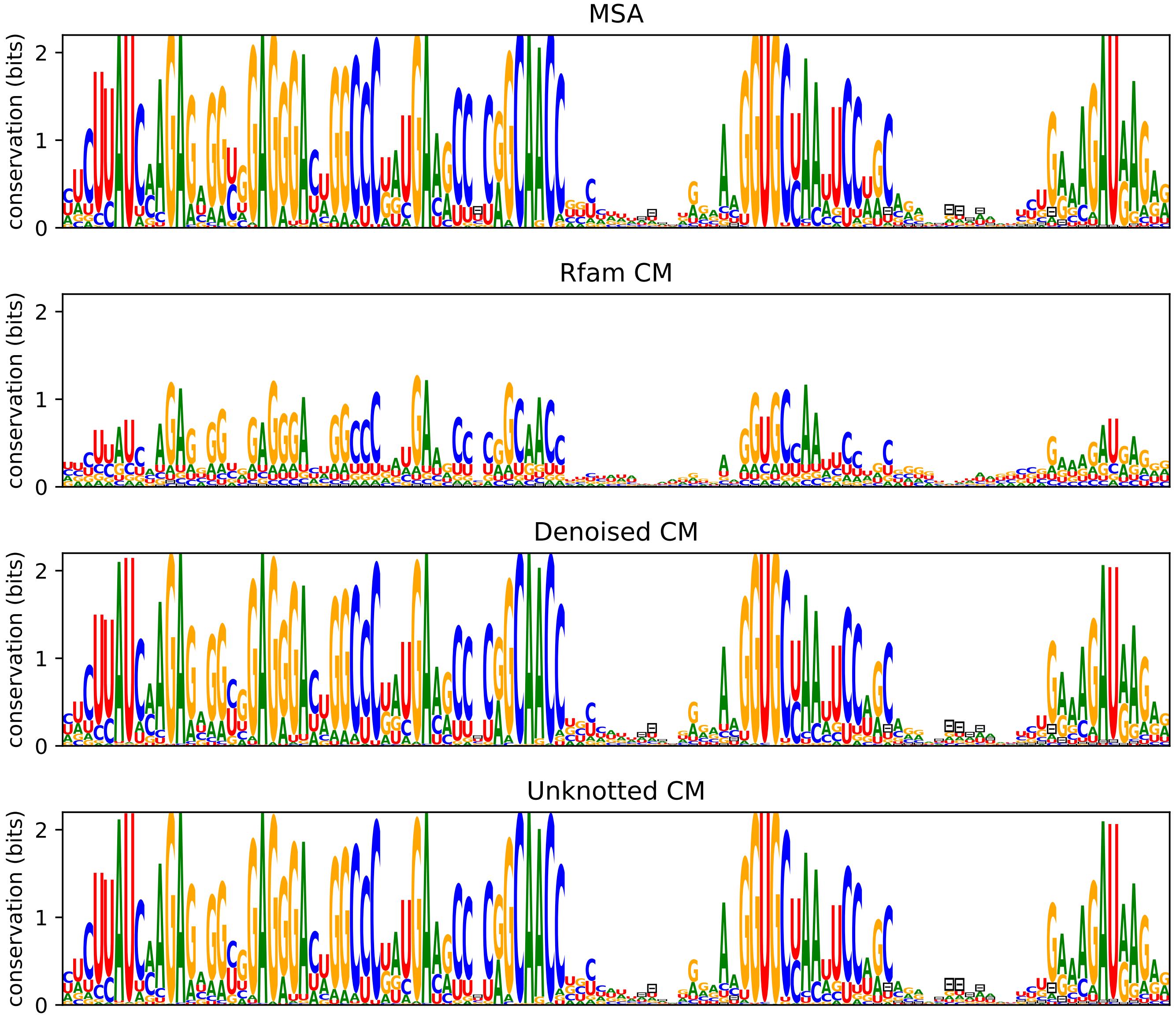


FIG. S8. Sequence logos of sequences sampled from the various CM variants defined in the text. The first row is the sequence logo of the natural MSA. The following rows, correspond to the Rfam CM, the Denoised CM, and the Unknotted CM.

As we saw in Figure 4A in the main-text, the Rfam CM is unlikely to generate sequences of high RBM scores. We corroborate here that also the Denoised and Unknotted CMs are unlikely to sample sequences of high RBM score. We repeated the analyses of Figure 4 from the main-text, replacing the Rfam CM with the Denoised and Unknotted CMs. Results are shown in Supplementary Figure S9).

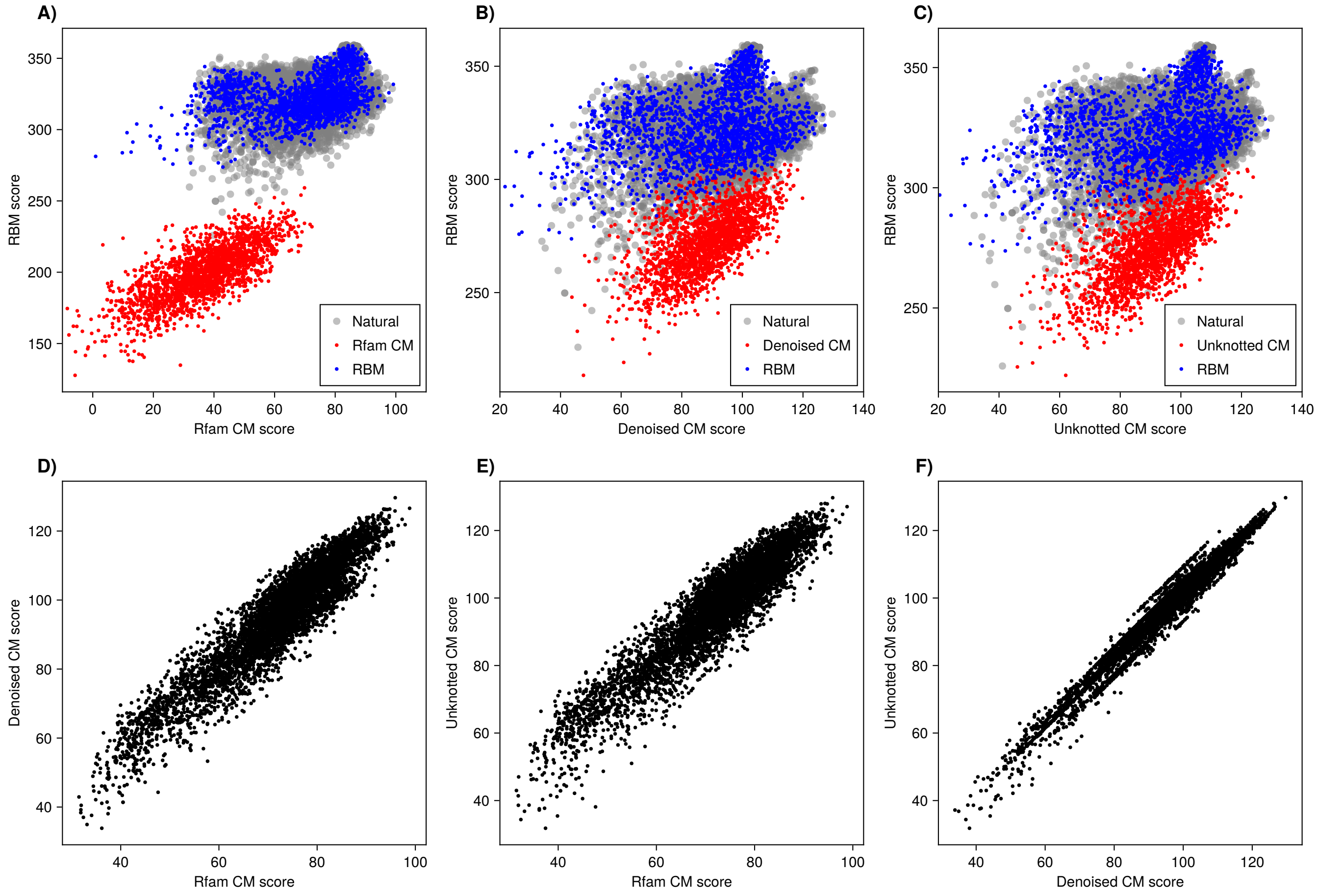


FIG. S9. In the panels in the top row, the $x$-axis gives CM scores and the $y$-axis gives RBM scores, for different classes of sequences. The bottom panels compare CM scores under three CM models of natural sequences. **(A)** Rfam CM. **(B)** Denoised CM. **(C)** Unknotted CM. **(D)** Rfam CM scores vs. Denoised CM scores, for natural sequences. Pearson correlation 0.93. **(E)** Rfam CM scores vs. Unknotted CM scores, for natural sequences. Pearson correlation 0.93. **(F)** Denoised CM scores vs. Unknotted CM scores, for natural sequences. Pearson correlation 0.98.

### Appendix G: Application of RBM to other riboswitch sequence families

Although in this work we have focused on the RF00162 RNA family of SAM-I riboswitch aptamers, we believe that our methodology is generally applicable to other riboswitch families. To support this claim, in this section we have trained RBMs using multiple-sequence alignments of other families from Rfam:

- RF00379, consisting of cyclic di-AMP riboswitches [17]. Like SAM-I riboswitches, these RNAs function as riboswitches, responding to the presence cyclic di-AMP molecules. They are found upstream and regulate the expression *ydaO* and *yuaA* genes in bacteria.
- RF00504, Glycine riboswitch. Regulates glycine degradation pathways [18].
- RF01051, Cyclic di-GMP-I riboswitches, that control the expression of genes involved in numerous fundamental cellular processes in bacteria [19].

We trained RBMs following the same methods used in the main-text. We then sampled new sequences from the RBM, and from the Rfam CM model corresponding to each family. The plots in the first column of SI Fig. S10 compare the RBM score vs. the Rfam CM score computed for these samples, as well as for the natural sequences. Like we found for SAM-I riboswitches (Figure 4 of the main text), we observe that RBM samples tend to have similar CM scores as natural sequences, suggesting they are compatible with the consensus secondary structure captured by the CM. In contrast, CM samples have lower RBM scores than natural sequences, suggesting that the RBM captures additional features beyond those modeled by the CM. The same phenomenon is observed for all three RNA families considered here.

We next considered a PCA projection of the RBM and CM samples, shown in the second and third columns of Figure S10. As we can observe, RBM samples seem to cover the same region of the plane spanned by the top two PCs as the natural sequences. In contrast, CM samples explore only a limited region of this space. Lastly, we consider the foldability of the sampled sequences, as evaluated by ViennaRNA [13] program (using the consensus secondary structure reported for each family). The histogram of folding energies of RBM samples is compatible with the distribution of folding energies of natural sequences. In contrast, Rfam CM samples have a bias towards more unstable energies.

Overall, these results are consistent with the observations we have made for SAM-I riboswitches in the main text. They suggest that our methods are widely applicable to other riboswitch families, and potentially also other RNA families.

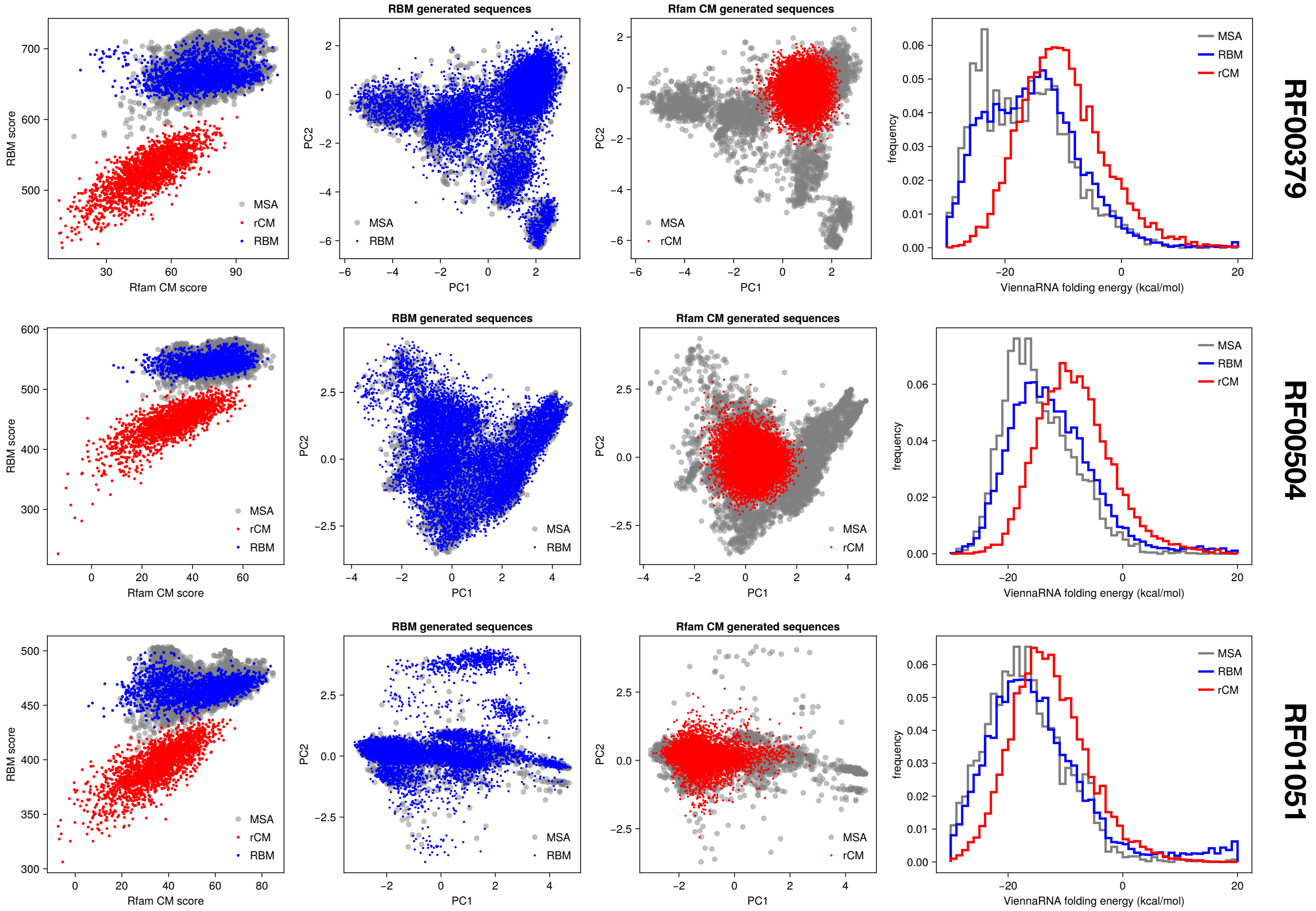


FIG. S10. Application of RBM to other RNA families from Rfam: RF00379 (cyclic di-AMP riboswitch), RF00504 (Glycine riboswitch) and RF01051 (Cyclic di-GMP-I riboswitch).

### Appendix H: Examples of SHAPE-MaP results with manual validation of natural and artificial aptamers

In order to provide a detailed illustration of the SHAPE-MAP results on some of the studied RNAs and on the manual inspection of the data that we have performed in parallel to the automatic analysis, we show in the following 8 examples (4 for natural and 4 for artificial aptamers) obtained from the sequencing data of the 206 natural aptamers and the 100 synthetic first generation designed aptamers, which we had in three consistent replicates (Pearson correlation coefficient > 0.7, Supp. Fig.S14 ). The raw sequencing data have been aggregated before being derived into reactivity maps by Shapemapper 2. Data have been treated this way instead of simply averaging reactivities in order to gain sequencing depth and increase the coverage notably on the 5' and 3' termini for each aptamer. This allows us to better monitor P1 folding and unfolding. We have also included as a benchmark the yitJ aptamer from B. subtilis (PDB id : 4KQY) for which the X-ray structure is available and has been widely studied by various methods6–9. Binding determinants as defined by analyzing 4KQY structure are U8, G12, A47, U79G80, U113 (yitJ aptamer nucleotide numbering). All those nucleotides should undergo a reactivity drop upon SAM binding. However, as U8, U79, G80, U113 are base paired in the apo form of the aptamer and therefore not reactive their reactivity may not change upon binding. Then G12 and A47 reactivity drop are diagnostic of SAM binding. In addition, for most of them an extensive structuration around SAM renders the binding even more obvious. This structural switch is evidenced by a reactivity drop at A25, U86, A110 (base triple), N26-N29 /N87-N90 (pseudoknot), A84-A85 (A minor motif), and N1-N9/N111-N119 (P1 stabilisation), and for some of them the stabilization of nucleotides within the kink-turn motif (positions 19-21/33-38). Aptamers display different degrees of pre-formation of the binding site and consequently the extent of the structural consequences may vary. For some of them we observe a real structural switch while for some other it should rather be qualified as a tertiary structure stabilization. Examples of reactivity profiles observed for natural aptamers (3 replicates), aptamers from first RBM generation (3 replicates) and second RBM generation (1 replicate) as well as the benchmark yitJ aptamer from B. subtilis (PDB id : 4KQY) that we have now added to the study are shown below. SHAPE reactivity are displayed as histograms and color encoded on the structure predicted by IPANEMAP (white: poor reactivity; yellow moderate reactivity, red : high reactivity). The left panel represents the SHAPE probing in absence of SAM while the right panel is the aptamer probing result in presence of SAM.

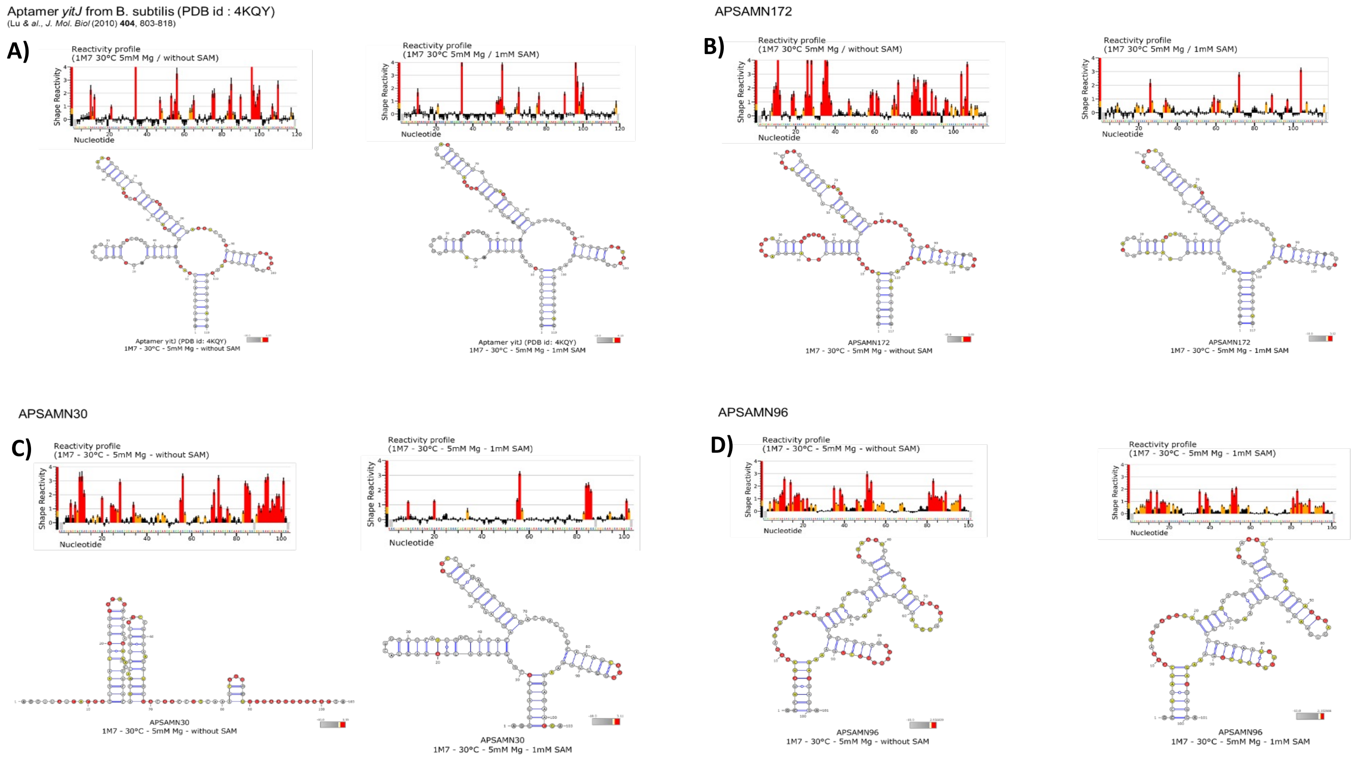


FIG. S11. Examples of SHAPE reactivity profiles on 4 Natural SAM-Riboswitches Aptamers. **A)** Binding is evidenced by G12 and A47 reactivity drop. Other sites are not reactive in the apo form. A structural stabilization is evidenced by A84-A85 and A110 reactivity drop. Other sites are not reactive in the apo form. The pseudoknot and P1 are stable in the apo form. **B)** Binding is evidenced by G11 A47 and U112 reactivity drop. Other sites are not reactive in the apo form. The structural switch is evidenced by the formation of the pseudoknot (U26-G29/C83-G86), the A-minor motif (A79-A80), the base triple (A24-U82-A100), as well as the stabilization of the kink turn as shown by the reactivity drop of the nucleotides involved. An overall stabilization of the aptamer is observed. However, P1 is preformed in the apo state. **C)** Binding is evidenced by U7, G11, A47-A48, U66, U98 reactivity drop. The extensive structural switch is evidenced by the formation of P1 (3-9/ 95-101), the pseudoknot (A25-G28/C74-G77), the A-Minor A72-A73, as shown by the reactivity drop of the nucleotides involved. **D)** This aptamer does not bind SAM (no reactivity profile change upon SAM addition) and is not folded in the proficient structure

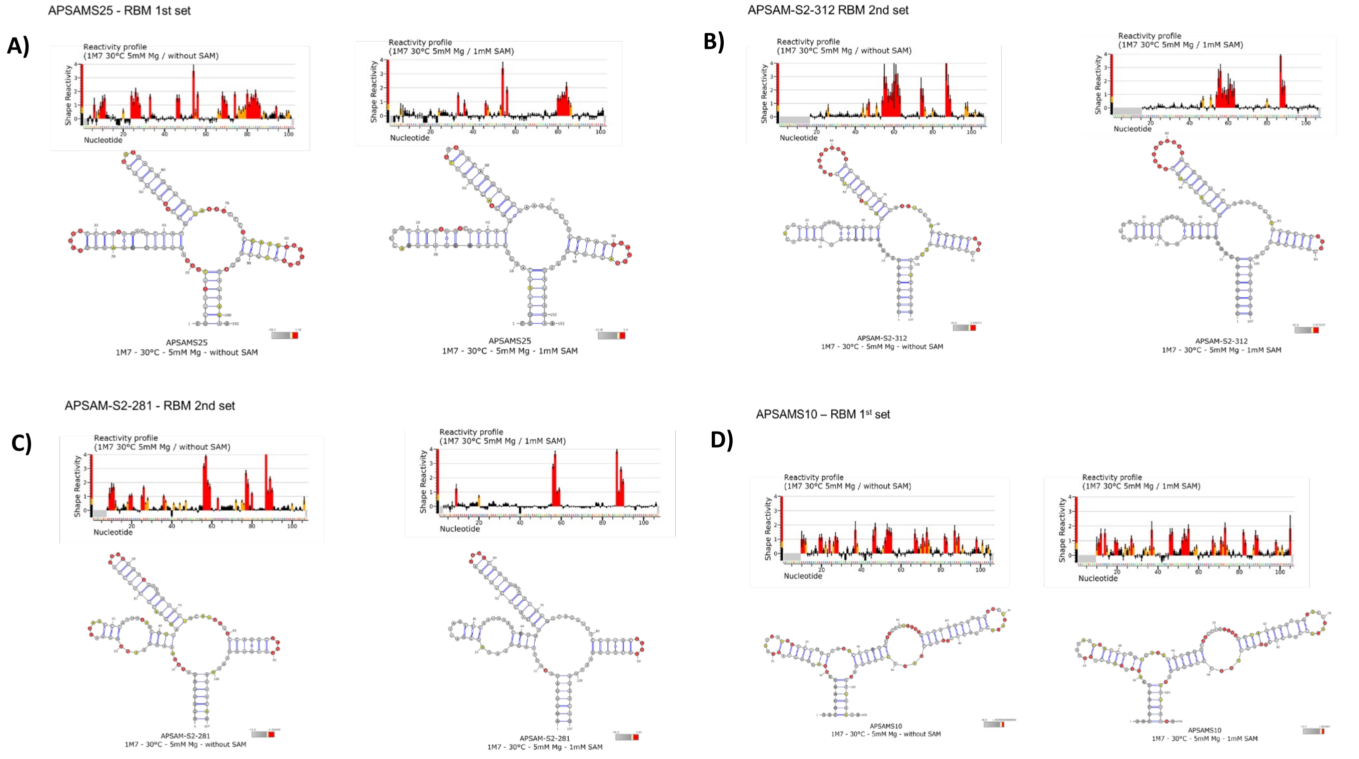


FIG. S12. Examples of SHAPE reactivity profiles on 4 Artificial SAM-Riboswitch Aptamer generated by RBM. **A)** Binding is evidenced by G11 reactivity drop, A46-A47 reactivity decrease but stays high probably reflecting some flexibility in this binding site. Other sites are not reactive in the apo form. The extensive structural switch is evidenced by the formation of the pseudoknot (C25-G28/C71-G74), the base triple (A24-U70-A94), the A minor motif (A68-69) resulting in a reactivity decrease of the nucleotides involved. A slight decrease is also observed for some P1 nucleotides. **B)** Binding is evidenced by G11, A47 reactivity drop other sites are not reactive in the apo form The structural switch is evidenced by the formation of the pseudoknot (U25-A28/U77-A80), and the kink-turn A18-G20/A36-A37). P1 is stable in the apo form. **C)** Binding is evidenced by A46 reactivity drop G11 is undetermined in our study. A structural stabilization is evidenced is evidenced by A74 (A minor) and U75 (base triple) reactivity drop. The aptamer is poorly reactive in its apo form showing that the tertiary structure pre-exists SAM binding for this aptamer. **D)** This aptamer does not bind SAM (no reactivity profile change upon SAM addition) and is not folded in the proficient structure

### Appendix I: Reactivity histograms of paired and unpaired sites for SHAPE data

Figure S13 shows the histograms of reactivities of paired and unpaired sites, under some modifications of the criteria used to produce Fig. 7A in the main-text. Overall, the histograms produced are similar, and thus robust, to these variations. We have also attempted to assess the impact of experimental noise (*e.g.*, from the finite number of sequencing reads) on the observed histograms in Fig. S14. Although we cannot directly access the idealized reactivities at infinite read depth, we can go in the opposite direction and resample the reactivities according to the distributions distributions $P_{ni}(r|\tilde{r}_{ni})$ to evaluate the effect adding *more* noise. If the histograms do not change significantly, this suggests that the noise, though present, might not play a significant role in affecting our conclusions. Figure S14 shows the resampled reactivity histograms of paired and unpaired sites (using the same site classification as in Fig. 7A of the main text). Although there are some small differences, overall the resampled histogram mostly overlaps with the original histograms from the reactivity dataset.

Figure S15 shows the reactivity histograms of the sites belonging to the P1 helix in two conditions: without SAM (left) and with SAM (right). In the same fashion as the pseudoknot (cf. Fig. 7C in the main text), the P1 reactivities shift from a distribution compatible with unpaired residues in absence of SAM, to a distribution closer to the paired sites reactivities, in the presence of SAM. This result supports the notion that P1 is stabilized when SAM is bound. We remark that, in comparison to the pseudoknot (Fig. 7C in the main text), P1 sites in presence of SAM exhibit slightly larger reactivities, indicating that some aptamers might not fully base-pair along this helix in response to SAM. Like Fig. 7C in the main text, this result also supports the validity of the histograms in Fig. 7A as accurate approximations of the statistical reactivities of paired and unpaired sites.

#### 1. On some limitations and robustness of the statistical analysis of reactivity data.

We acknowledge some possible limitations of the pipeline described in the Methods section for the automated analysis of SHAPE-MaP reactivity responses to SAM. First, the definition of the histograms of base-paired and unpaired reactivities (red and blue in Fig. S23A of the main text) rely on the consensus secondary structure of the SAM aptamer. However, it is well known that the riboswitch is flexible and the set of base-paired residues will depend on condition and aptamer.

To further validate the robustness of this pipeline, we have also repeated the analysis, varying the following settings:

- To assess the impact of wrongly annotated sequences, we replaced the histograms 7A, originally computed on all natural sequences, by analogous histograms computed on seed natural sequences only in the alignment. The resulting histograms are very close to 7A in the main text, see also Fig. S13B.
- One can argue that the histograms in Fig. 7A might be convoluted by noise, and further improvements could be obtained by attempting to deconvolve this noise. To assess the impact of the noise, we can go in the opposite direction, of further convolving $P(r|\mathrm{bp})$ and $P(r|\mathrm{np})$ with experimental noise and inspecting the effect on the resulting histograms. More precisely, we can resample the reactivities Eq. (6) from the site distributions $P_{in}(r|\tilde{r}_{in})$ and recompute the histograms using these resampled reactivities in place of the original ones. The resulting histograms suffer minor variations in comparison to 7A, see Supplementary Fig. S14).
- Sites labeled as base-paired / unpaired in the consensus secondary structure can be regarded as being so in only some conditions, and for some aptamers. Two examples are the P1 helix and the pseudoknot, which are generally expected to undergo rearrangements related to SAM binding. In the Supplementary Materials, we have performed the following experiments: i) exclude P1 from both histograms; and ii) include the pseudoknot in the unpaired histogram. See Supplementary Fig. S13C,D. In both cases, the histograms suffer minor variations.

As a consequence, despite these limitations, our conclusions remained unaffected in all the cases we tested.

#### 2. Assessment of false positive rates.

To carry out an assessment of the probability of false positives in our automated analysis, we recomputed protection scores resampling reactivities only from the unpaired sites in the condition with SAM (while the condition without SAM is left unchanged). Focusing on the experiments reported in Fig. 7 of the main text, we then obtain 13 (false) positive aptamers classified as “responsive”. This is to be compared with 111 found in the actual experimental data.

Similarly, we can estimate false negatives by resampling reactivities only from paired sites in the condition with SAM. However in this case, some of the aptamers fail to respond because the switching contacts are detected preformed in absence of SAM, and we cannot claim that these are actual False Negatives. The results are shown in Table S1.

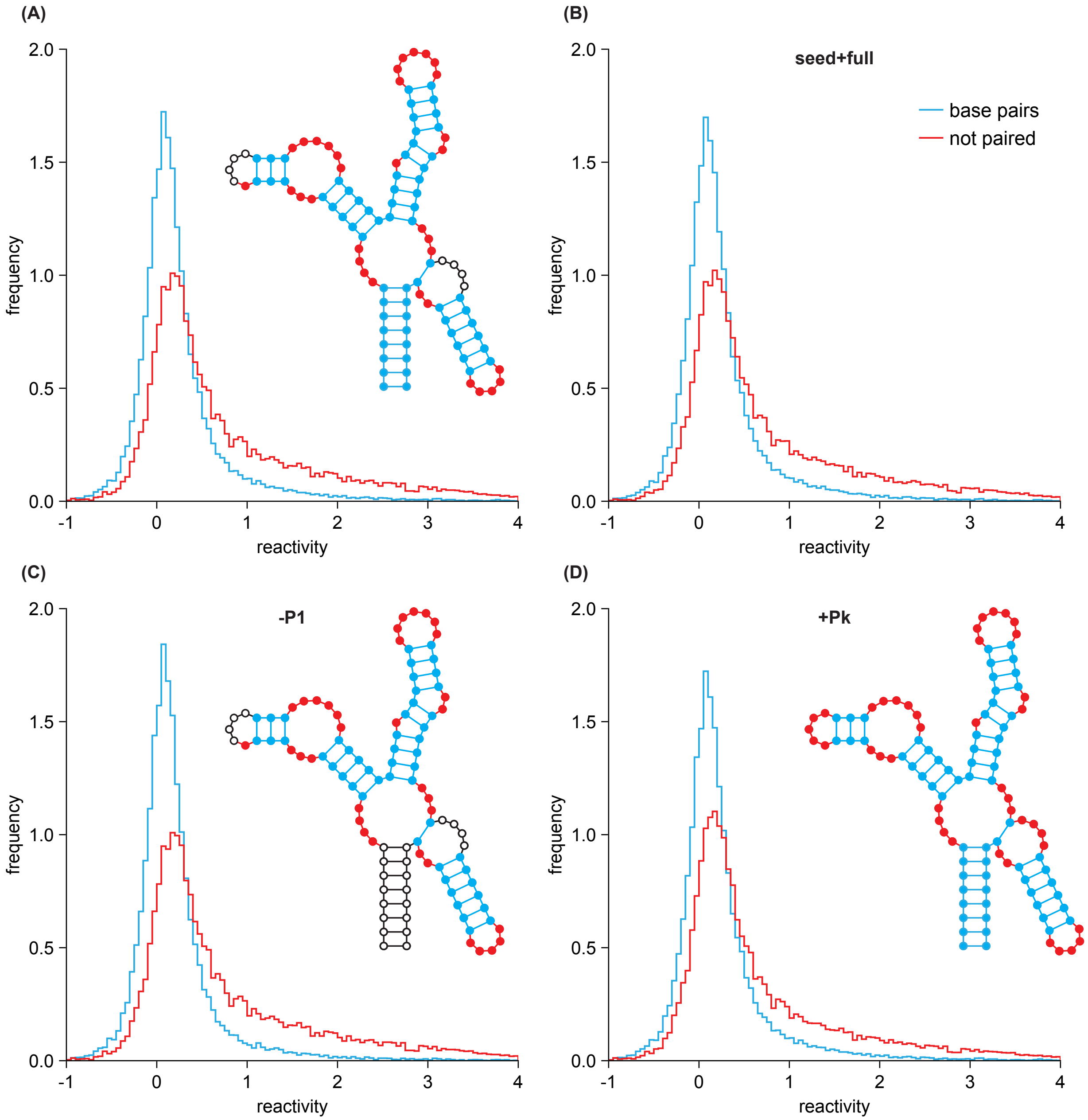


FIG. S13. Histograms of paired and unpaired site reactivities, according to different criteria. **(A)** Same as Figure 7A of the main text. **(B)** Like (A), but including all natural sequences (seed MSA + full MSA). **(C)** Like (A), but excluding the P1 helix from the set of base-paired sites. **(D)** Like (A), but including the sites involved in the pseudoknot in the set of unpaired sites. In the four panels, the inset secondary structure indicates the sites included as base-paired (blue), as unpaired (red), and sites excluded from both histograms (unfilled circles).

Out of the 78 aptamers found to be non-responsive in the always-paired resampled data (last column of the table), 12 fail because the switch contacts are detected as preformed. We exclude them from the analysis. Therefore, the false positive rate (FPR) and the false negative rate (FNR) are estimated as:

$$\mathrm{FPR} = 13/(13+208) = 0.059 \tag{S1}$$

$$\mathrm{FNR} = 66/(134+66) = 0.33 \tag{S2}$$

According to these estimates, our analysis has a low probability (only $\approx 6\%$) of detecting false responses. However there is a larger probability ($\approx 33\%$) that true responses are left undetected. This reflects the nature of the SHAPE

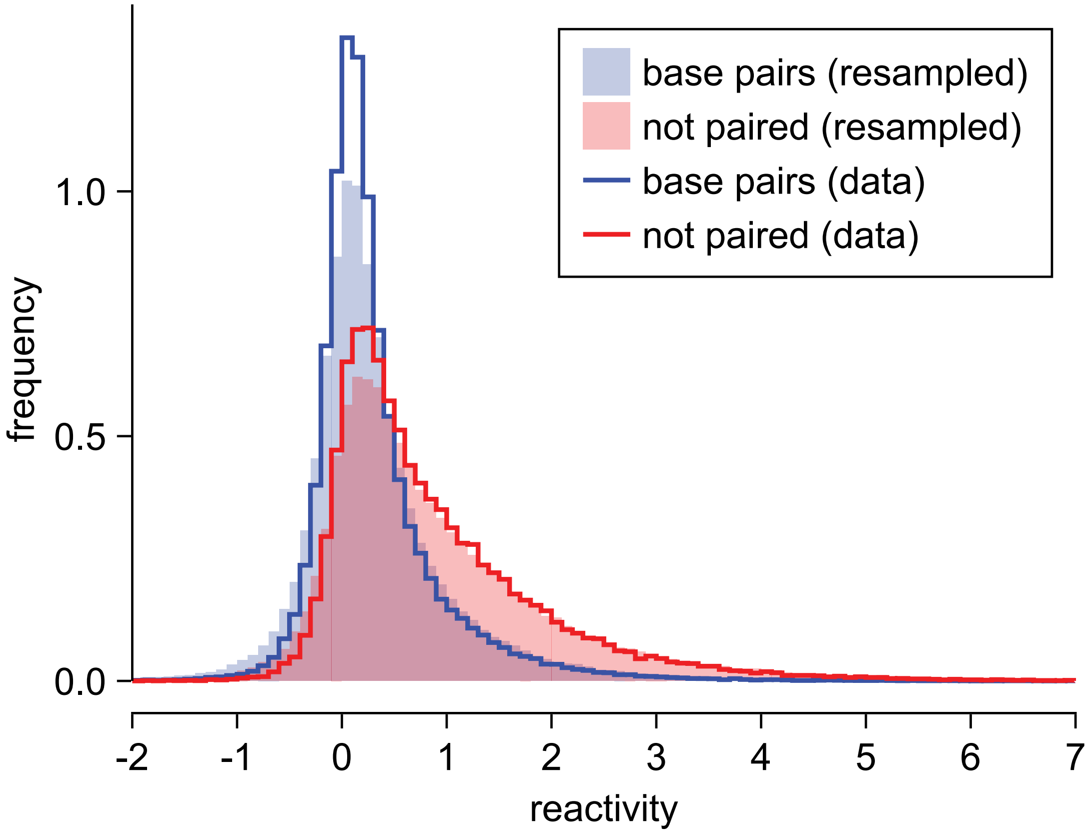


FIG. S14. Histograms of paired and unpaired site reactivities, after resampling reactivities from the site distributions $P_{ni}(r|\tilde{r}_{ni})$ defined in the main-text. The original histograms are shown in continuous line, while the resampled histograms are filled.

| | Data | Resampled (unpaired) | Resampled (paired) |
|---|---|---|---|
| Responsive | 111 | 13 | 134 |
| Non-Responsive | 107 | 208 | 78 |
| Inconclusive | 88 | 85 | 94 |

TABLE S1. False positive rate estimation for protection score analysis.

data, whereby unreactive sites cannot be concluded to be paired. Thus our method tends to be stringent in the kinds of responses it detects (as we also saw in the comparison with a manual analysis).

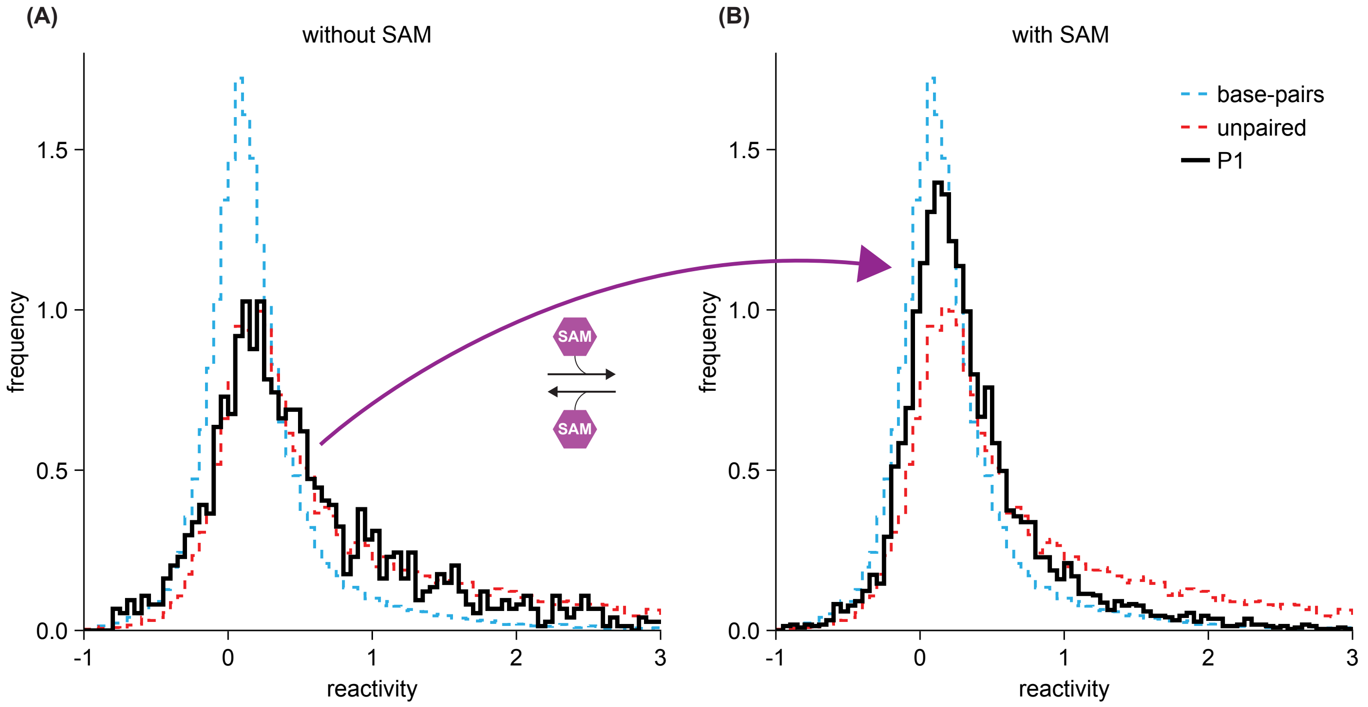


FIG. S15. Stabilization of P1 formation is reflected in SHAPE reactivity histograms. **(A)** Histogram of reactivities of base-paired (blue) and unpaired sites (red) in probed natural sequences belonging to the manually curated seed alignment. Histogram of P1 sites, in absence of SAM, is shown in black, and agrees with the distribution of unpaired sites in absence of SAM. **(B)** In presence of SAM, the reactivities of P1 sites move towards a distribution compatible with base-pairing.

### Appendix J: Supplementary average reactivity profiles

Rfam alignments are constituted of a manually curated alignment of "seed" sequences (which consists of 457 entries for the RF00162 family), and in addition a set of "hits", fetched from genome databases by Infernal as having high bit-score with the Rfam CM. It is plausible that some of the hits could be false positives and thus not true aptamers. In Fig. S16, we show that the average reactivity differences in response to SAM of the full MSA is in excellent agreement to the average reactivities differences of the seed MSA. This suggests that the set of natural sequences in our experiments are able to bind SAM, and respond by performing the expected structural switch.

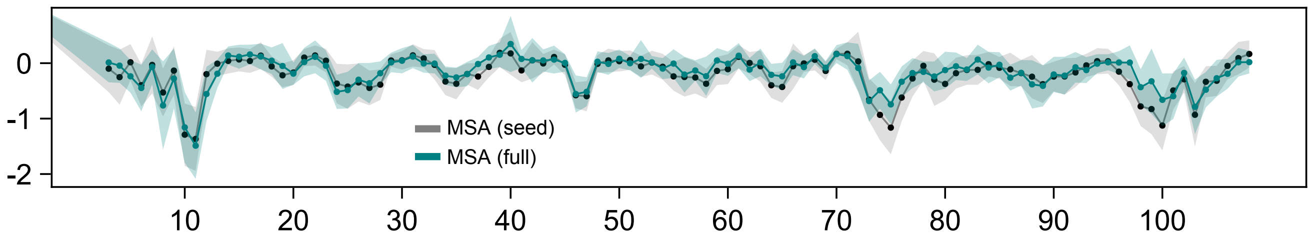


FIG. S16. Average differential reactivities in response to SAM. We compute the reactivity difference of the Mg+SAM condition minus the Mg condition for probed sequences from the full MSA (teal). The plot shows the profile averaged over sequences, with bands showing the ± half one standard deviation. In gray, we show the same plot but for seed sequences.

We also performed experiments in absence and presence of magnesium (Mg), which is known to stabilize the secondary structure of RNA. In Figure S17, we show the average reactivity response to magnesium (Mg) of seed alignment natural sequences, RBM of high and low scores ($> 300$ and $< 300$, respectively), and of the Rfam CM generated sequences. The reactivity responses to Mg of RBM designed sequences of high score are in excellent agreement with those of the natural sequences (Pearson correlation $= 0.89$). In contrast, RBM generated sequences of low RBM scores, or Rfam generated sequences, exhibit larger deviations (Pearson correlations of 0.68 in both cases).

As mentioned in the text, RBM generated sequences of low score also exhibit larger deviations in their reactivity responses to SAM in comparison to the natural sequences. These comparisons are shown in Fig. S18.

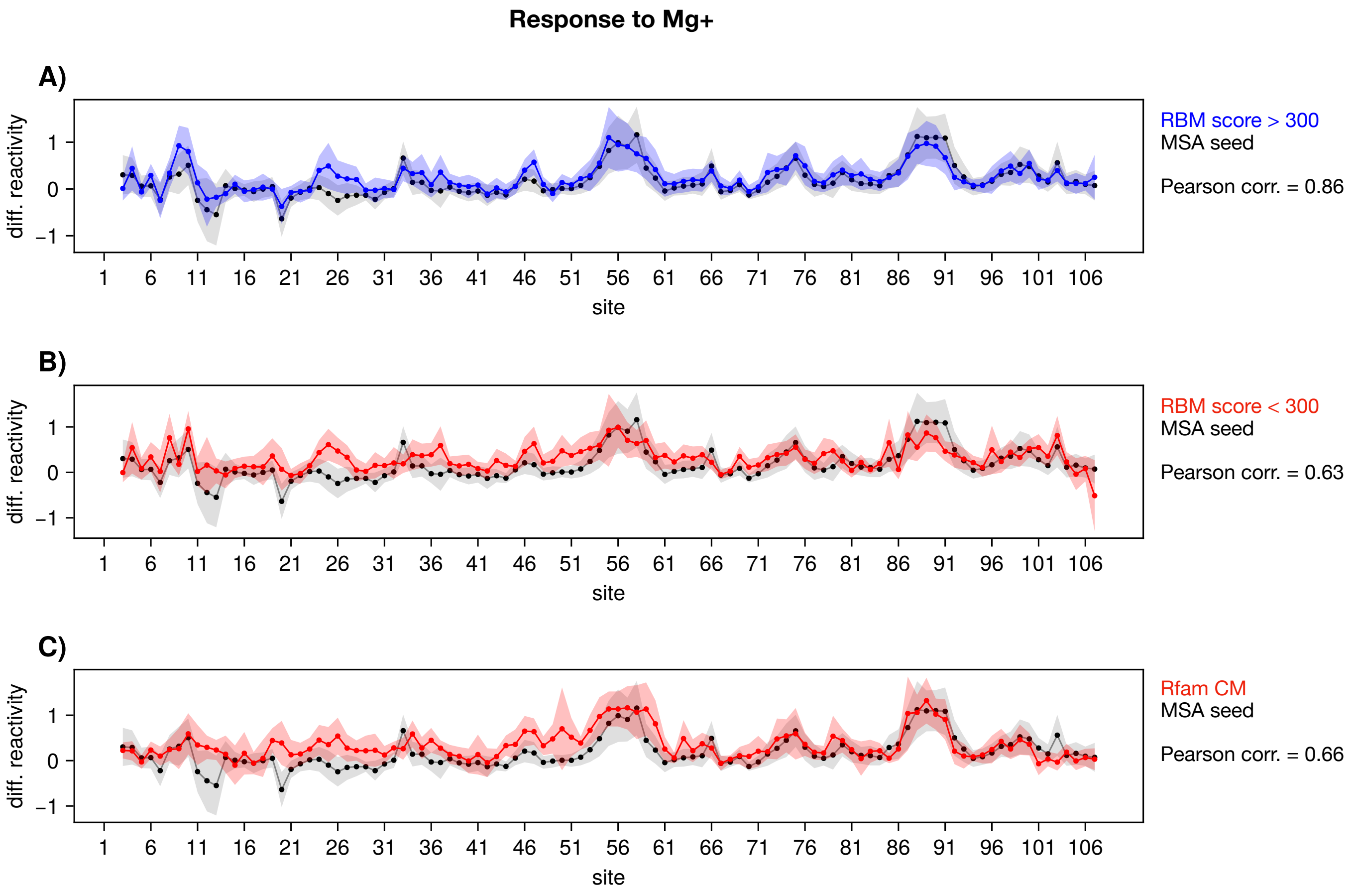


FIG. S17. Average differential reactivities in response to magnesium (Mg). We compute the reactivity difference of the Mg condition minus the 30C (no ligand) condition for different groups of probed sequences. **(A)** Shows the natural seed sequences (in gray) and the RBM sequences with high RBM scores $> 300$. The plot shows the profile averaged over sequences, with bands showing the $\pm$ half one standard deviation. In gray, we show the same plot but for seed sequences. Then panels **(B)** and **(C)** are are analogous but for RBM sequences with low RBM scores $< 300$, and for Rfam CM sequences, respectively. The Pearson correlations to the seed natural sequence reactivity profiles are indicated.

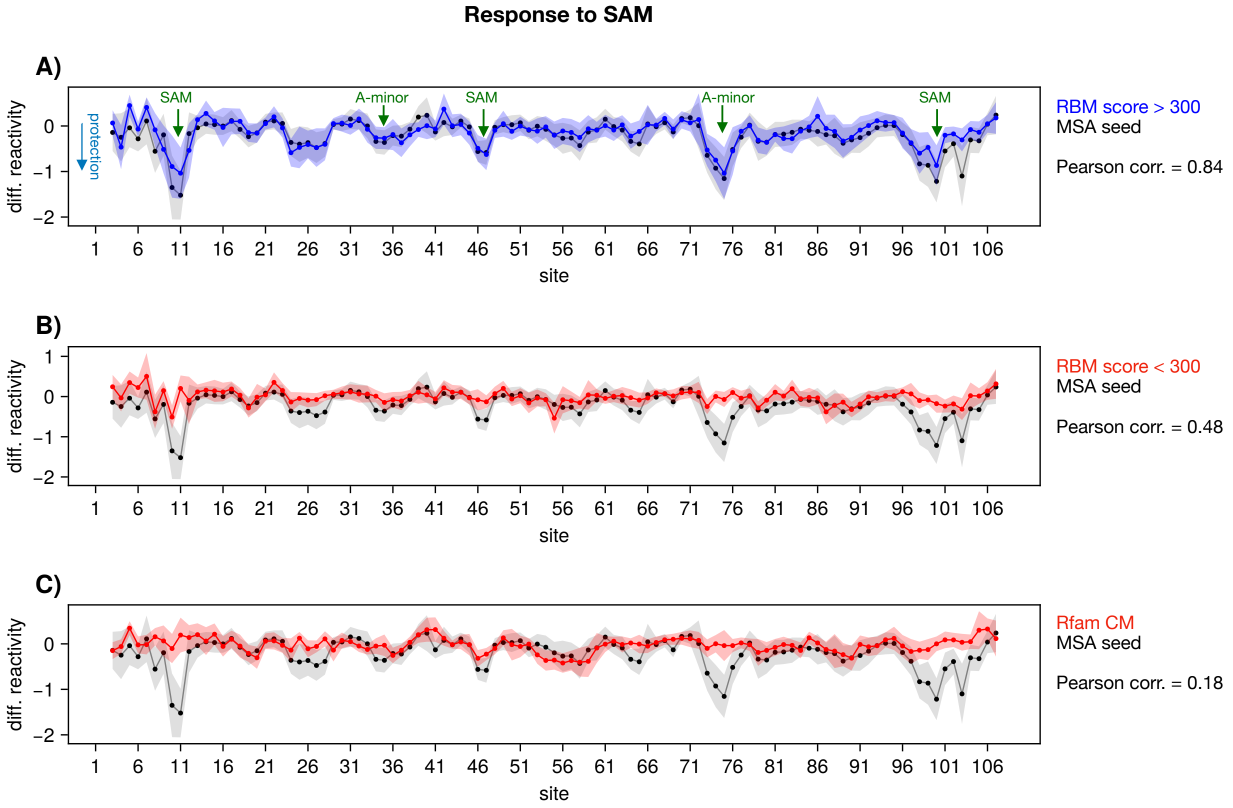


FIG. S18. Average differential reactivities in response to SAM: **(A)** for natural sequences (in gray) and high RBM score (> 300) RBM generated sequences (in blue), **(B)** for natural sequences (in gray) and low RBM score (< 300) RBM generated sequences (in red), and **(C)** for Infernal generated sequences, using the Rfam CM. For each group of sequences and each site, we computed the average reactivity difference of the condition with SAM+Mg minus the condition with Mg only. The plots show the average profiles of sequences in the group, with the bands indicating $\pm$ half one standard deviation. The green arrows indicate locations expected to exhibit reactivity differences in response to SAM binding, while the red arrows highlight lack of protection in some groups of sequences in those locations.

### Appendix K: Replicate experiments

An independent experimental replicate was performed where the same set of 306 sequences as in Fig. 7 of the main text, was probed again by SHAPE-MaP (following the same protocol as in the main text, Section H). We then applied the same analysis pipeline described in the main text (see Section I of the main text) to the replicate experimental data. The overall response to SAM of functional sequences was weaker in both natural and artificial aptamers. Although 112 of the probed aptamers were responsive in the first experiment (hereby called Replicate 0, see Fig. 7), the total number of responsive sequences in the replicate (hereby called Replicate 1) was 75 (see Fig. S19). Among the 75 functional aptamers in Replicate 1, 60 (80%) were also responsive in the first experiment, suggesting that our analysis pipeline is capable of delivering statistically consistent conclusions, across independent experimental realizations.

Fig. 7 in the main text and Supplementary Fig. S19 summarizes these results. Figure S20 compares the reactivities in the replicates to each other in the different conditions probed.

**A)**

| **Group** | **Conclusive** | **Switchers** | **Non-switchers** |
|---|---|---|---|
| Natural | 131 of 201 | 70 (53.4 ± 4.4%) | 61 (46.6 ± 4.4%) |
| Nat.(Seed) | 97 of 151 | 56 (57.7 ± 5.0%) | 41 (42.3 ± 5.0%) |
| Nat.(Hits) | 34 of 50 | 14 (41.2 ± 8.4%) | 20 (58.8 ± 8.4%) |
| Nat.(RBMscore>300) | 87 of 137 | 47 (54.0 ± 5.3%) | 40 (46.0 ± 5.3%) |
| Nat.(RBMscore>310) | 62 of 96 | 38 (61.3 ± 6.2%) | 24 (38.7 ± 6.2%) |
| Rfam CM | 15 of 16 | 0 (0%) | 15 (100%) |
| RBM | 50 of 84 | 7 (14.0 ± 4.9%) | 43 (86.0 ± 4.9%) |
| RBM(RBMscore>300) | 33 of 53 | 7 (21.2 ± 7.1%) | 26 (78.8 ± 7.1%) |
| RBM(RBMscore>310) | 26 of 40 | 7 (26.9 ± 8.7%) | 19 (74.1 ± 8.7%) |
| All | 196 of 301 | 77 (39.3 ± 3.5%) | 119 (60.7 ± 3.5%) |
| All(RBM score>300) | 120 of 190 | 54 (45.0 ± 4.5%) | 66 (55.0 ± 4.5%) |
| All(RBM score>310) | 88 of 136 | 45 (51.1 ± 5.3%) | 43 (48.9 ± 5.3%) |

**B)**

| | Responsive | Non-responsive | Inconclusive | **Total** |
|---|---|---|---|---|
| Switcher | **68** | 2 | 0 | 70 |
| Non-switcher | 38 | **23** | 0 | 61 |
| Inconclusive | 39 | 4 | **27** | 70 |
| **Total** | 145 | 29 | 27 | **201** |

FIG. S19. Panels A) and B) are like Fig. 7A,B of the main text, but for the Replicate experiment.

A further experimental replicate was carried out, restricted to the subset of 206 natural sequences in our dataset. Again, we follow the same protocol as in the main text, Section H. Results are shown in Fig. S22. Although an overall lower response to SAM is observed, the results are overall compatible with Fig. 7 in the main text. Out of the 85 responsive natural aptamers identified in the replicate, 65 (77%) are also responsive in the original experiment. In addition, the Pearson correlations between SHAPE reactivities are $> 0.7$ for all conditions.

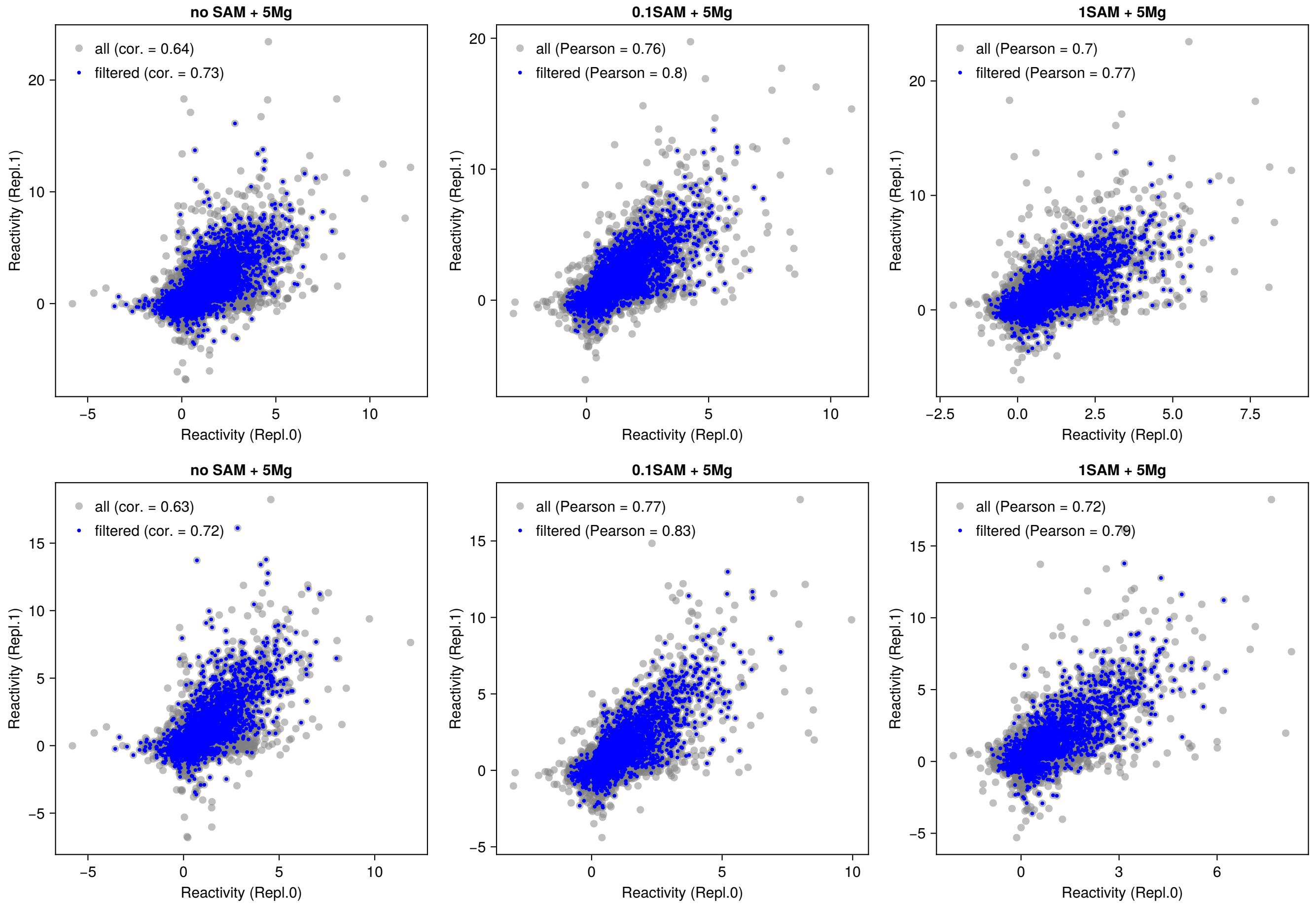


FIG. S20. Comparison of reactivities in both replicates (for the first experimental run). Each panel corresponds to one experimental condition. Pearson correlations are indicated in the legend, for all reactivities (gray), and for filtered reactivities after removing the most noisy values. The retained blue points satisfy: $\mathrm{stderr}(R) < 0.5\sqrt{|R|}$ for both replicates, where $R$ is the SHAPE reactivity and $\mathrm{stderr}(R)$ the standard error, as reported by ShapeMapper [20]. The top row plots all reactivities, while the bottom row is restricted to natural aptamers only (excluding all artificial ones).

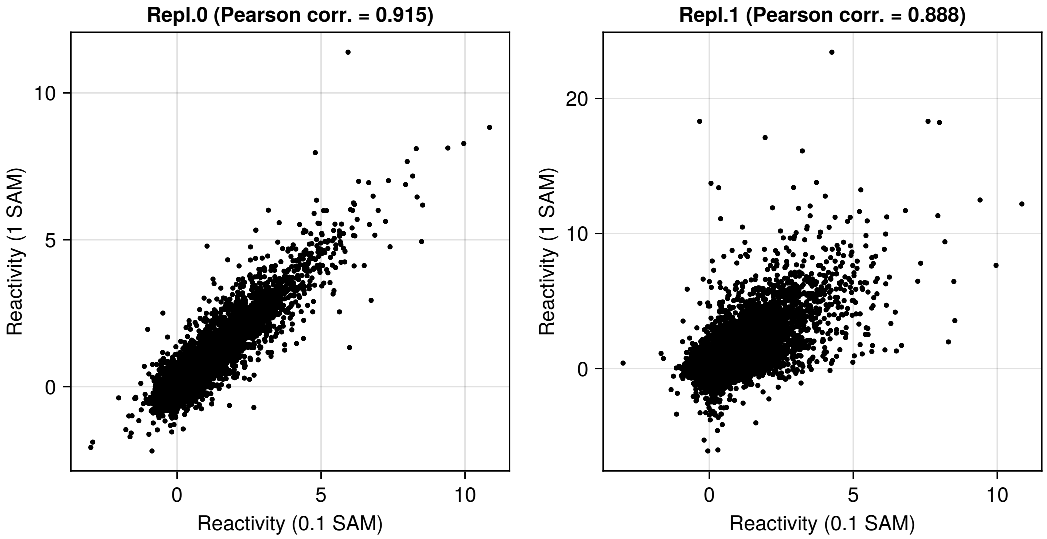


FIG. S21. Comparison of reactivities in both replicates at two concentrations of SAM.

A)

| | Conclusive | Responsive | Non-responsive |
|---|---|---|---|
| Natural | 156 of 201 | 85 (54.5 ± 4.0%) | 71 (45.5 ± 4.0%) |
| Nat.(Seed) | 121 of 151 | 65 (53.7 ± 4.5%) | 56 (46.3 ± 4.5%) |
| Nat.(Hits) | 35 of 50 | 20 (57.1 ± 8.4%) | 15 (42.9 ± 8.4%) |
| Nat.(RBMscore>300) | 106 of 137 | 60 (56.6 ± 4.8%) | 46 (43.4 ± 4.8%) |
| Nat.(RBMscore>310) | 72 of 96 | 47 (65.3 ± 5.6%) | 25 (34.7 ± 5.6%) |

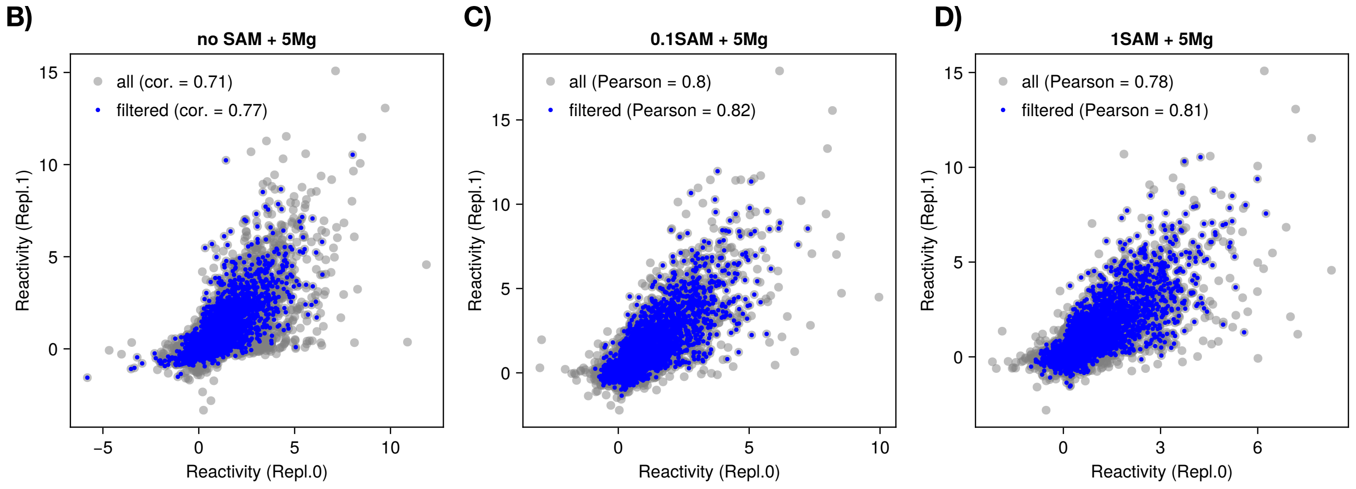


FIG. S22. Experimental probing replicate for natural aptamers. **A)** SAM responses inferred from the automated analysis, for the replicate for the natural aptamers. **B,C,D)** Correlations between SHAPE reactivities (estimated by ShapeMapper) between the replicates for the natural aptamers for the different conditions tested.

### Appendix L: SHAPE analysis

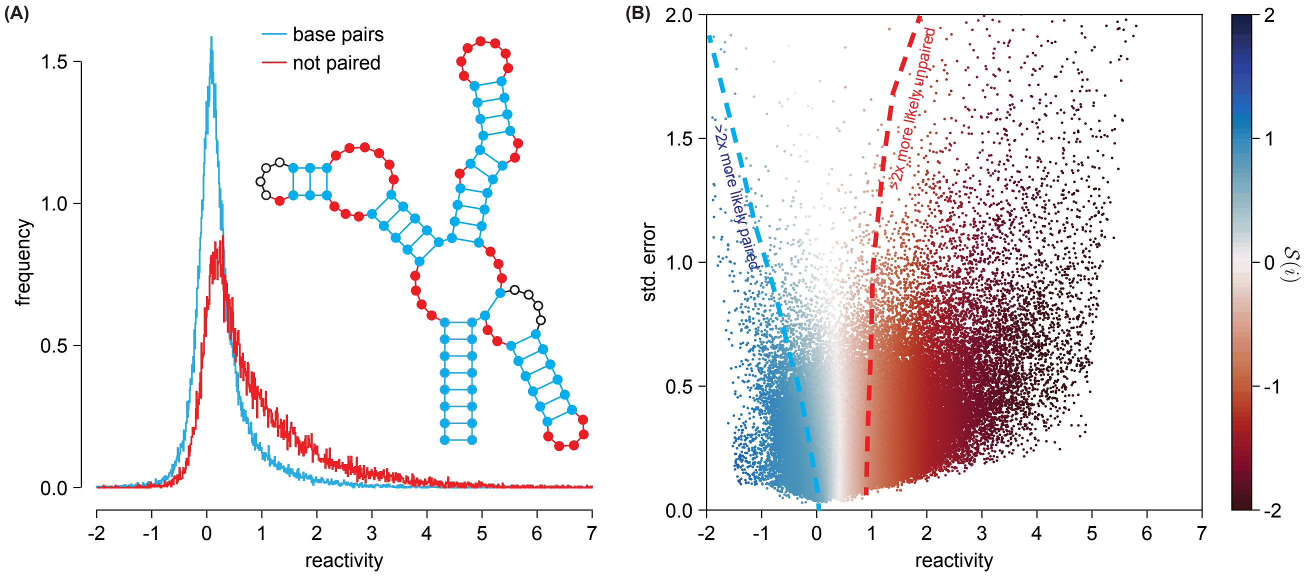


FIG. S23. Statistical differences of SHAPE reactivities in paired and unpaired sites. **(A)** Histogram of reactivities of base-paired (blue) and unpaired sites (red) in probed natural sequences belonging to the manually curated seed alignment. The inset shows the consensus secondary structure with sites colored according to whether they are paired or unpaired. Sites with ambiguous behavior (pseudoknot, indicated by empty circles in the inset secondary structure) are excluded from both histograms. **(B)** Scatter plot of measured reactivities (on the $x$-axis) and their estimated standard errors (on the $y$-axis), as estimated by the standard ShapeMapper protocol [20], by first-order error propagation through the Poisson statistics of the mutation counts. The points are colored by the value of the log-odds ratio $\ln(P(\tilde{r}|\text{bp})/P(\tilde{r}|\text{np}))$ (see color bar), computed as explained in the text surrounding equation (8). The blue (red) dashed line indicates a contour separating sites over two times more likely to be paired (unpaired) than not.

### Appendix M: Read depths of chemical probing experiments

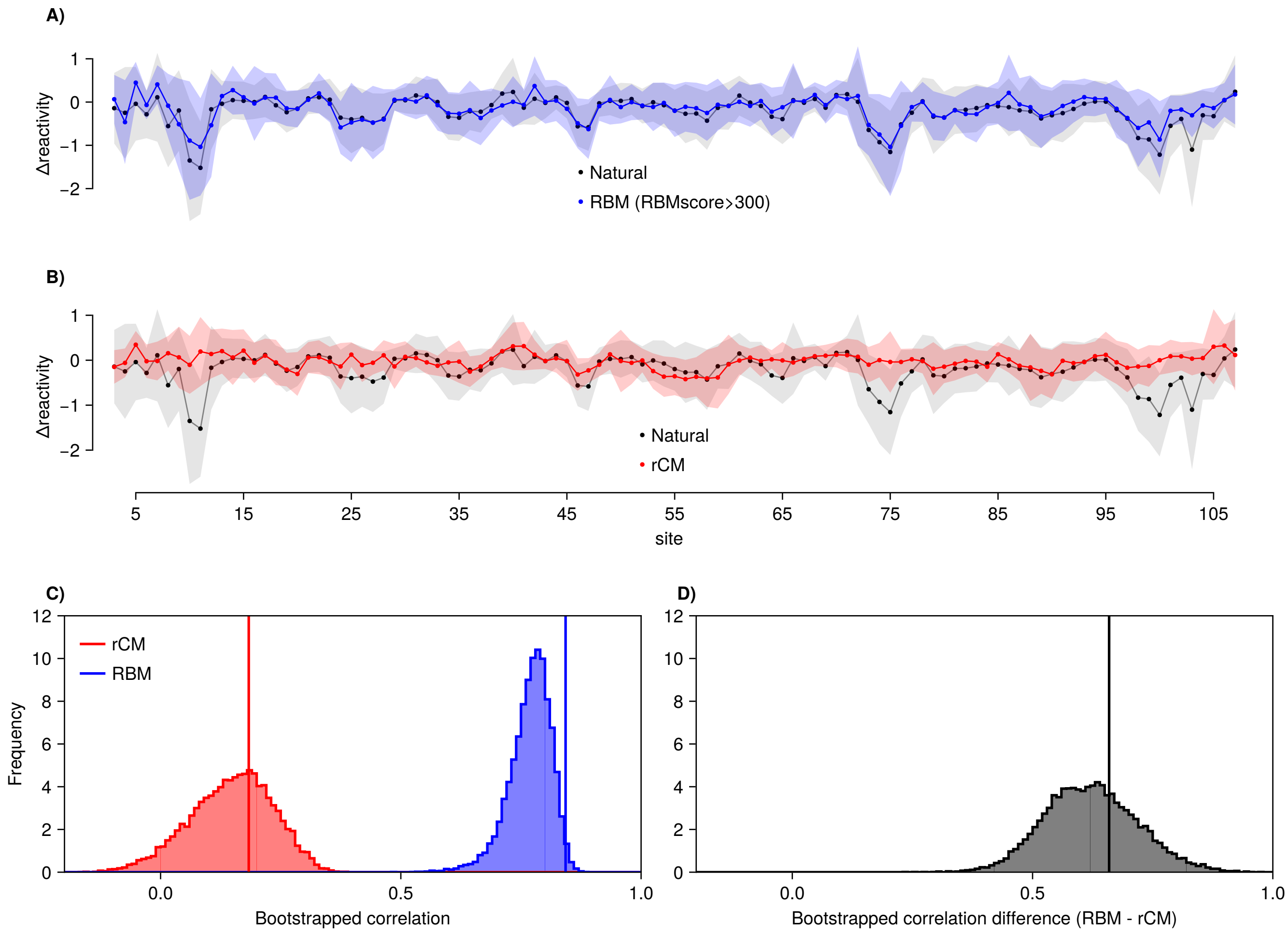


FIG. S24. SHAPE-MaP probing of aptamers. **A)** Average differential reactivities in response to SAM of RBM generated sequences with high RBM scores (> 300) (blue), across the 108 sites of the alignment. For comparison, the average differential reactivities for natural sequences are shown in the background (gray). High-RBM score sequences recapitulate protection of sites involved in the structural switch in response to SAM binding. **B)** Average differential reactivities in response to SAM of Rfam CM (rCM) generated sequences (red). Natural sequences are shown in background for comparison. Rfam CM sequences fail to recapitulate the expected protections associated to the structural switch (red arrows). Panels A,B are like panels 7C,D from the main-text, but with the bands now indicating ± one standard deviation (instead of 1/2 s.t.d. like in the main text). **C)** We resampled aptamer reactivities belonging to different groups of sequences (Natural, RBM, rCM) from the data, and estimated the distribution of Pearson correlations between the average reactivity profiles of the different groups: rCM vs Natural (in red), and RBM vs natural (in blue). The empirical histograms shown correspond to $10^6$ bootstraped samples, while the vertical lines correspond to the correlations computed on the actual observed data. In all realizations the correlation between RBM and Natural reactivity profiles is higher than between rCM and Natural. **D)** For each bootstrapped sample, we show the histogram of the differences between the Pearson correlation of RBM and Natural minus rCM and Natural.

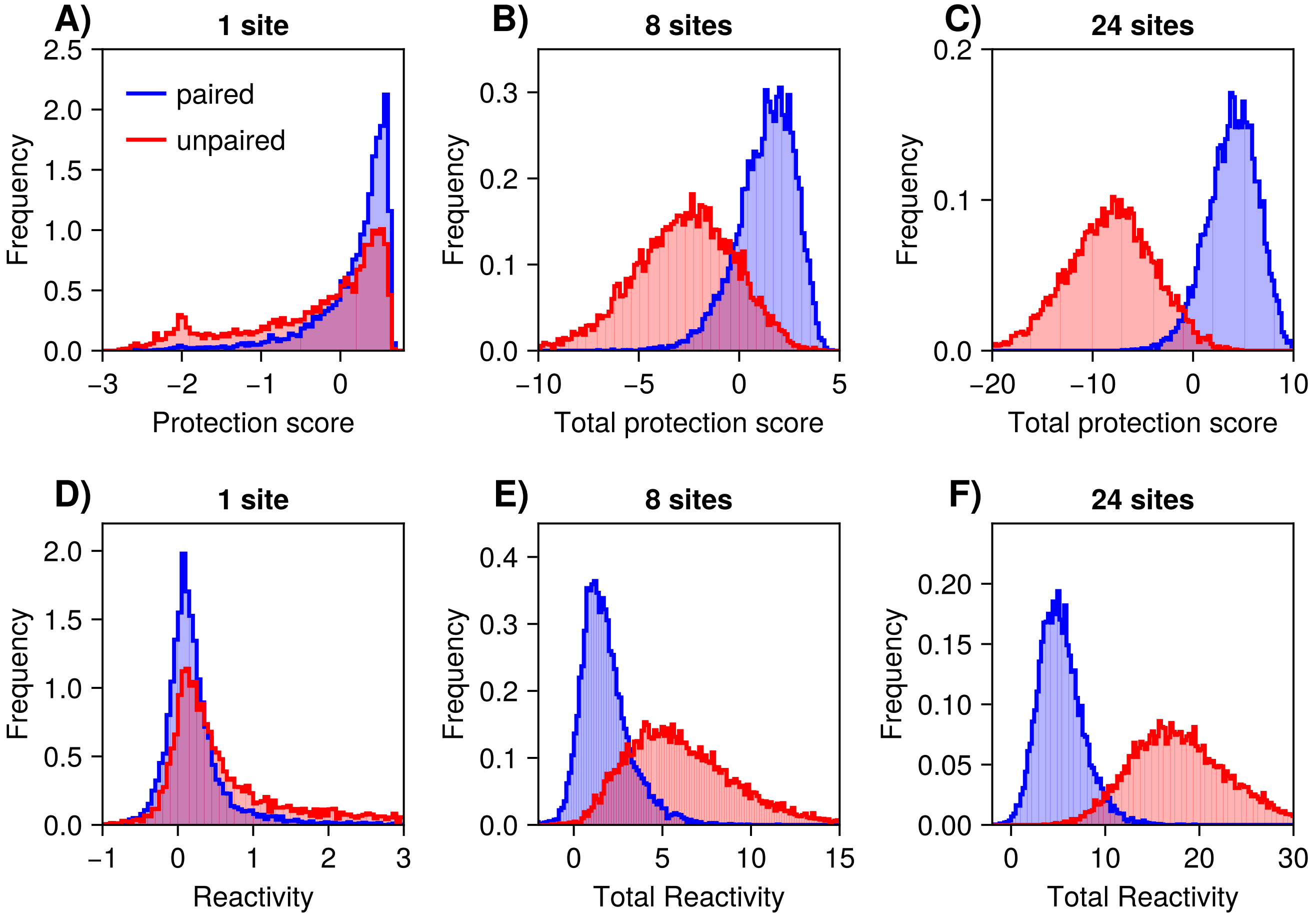


FIG. S25. Using SHAPE reactivity measurements from multiple sites leads to higher discriminative power. **A)** Empirical histogram of the protection scores of paired (blue) and unpaired (red) sites in the natural aptamers probed by SHAPE-MaP. **B)** Total protection score of 8 sites (same number involved in the pseudoknot of SAM-I riboswitches). **C)** Total protection score of 24 sites (same number involved in the Hallmark sites). **D)** Raw SHAPE-MaP reactivities of paired and unpaired sites. **E)** Total reactivities of 8 sites. **F)** Total reactivities of 24 sites. See also Fig. S39 for the counterpart of this figure for DMS data.

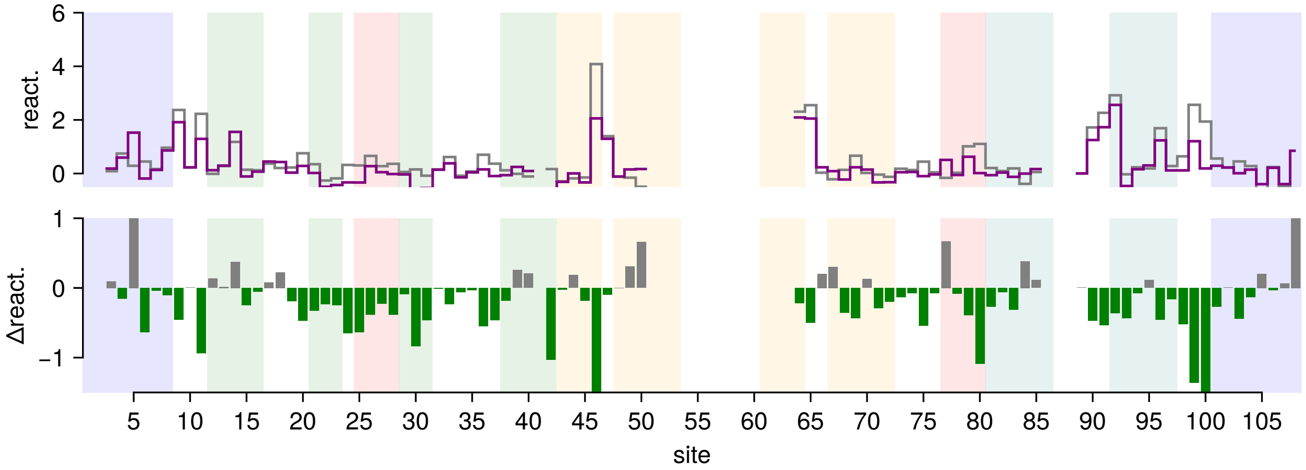


FIG. S26. Reactivity profile of SAM riboswitch aptamer from *T. tengcongensis* (PDB id: 2GIS).

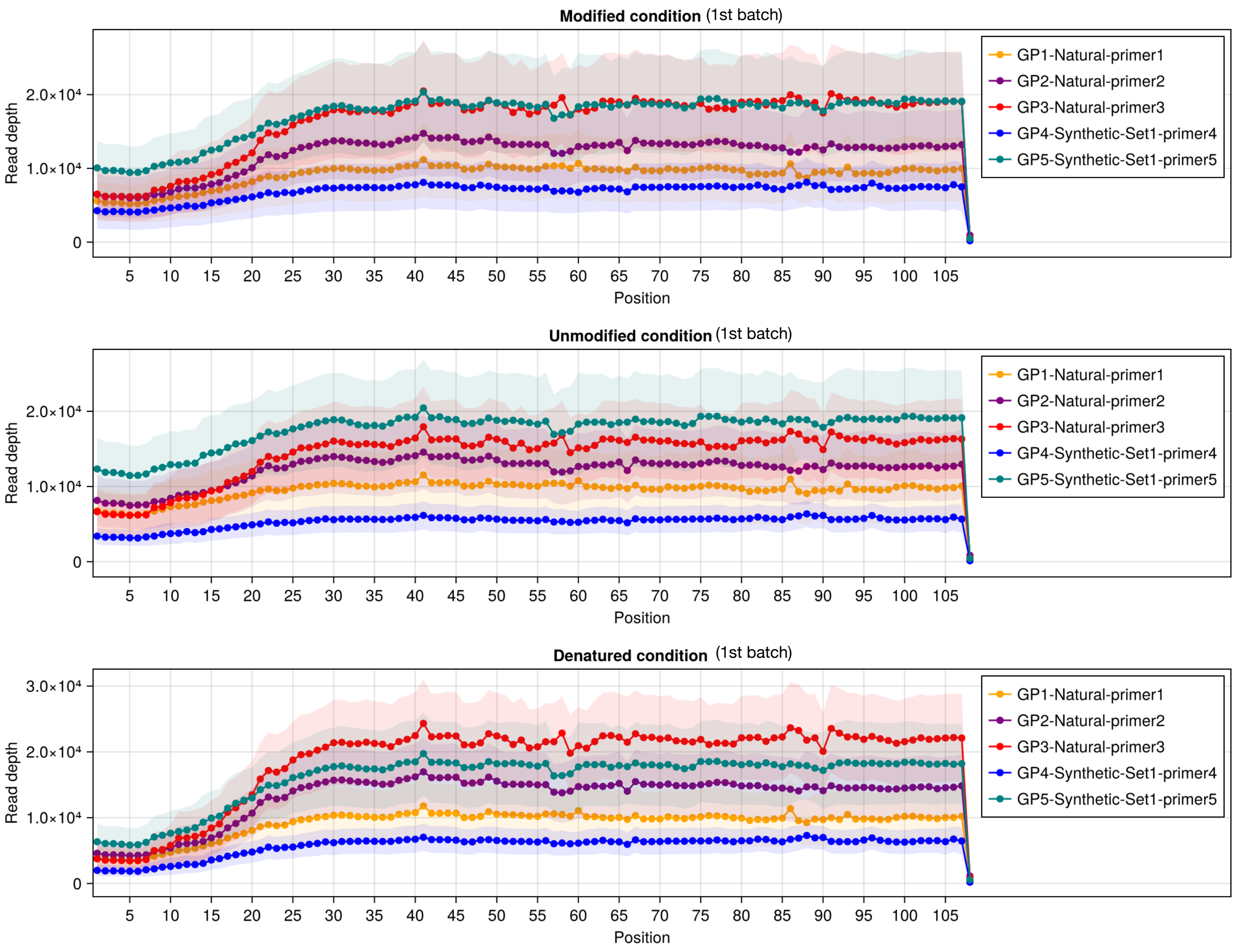


FIG. S27. Read depths for each position in the 1st batch SHAPE-MaP experiments, for each condition: Modified (in presence of the SHAPE reagent), Unmodified (in absence of the SHAPE reagent), and Denatured condition. Aptamers are split into groups according to the primer used (color legend). The solid lines show the average read depths, while the standard deviations are shown as light bands. The read depth is as reported by ShapeMapper [20].

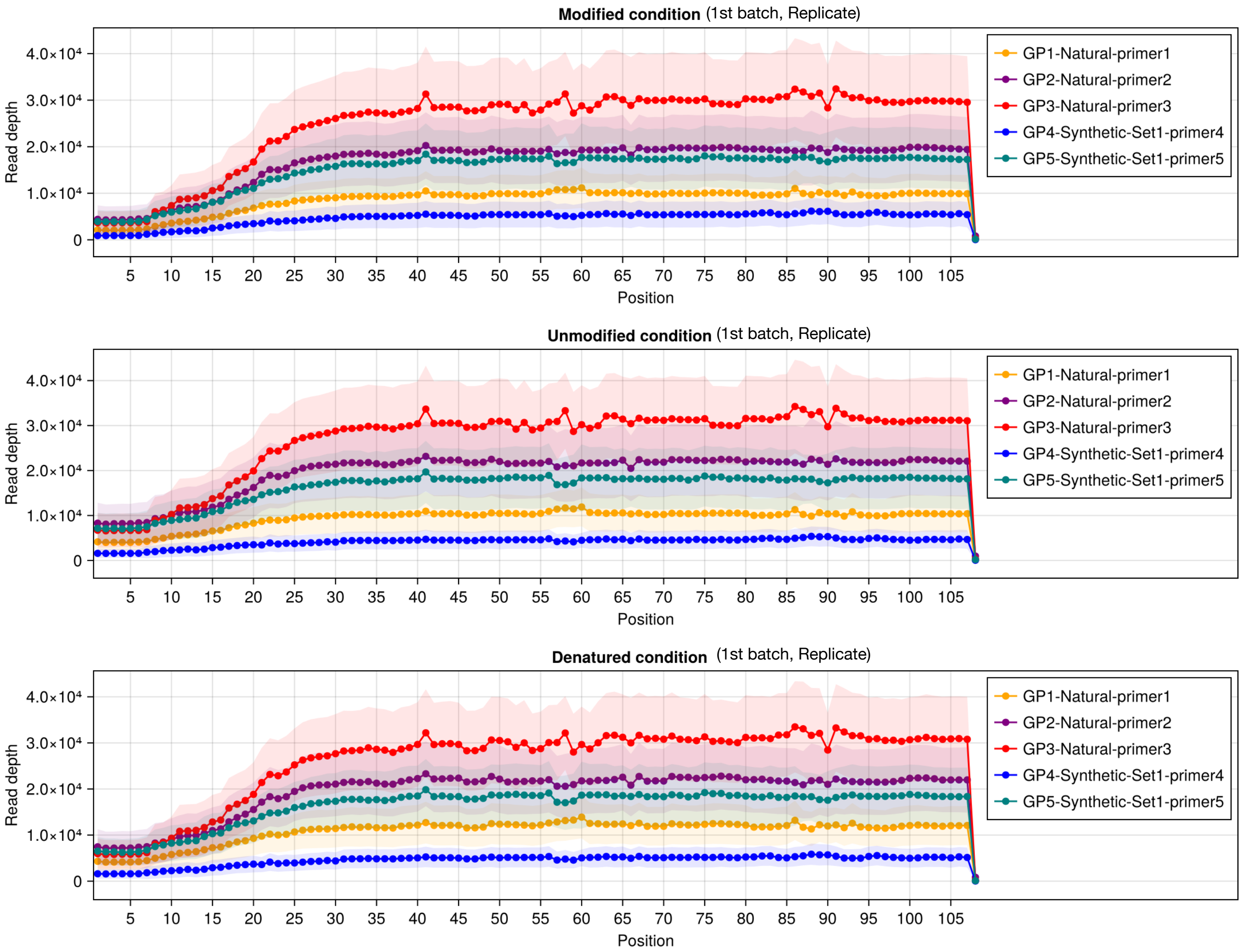


FIG. S28. Read depths for each position in the 1st batch replicate SHAPE-MaP experiments, for each condition: Modified, Unmodified, and Denatured. Aptamers are split into groups according to the primer used (color legend).

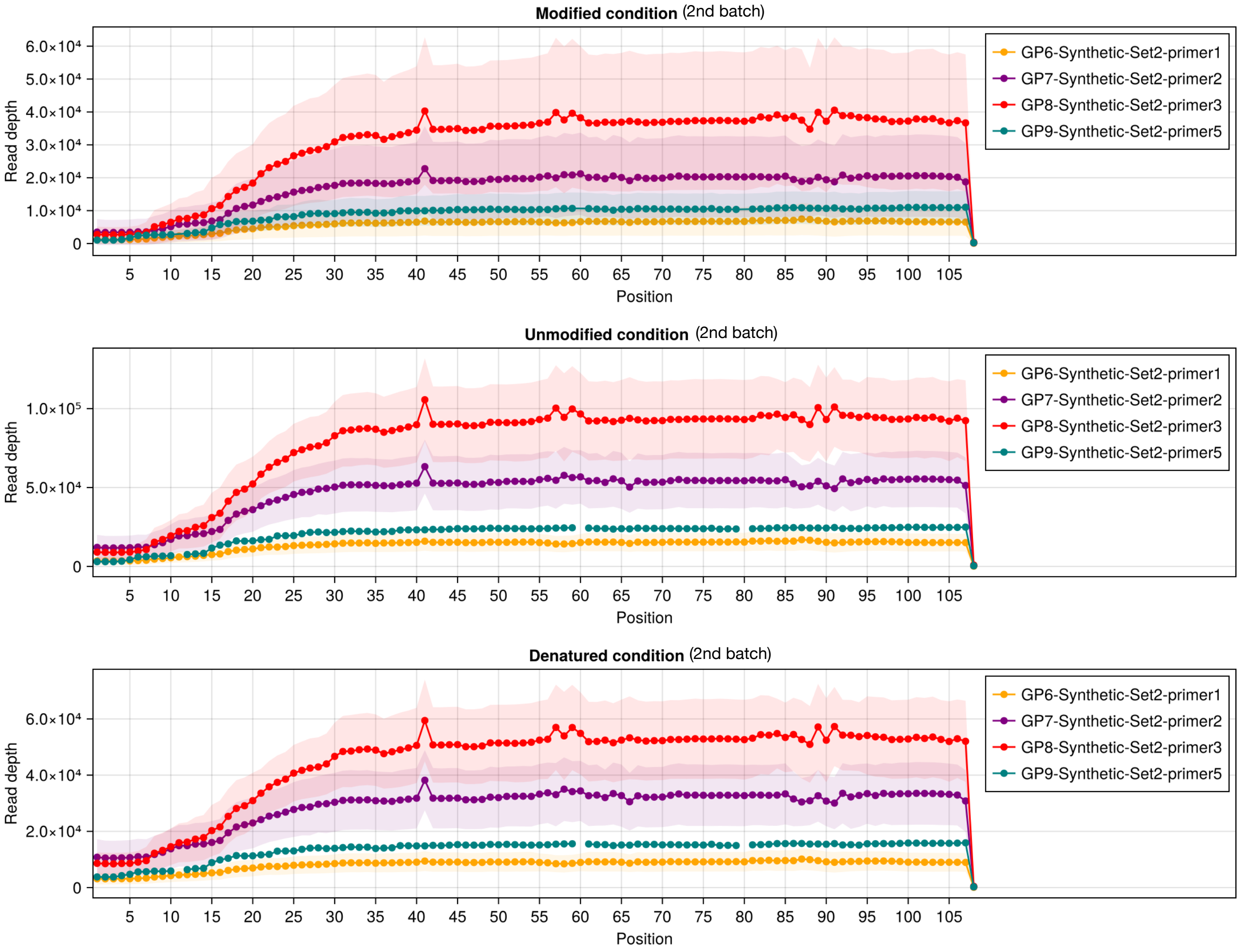


FIG. S29. Read depths for each position in the 2nd batch SHAPE-MaP experiments, for each condition: Modified (M), Unmodified (U), and Denatured (D). Aptamers are split into groups according to the primer used (color legend).

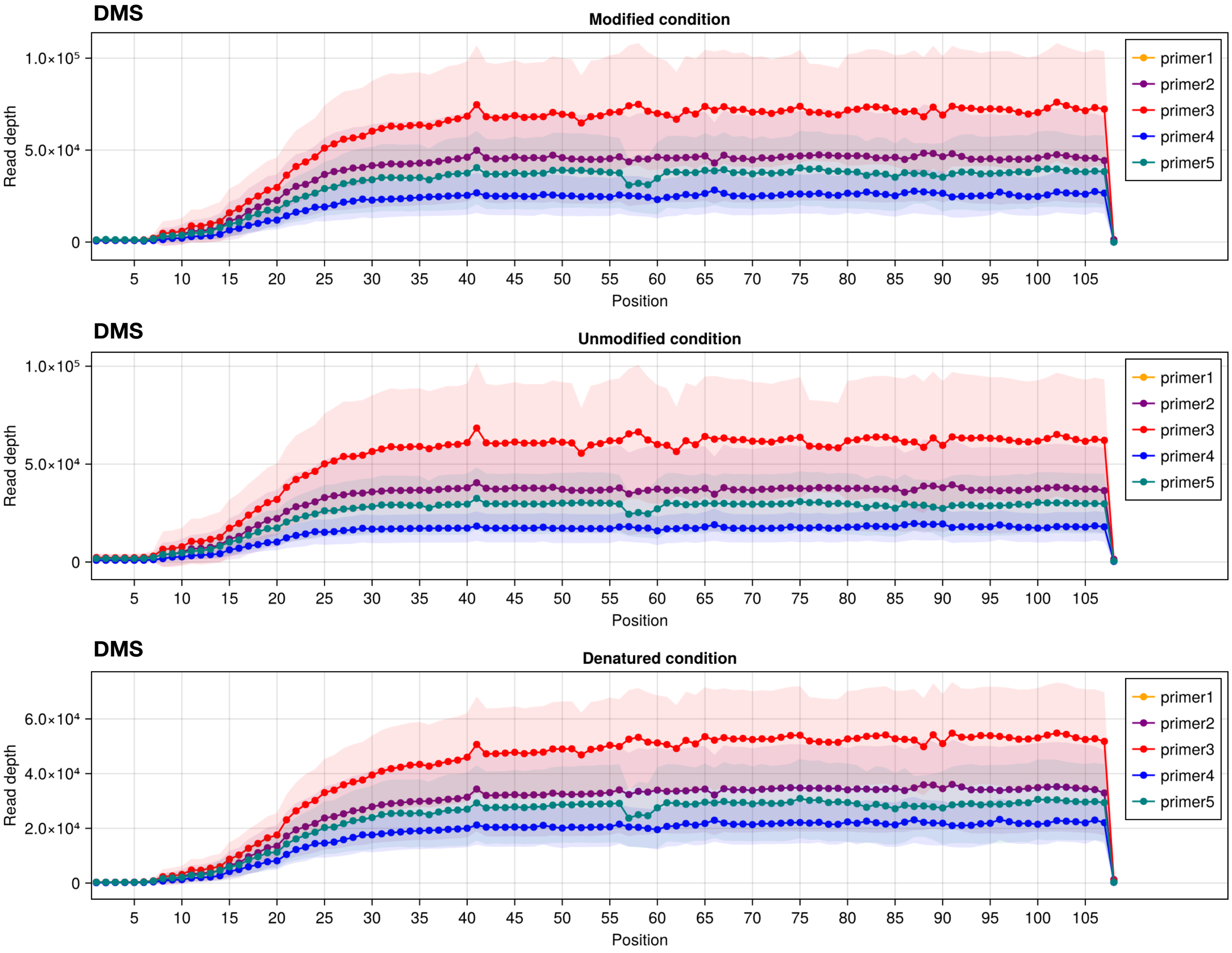


FIG. S30. Read depths for each position in the DMS experiments, for each condition: Modified (M), Unmodified (U), and Denatured (D). Aptamers are split into groups according to the primer used (color legend).

SHAPE ↓ ; DMS →

1st batch

| DMS \ SHAPE | Yes | No | Inc. |
|---|---|---|---|
| Yes | 32 | 9 | 25 |
| No | 1 | 20 | 7 |
| Inc. | 3 | 2 | 22 |

| | Yes | No | Inc. |
|---|---|---|---|
| Yes | 4 | 6 | 4 |
| No | 1 | 4 | 3 |
| Inc. | 0 | 4 | 5 |

| | Yes | No | Inc. |
|---|---|---|---|
| Yes | 0 | 0 | 0 |
| No | 0 | 9 | 2 |
| Inc. | 2 | 9 | 6 |

| | Yes | No | Inc. |
|---|---|---|---|
| Yes | 4 | 4 | 6 |
| No | 0 | 42 | 7 |
| Inc. | 1 | 0 | 8 |

Primer 1: CGTCTGCCCGCCTCCTTCC

Primer 2: CCAGCAGCCGCGGTAATACG

Primer 3: CCAGTGTCCGCCTATCTCGTCG

Primer 4: AGTGTCCGCCTATCTCGTC

Primer 5: ATCGGCTACCTTGTTACGACTTC

2nd batch

| | Yes | No | Inc. |
|---|---|---|---|
| Yes | 3 | 6 | 4 |
| No | 0 | 34 | 3 |
| Inc. | 0 | 10 | 2 |

| | Yes | No | Inc. |
|---|---|---|---|
| Yes | 6 | 3 | 4 |
| No | 0 | 29 | 2 |
| Inc. | 0 | 4 | 2 |

FIG. S31. Comparison of success rates in SHAPE and DMS probing experiments, per primer. The first row corresponds to the first batch of experiments, while the second row corresponds to the second batch. Each column corresponds to one of the primers used in the experiments (the primer sequence is indicated). The numbers of responsive (Yes), non-responsive (No) and inconclusive (Inc.) sequences is then indicated.

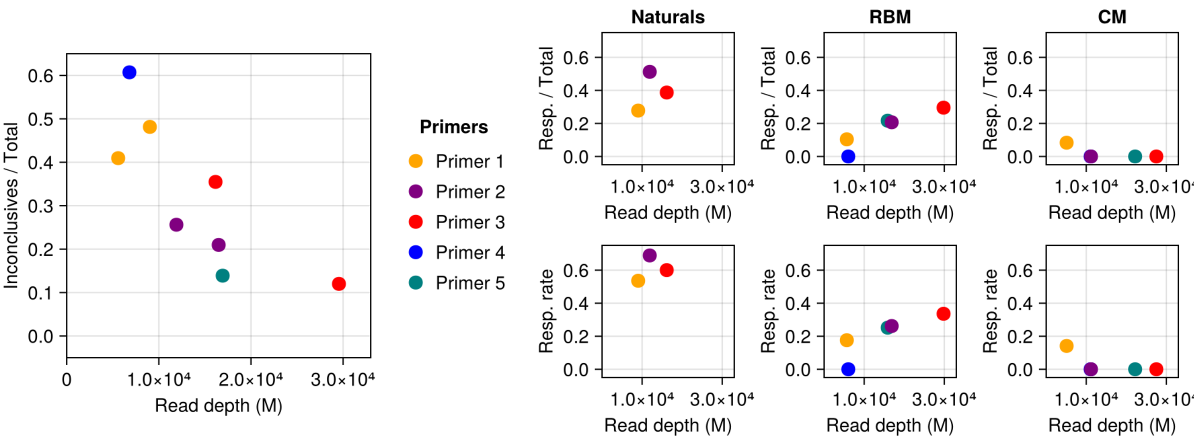


FIG. S32. Rates of response compared to read-depth in the Modified condition, for all the SHAPE experiments conducted in this work. The left panel shows the number of inconclusive aptamers divided by the total number of sequences for each group of aptamers that were sequenced together, colored by the primer used. As expected, the fraction of inconclusive aptamers decreases as the effective read depth increases (Pearson correlation −0.51). The right panels show the numbers of responsive aptamers among Naturals (left column), RBM (center column) and rCM (right column) as a function of the read depth. The top panels show the absolute numbers of responsive aptamers divided by the total numbers of aptamers probed. The bottom panels instead plot the response rate, defined as the ratio between responsive aptamers and total number of conclusive aptamers.

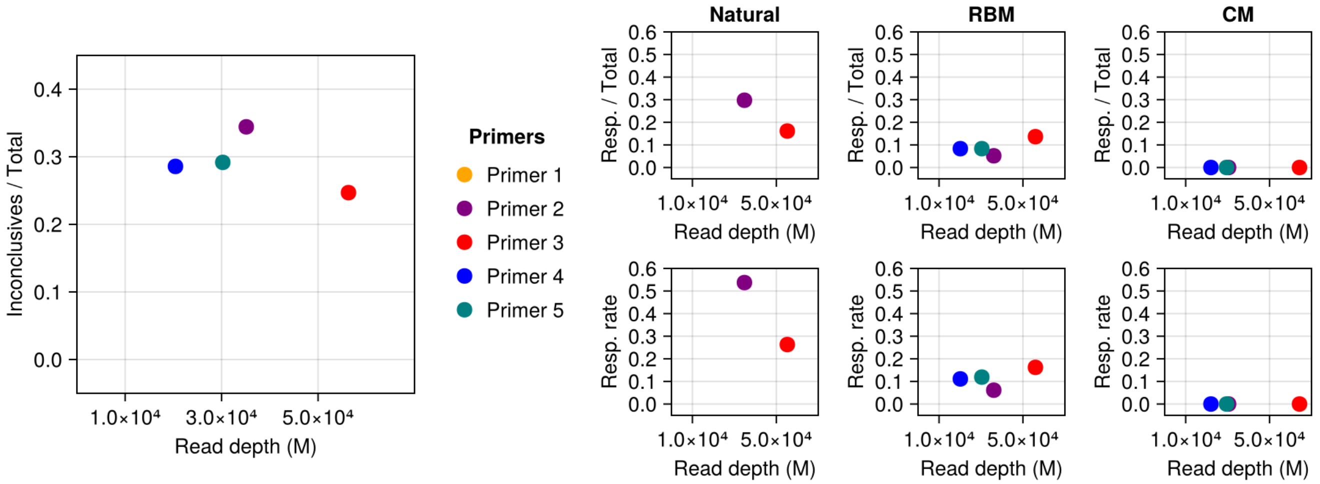


FIG. S33. Rates of response compared to read-depth in the Modified condition, for all the DMS experiments conducted in this work. Same as Figure S32 but for DMS.

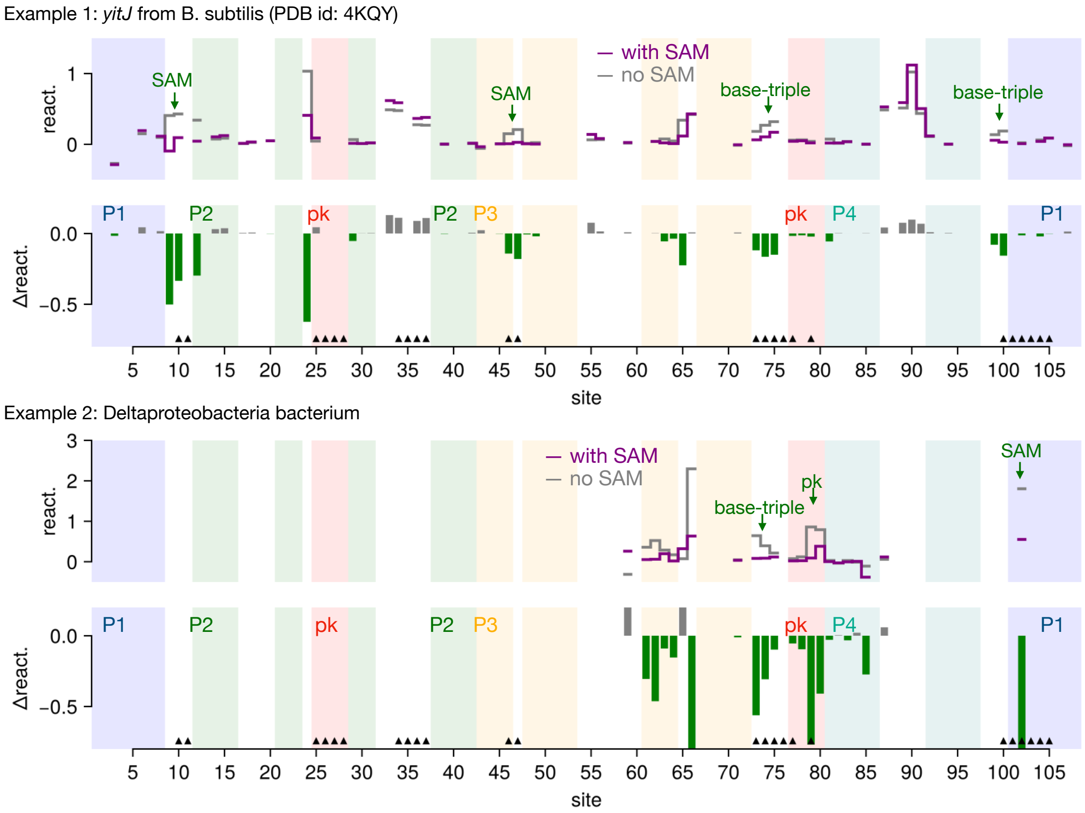


FIG. S34. DMS reactivity profiles for the two natural aptamers shown in the main text, Figure 5. Note that poor DMS coverage arises due to the fact that DMS reactivity probes only sites with A or C. See also Fig. S35C.

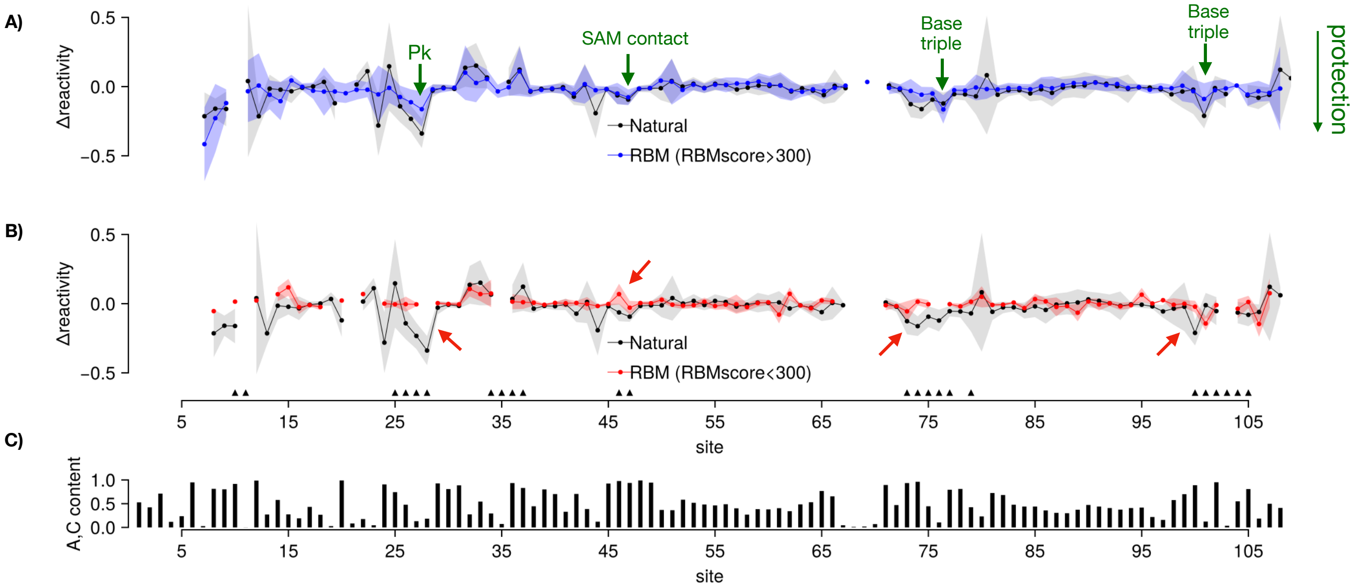


FIG. S35. Comparison of average DMS Delta-reactivities per site (with vs. without SAM) for natural sequences (gray), RBM generated sequences with RBMscore > 300 (blue) and RBM generated sequences with RBMscore < 300. Note that only sites with A or C are considered for DMS. The bottom panel shows the average A, C content in the set of aptamers probed with DMS.

### Appendix N: Some aptamers classified as non-responsive exhibit localized responses

Some of the RBM generated sequences classified as non-responsive, exhibit nonetheless localized reactivity changes in response to SAM, compatible with SAM binding, but not enough to conclude that a structural switch from the open to a closed conformation has taken place. Focusing on the first experiment, we find a total of 8 non-switcher RBM sequences exhibiting localized reactivity responses:

- 2 that respond at the SAM binding pocket
- 3 at the pseudoknot
- 3 at A minor
- 1 at base-triple

These are all different sequences. These results again highlight the diversity of structural responses displayed by the aptamer sequences tested in our experiments.

### Appendix O: Negative control for SHAPE detection of SAM response

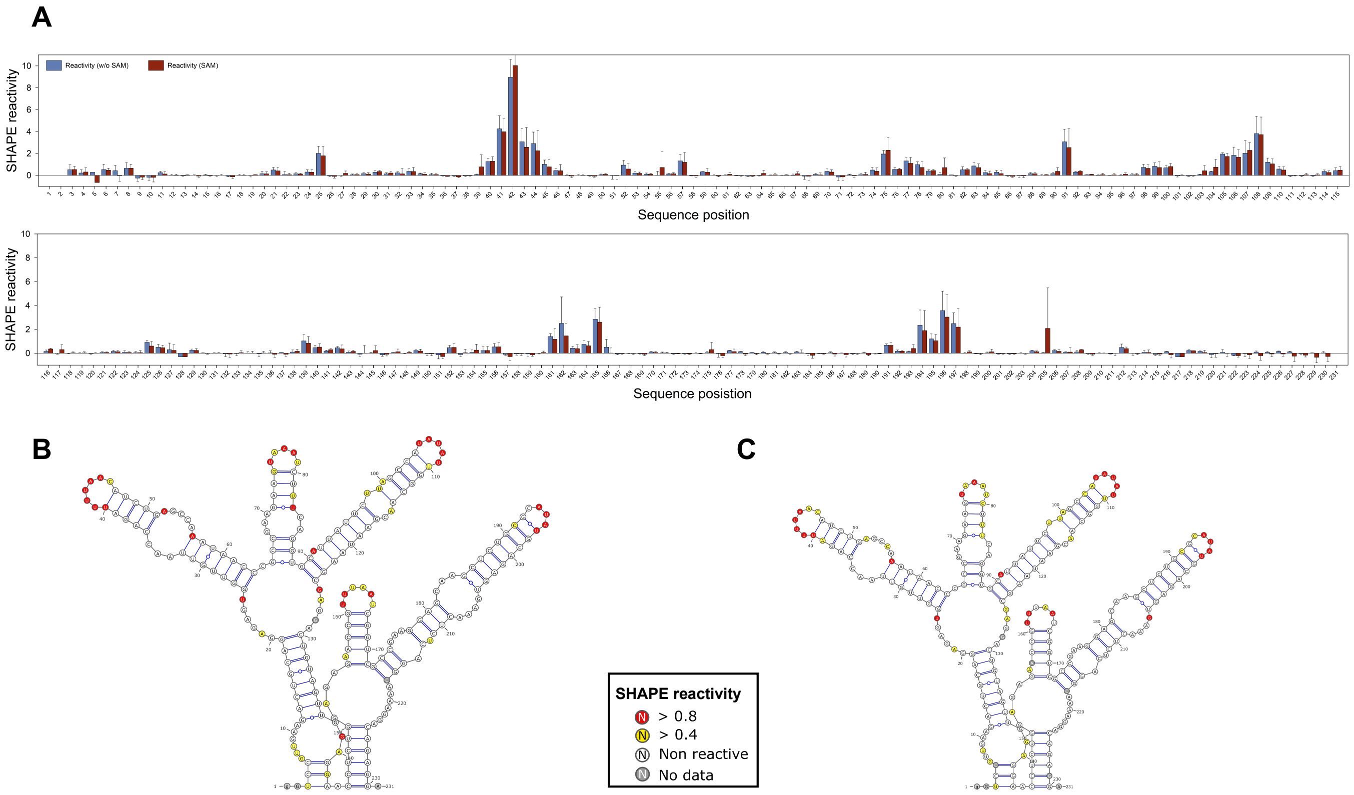


FIG. S36. Presence of SAM does not affect the SHAPE profile of a glycine riboswitch. **(A)** SHAPE reactivity profiles of the Glycine riboswitch from V.cholerae obtained in absence (blue) or presence (burgundy) of 1mM SAM. **(B)** and **(C)** Secondary structure scheme and SHAPE reactivity of nucleotides of the glycine riboswitch in absence (B) or presence (C) of 1mM SAM. Base pairs have been extracted from the RNA crystal structure (PDB id: 6WLT). Nucleotide reactivity are colour encoded as mentioned in the box

As a negative control of the ability of SHAPE reactivities to detect interactions with SAM, we probed a benchmark glycine riboswitch from V. Cholerae (PDB id: 6WLT) in presence and absence of SAM. This aptamer is not supposed to bind SAM, and therefore no reactivity changes are expected to occur when SAM is added. The results of this control experiment are shown in Fig. S36. The two reactivity profiles (SAM vs. no SAM) exhibit no significant differences. This confirms that alterations in the reactivity profile upon SAM addition are specific to aptamers that interact with SAM.

### Appendix P: Comparison of DMS and SHAPE probing results

As a further validation of our results, we performed complementary chemical probing with DMS of a subset of aptamers. In Figure S37 we compare here the results of the analysis of the data to the results obtained with SHAPE. In contrast to SHAPE, DMS probing is efficient in detecting interactions involving nucleotides A or C predominantly, but not G or U. DMS probing reactivities were analysed using the same methods. However to avoid to consider as non-switching residues with U or G, simply because the do not react, for a given aptamer we will remove from the hallmark sites a site carrying G or U nucleotides. As expected both in natural and artificial aptamers, we are able to detect less switching molecules. As can be appreciated in Fig. S37, DMS is more stringent in that it rarely detects a response to SAM in cases where SHAPE does not detect a response (only 2 cases in the 1st batch, 0 in the 2nd batch). The results show that the rate of success in switching behavior obtained by DMS is about one half of the one obtained from SHAPE probing both in natural and RBM designed molecules, and is kept to zero for the CM models in the first batch. We have investigated the dependence of DMS results on the read depth (Fig. S33). As for SHAPE the inconclusive rate for switching behavior increases by decreasing the read depth.

1st batch

**DMS+SHAPE** (columns) vs **SHAPE** (rows)

**Natural**

| SHAPE | Yes | No | Inc. |
|---|---|---|---|
| Yes | 72 | 2 | 6 |
| No | 1 | 32 | 2 |
| Inc. | 8 | 3 | 26 |

**RBM**

| SHAPE | Yes | No | Inc. |
|---|---|---|---|
| Yes | 13 | 1 | 0 |
| No | 1 | 44 | 0 |
| Inc. | 3 | 9 | 13 |

**RBM (RBMscore > 300)**

| SHAPE | Yes | No | Inc. |
|---|---|---|---|
| Yes | 13 | 1 | 0 |
| No | 1 | 25 | 0 |
| Inc. | 3 | 5 | 5 |

**rCM**

| SHAPE | Yes | No | Inc. |
|---|---|---|---|
| Yes | 0 | 0 | 0 |
| No | 0 | 15 | 0 |
| Inc. | 0 | 0 | 1 |

**DMS** (columns) vs **SHAPE** (rows)

**Natural**

| SHAPE | Yes | No | Inc. |
|---|---|---|---|
| Yes | 36 | 15 | 29 |
| No | 2 | 24 | 10 |
| Inc. | 3 | 6 | 27 |

**RBM**

| SHAPE | Yes | No | Inc. |
|---|---|---|---|
| Yes | 4 | 4 | 6 |
| No | 0 | 40 | 5 |
| Inc. | 3 | 9 | 13 |

**RBM (RBMscore > 300)**

| SHAPE | Yes | No | Inc. |
|---|---|---|---|
| Yes | 4 | 4 | 6 |
| No | 0 | 25 | 1 |
| Inc. | 3 | 5 | 5 |

**rCM**

| SHAPE | Yes | No | Inc. |
|---|---|---|---|
| Yes | 0 | 0 | 0 |
| No | 0 | 11 | 4 |
| Inc. | 0 | 0 | 1 |

2nd batch

**DMS+SHAPE** (columns) vs **SHAPE** (rows)

**RBM (RBMscore > 300)**

| SHAPE | Yes | No | Inc. |
|---|---|---|---|
| Yes | 18 | 4 | 4 |
| No | 0 | 57 | 3 |
| Inc. | 2 | 9 | 5 |

**CM**

| SHAPE | Yes | No | Inc. |
|---|---|---|---|
| Yes | 0 | 0 | 0 |
| No | 0 | 8 | 0 |
| Inc. | 0 | 2 | 0 |

**DMS** (columns) vs **SHAPE** (rows)

**RBM (RBMscore > 300)**

| SHAPE | Yes | No | Inc. |
|---|---|---|---|
| Yes | 9 | 9 | 8 |
| No | 0 | 56 | 4 |
| Inc. | 0 | 12 | 4 |

**CM**

| SHAPE | Yes | No | Inc. |
|---|---|---|---|
| Yes | 0 | 0 | 0 |
| No | 0 | 7 | 1 |
| Inc. | 0 | 2 | 0 |

FIG. S37. Experimental results with DMS chemical probing and comparison to SHAPE-MaP. The first row corresponds to DMS probing of a subset of aptamers from the first batch of experiments. We compare the results of the analysis using SHAPE data only to the analysis combining SHAPE and DMS data, as explained in the main text, see Eq. 11. The second row compares the results from SHAPE only to DMS data only. The last two rows perform the same comparisons for the subset of aptamers probed with DMS selected from the second batch. From left to right, the tables show the results for natural aptamers, RBM generated aptamers (all or those with RBMscore $>$ 300), and CM generated aptamers. Note that in the second batch, all aptamers were artificial and those generated by the RBM all had RBMscore $>$ 300.

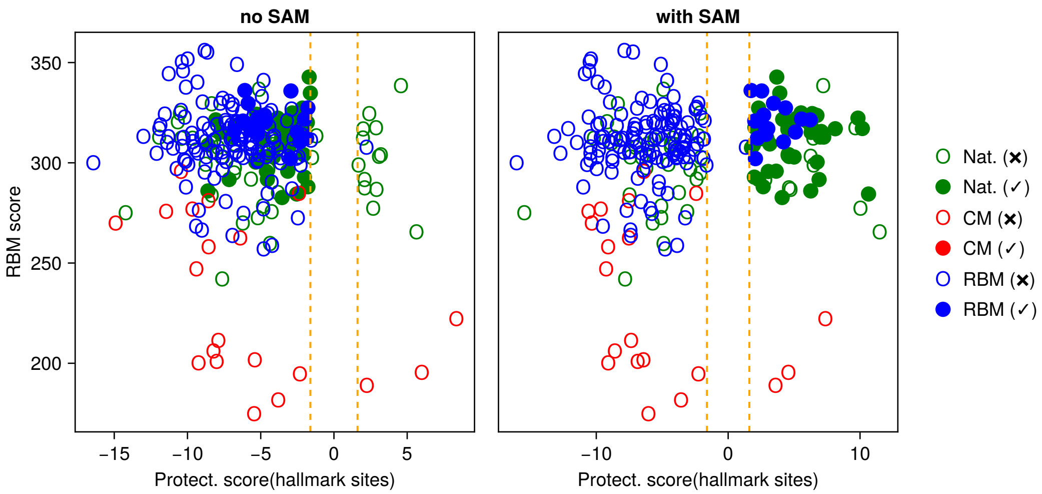


FIG. S38. DMS protection scores vs. RBM scores for all DMS probed aptamers. Panels: Left without SAM, Right with SAM. Responsive aptamers are shown with filled circles. Colors refer to the sequence origin: Natural, rCM, or RBM. Dashed orange vertical lines locate significance thresholds. Similar to Figure 7E & F in the main text, but for protection scores computed using the DMS probing data.

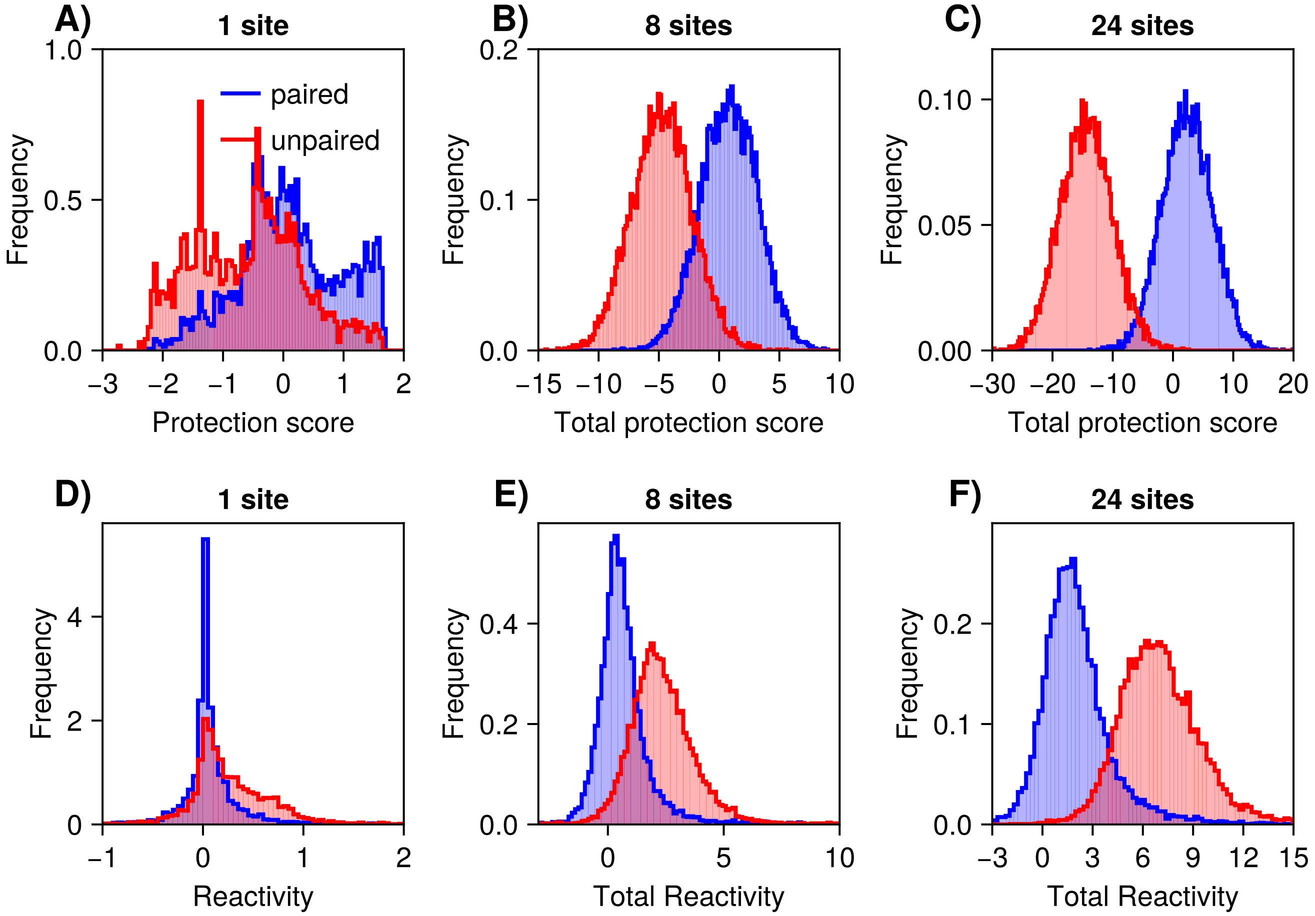


FIG. S39. Using DMS reactivity measurements from multiple sites leads to higher discriminative power. **A)** Empirical histogram of the protection scores of paired (blue) and unpaired (red) sites in the natural aptamers probed by DMS. **B)** Total protection score of 8 sites (same number involved in the pseudoknot of SAM-I riboswitches). **C)** Total protection score of 24 sites (same number involved in the Hallmark sites). **D)** Raw DMS-MaP reactivities of paired and unpaired sites. **E)** Total reactivities of 8 sites. **F)** Total reactivities of 8 sites. See also Fig. S25 for the counterpart of this figure for SHAPE data.

| Cluster | Sites |
|---|---|
| Pseudoknot | 25, 26, 27, 28, 77, 79 |
| Kink-turn | 34, 35, 36, 37 |
| Base-triple | 73, 74, 76, 100 |
| SAM contact | 10, 11, 46, 47, 102, 103 |
| P1 | 101, 104, 105 |
| Other sites | 75 |

TABLE S2. Hallmark sites of structural switch in response to SAM. List of sites that exhibit observable SHAPE reactivity differences upon SAM binding. Positions are numbered following the Rfam reference alignment. See also Fig. 10.

### Appendix Q: Hallmark sites

We selected a number of hallmark sites across the aptamer sequence, for which we could rationalize observed reactivity changes in response to SAM binding, and which are consistent with expectations from previous chemical probing studies on SAM-I riboswitches and previous structural data. They are listed in Table S2.

#### 1. Supporting literature

In this section, we discuss previous reports on the literature that support our selection of Hallmark sites.

*Pseudoknot.* The pseudoknot is a tertiary contact formed between sites 24-28 on loop L2 and sites 77-80, along the junction between P3 and P4. Multiple sources of evidence point to the importance of this motif to SAM binding, from genetic studies [21], to crystal structure [22, 23], to sequence based statistical modelling [24]. Consistently with these previous observations, we find in our experiments that sites 24-28, 77 and 79 in natural sequences exhibit significant reactivity decrease upon SAM-binding, as can be appreciated in Fig. 7 and 10D. We therefore include these sites in Table S2.

Sites 78, 80, also belonging to the pseudoknot, do not show significant protection upon SAM binding. As can be observed in the crystal structure (pdb 2GIS, [22]) site 80 at the edge of the pseudoknot is in a context and conformation favorable for the 1M7 probe to stack under the guanine and react with the cognate ribose, even when immobilized (E. Frezza, personal communication). This probably explains why site 80 remains slightly reactive even upon pseudoknot stabilization in the presence of the ligand. Site 78 is seen to exhibit protection in both conditions, likely due to other contacts formed outside the pseudoknot in absence of $Mg^{2+}$.

*Base-triple.* The A-minor motif helps create a groove where SAM is placed upon binding. It is an important ligand-dependent structural element [22, 23, 25]. While the two conserved G-C base pairs (21-30, 22-29) involved in this motif are stable and not reactive in any of the conditions assayed, our data clearly show consistent protections of A73 and 74.

The base-triple is an important tertiary motif observed in the bound structure of the aptamer [22, 23], involving nucleotides (24, 76 and 100) in between the A minor and the pseudoknot. Stabilization of this contact in response to SAM has been reported previously [22, 26]. Our data shows consistent protection of sites 76 and 100 in response to SAM, two of the sites involved in the base-triple.

*Kink-turn.* The kink-turn is a well characterized structural element of the SAM-riboswitch, with a central role in the fold of the aptamer, helping stabilize the coaxial stacking of the four helices and supporting the formation of the pseudoknot [25]. Stabilization of this tertiary structure in response to SAM has been observed, and evidenced both in the crystal structure [22, 23, 27], previous SHAPE experiments [26], and simulations [25]. Consistent with these previous observations, our data reveals significant protection at positions 34, 35, 36, 37 located in the kink-turn in response to both $Mg^{2+}$ and SAM. We therefore include these sites in Table S2.

*SAM contacts.* A number of sites are in direct contact with SAM in the bound structure, as has been established in published 3-dimensional structures [22]. SAM sits in a pocket between the interwoven P1 and P3 helices and the junction between P1 and P2, forming a network of contacts that results in stabilization of a number of sites. In particular sites 46, 47, belonging to a bulge in P3, directly contact SAM. Since these sites are initially unpaired, we expect a reactivity decrease upon SAM binding. Sites 102, 103 of the 5'-end arm of the P1 helix are similarly embracing SAM. Since even in the isolated aptamer domain, P1 might be disordered in absense of SAM (see Fig. 1A), we expect significant reactivity decreases in this region as well upon SAM binding. Also sites 10-11 show response in the averaged reactivity profiles (Fig. 7), but for many individual sequences we had few data at these sites and therefore were not included.

*P1 & other sites.* Stabilization of the P1 helix in response to SAM has a key regulatory role, releasing a complementary sequence that forms the hairpin loop and blocks downstream transcription, see Fig. 1B. We observe reactivity changes in sites 101, 104 and 105 of P1, likely reflecting the SAM induced stabilization of P1. These sites are also near SAM in the bound structure, though not in direct contact [22].

Finally, we observe significant protection in site 75 upon SAM binding. It is flanked by sites participating in the A-minor (74) and base-triple (76) motifs, both of which are significantly protected in response to SAM. The protection of neighboring sites is likely to promote low reactivity at 75.

### 2. Alternative definitions

We attempted several minor variations in the selection of Hallmark sites (see Table S2). We here report the results of some of these experiments, focusing on the first batch of probed aptamers (which includes natural sequences).

Figure S40 corresponds to Figure 7 in the main text, but reporting the results after minor variations in the definition of the Hallmark sites. First, we observe reactivity decreases in response to SAM in natural sequences at sites 24, 98, 99, which to the best of our knowledge had not been reported in the literature. Site 24 is involved in a base-triple and is neighbor to site 25 involved in the pseudoknot, which can explain its protection. Sites 98 and 99 are found in the junction J4/1. Panel A) of Fig. S40 reports the numbers of responding aptamers after including sites 24, 98, 99 Hallmark sites (in addition to Table S2). Then we considered sites 10, 11, of helix P1. Although these sites show reactivity responses to SAM, they are close to the beginning of the sequence and tend to have poorer reading depths than other more central nucleotides (see Fig. S27, S28, and following figures). This means that their SHAPE reactivities can be less reliable. Panel B) of Fig. S40 reports the numbers of responding aptamers after removing sites 10, 11 from the Hallmark sites.

In both cases, we see that the variations in the numbers of responsive or non-responsive aptamers are minor, $< 3\%$ in all cases. *C.f.* also results in Figure 7 of the main text. We conclude that our results are robust to minor variations in the definition of the set of Hallmark sites used in the main text.

**A)** Add sites 24, 98, 99.

| | Conclusive | Responsive | Non-responsive |
|---|---|---|---|
| Natural | 147 of 201 | 99 (67.3 ± 3.9%) | 48 (32.7 ± 3.9%) |
| Nat.(Seed) | 113 of 151 | 76 (67.3 ± 4.4%) | 37 (32.7 ± 4.4%) |
| Nat.(Hits) | 34 of 50 | 23 (67.6 ± 8.0%) | 11 (32.4 ± 8.0%) |
| Nat.(RBMscore>300) | 99 of 137 | 68 (68.7 ± 4.7%) | 31 (31.3 ± 4.7%) |
| Nat.(RBMscore>310) | 66 of 96 | 49 (74.2 ± 5.4%) | 17 (25.8 ± 5.4%) |
| Rfam CM | 15 of 16 | 0 (0%) | 15 (100%) |
| RBM | 59 of 84 | 14 (23.7 ± 5.5%) | 45 (76.3 ± 5.5%) |
| RBM(RBMscore>300) | 39 of 53 | 14 (35.9 ± 7.7%) | 25 (64.1 ± 7.7%) |
| RBM(RBMscore>310) | 30 of 40 | 12 (40.0 ± 8.9%) | 18 (60.0 ± 8.9%) |
| All | 221 of 301 | 113 (51.1 ± 3.4%) | 108 (48.9 ± 3.4%) |
| All(RBM score>300) | 138 of 190 | 82 (59.4 ± 4.2%) | 56 (40.6 ± 4.2%) |
| All(RBM score>310) | 96 of 136 | 61 (63.5 ± 4.9%) | 35 (36.5 ± 4.9%) |

**B)** Remove sites 10, 11.

| | Conclusive | Responsive | Non-responsive |
|---|---|---|---|
| Natural | 145 of 201 | 97 (66.9 ± 3.9%) | 48 (33.1 ± 3.9%) |
| Nat.(Seed) | 111 of 151 | 75 (67.6 ± 4.4%) | 36 (32.4 ± 4.4%) |
| Nat.(Hits) | 34 of 50 | 22 (64.7 ± 8.2%) | 12 (35.3 ± 8.2%) |
| Nat.(RBMscore>300) | 96 of 137 | 67 (69.8 ± 4.7%) | 29 (30.2 ± 4.7%) |
| Nat.(RBMscore>310) | 65 of 96 | 50 (76.9 ± 5.2%) | 15 (23.1 ± 5.2%) |
| Rfam CM | 14 of 16 | 0 (0%) | 14 (100%) |
| RBM | 59 of 84 | 14 (23.7 ± 5.5%) | 45 (76.3 ± 5.5%) |
| RBM(RBMscore>300) | 40 of 53 | 14 (35.0 ± 7.5%) | 26 (65.0 ± 7.5%) |
| RBM(RBMscore>310) | 31 of 40 | 12 (38.7 ± 8.7%) | 19 (61.3 ± 8.7%) |
| All | 218 of 301 | 111 (50.9 ± 3.4%) | 107 (49.1 ± 3.4%) |
| All(RBM score>300) | 136 of 190 | 81 (59.6 ± 4.2%) | 55 (40.4 ± 4.2%) |
| All(RBM score>310) | 96 of 136 | 62 (64.6 ± 4.9%) | 34 (35.4 ± 4.9%) |

FIG. S40. Same as Figure 7 in the main text, but with alternative definitions of the Hallmark sites. **A)** Adding hallmark sites 24, 98, 99. **B)** Removing hallmark sites 10, 11.

### Appendix R: Stability of P1 helix in the context of the full riboswitch sequence

To estimate the impact of the full riboswitch on the stability of the P1 helix, we have collected a total of 1306 full SAM-I riboswitch sequences (aptamer domain + expression platform) from the RiboD database [28]. For each of these full sequences, we estimated the matrix of base-pairing probabilities using ViennaRNA. We also estimated base-pairing probabilities in the corresponding aptamer domain sequences (removing the expression platform). The following Figures S41, S42, and S43 show the results.

First, in Figure S41 the different panels show the histograms of the base-pair probabilities for pairs of sites along the P1 helix. All pairs show a decrease of pairing probability in presence of the full riboswitch.

As negative controls, we considered P2 and P4 helices, with results shown in Figs. S42 and S43. For P2 and P4 the effect is much smaller or negligible.

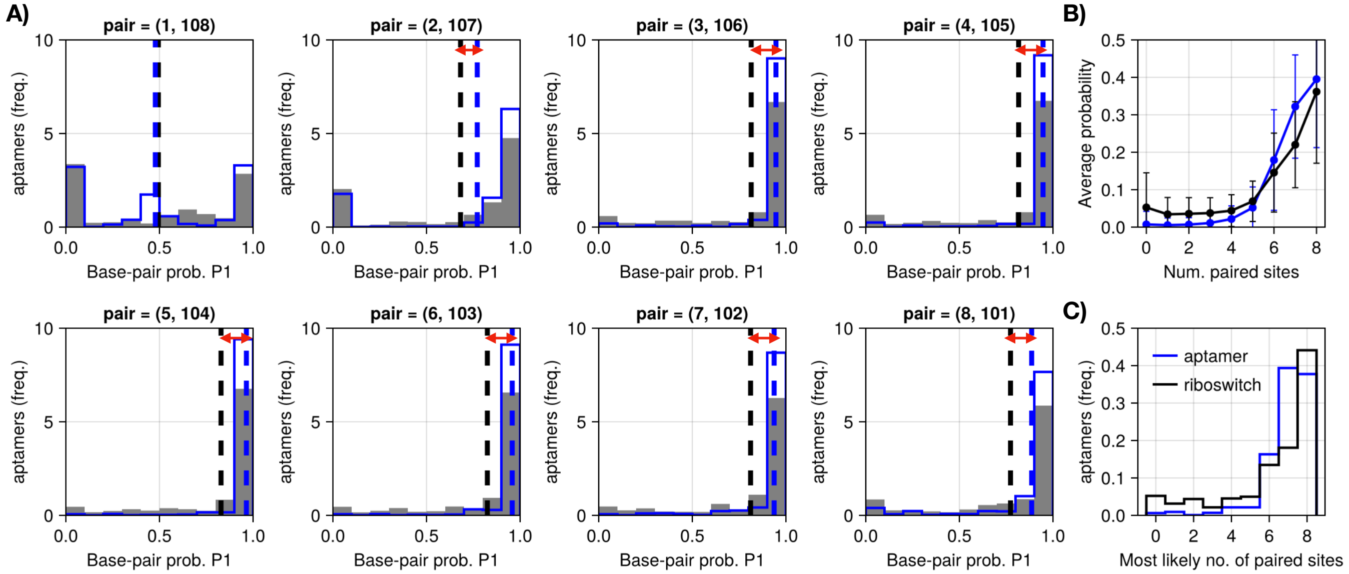


FIG. S41. Base-pairs along P1 are generally destabilized in the full riboswitch sequence in comparison to the aptamer only. **A)** The histograms show the base-pairing probabilities along the base-pairs in the P1 helix (predicted with ViennaRNA package), for the aptamer only (blue) and for full riboswitch sequences (gray). The average base-pairing probabilities are shown with dashed lines. **B)** To give another metric of the destabilization of P1, we work under the approximate simplifying assumption that base-pairs are independent. Then the number of base-pairs along P1 follows a Poisson-Binomial distribution. The plot shows the the average probability distribution of the number of base-pairs along P1 for different sequences. **C)** Under the same Poisson-Binomial approximation, we estimate the most likely number of base-paired sites for each sequence, and plot the resulting histogram.

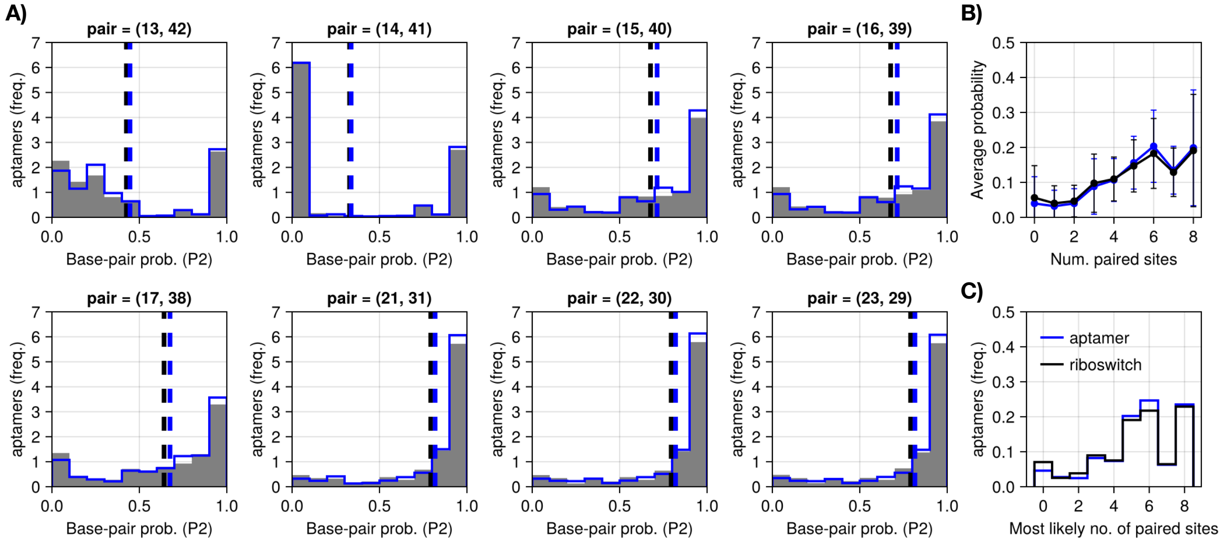


FIG. S42. Panels A), B) are similar to S41A,B, but for sites along the P2 helix. In contrast to P1, the P2 base-pairs are not destabilized by the presence of the full riboswitch sequence.

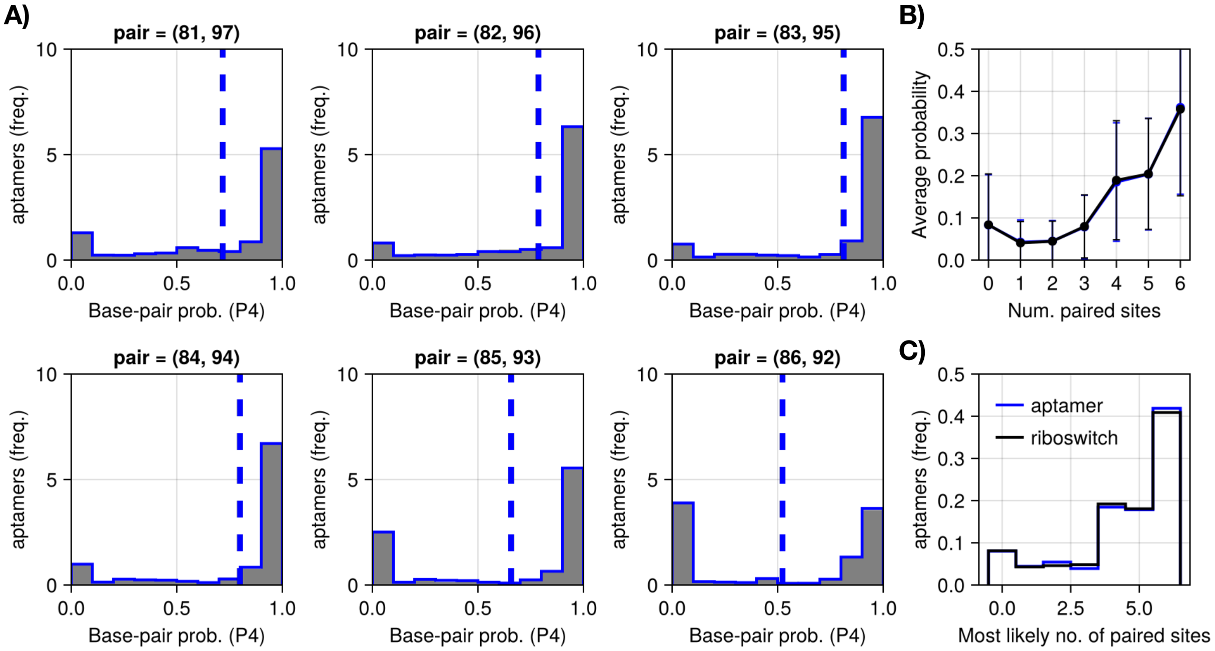


FIG. S43. Panels A), B) are similar to S41A,B, but for sites along the P4 helix. In contrast to P1, P4 base-pairs are not destabilized by the presence of the full riboswitch sequence.

### Appendix S: In vitro transcription protocol

RNA was *in vitro* transcribed by using T7 polymerase in 40 mM Tris-HCl pH 8.0, 25 mM MgCl2, 5 mM DTT, 5 mM NTPs and 20U of RNase inhibitor (Jena Bioscience®). Template DNA was then digested at 37°C for 30 min with DNAse I, RNase-free (ThermoFisher®) and RNA was precipitated during 30min by addition of 2.5M lithium chloride and centrifugation at 16,100xg for 30 min at 4°C. RNA pellet was washed with 70% ethanol then resuspended in nuclease-free water. RNA was purified through G50 size exclusion chromatography and quantified spectrometrically. Its integrity as well as the absence of aberrant products were confirmed by gel electrophoresis.
